# Efficient transgene-free genome editing in plants in the T0 generation based on a co-editing strategy

**DOI:** 10.1101/2023.03.02.530790

**Authors:** Xiaoen Huang, Hongge Jia, Jin Xu, Yuanchun Wang, Jiawen Wen, Nian Wang

## Abstract

Transgene-free genome editing of plants in the T0 generation is highly desirable but challenging, especially in perennials and vegetatively propagated plants. Here, we investigated the co-editing strategy for generating transgene-free, gene-edited plants via *Agrobacterium*-mediated transient expression of cytosine base editor (CBE)/gRNA-Cas12a/crRNA-GFP *in planta*. Specifically, CBE/gRNA was used to base edit the *ALS* gene to confer resistance to herbicide chlorsulfuron as a selection marker, which has no negative effects on plant phenotypes; Cas12a/crRNA was used for editing genes(s) of interest; GFP was used for selecting transgene-free transformants. Using this approach, transgene-free genome-edited plants were efficiently generated for various genes (either individual or multiplex) in tomato, tobacco, potato, and citrus in the T0 generation. The biallelic/homozygous transgene-free mutation rates for target genes among herbicide-resistant transformants ranged from 8% to 50%. Whole genome sequencing further confirmed transgene-free and absence of off-target mutations in the edited plants. The co-editing strategy is efficient for generating transgene-free, genome-edited plants in the T0 generation, thus being a potent tool for plant genetic improvement.

## Main

Transgene-free genome editing is highly desirable for plant genetic improvement. Cas9 and Cas12a DNA, mRNA or ribonucleoprotein complex (RNP) were successfully used to generate transgene-free plants ^1, 2^, which often require the transformation of embryogenic protoplasts. However, the regeneration of plants from protoplasts remains technically challenging and/or limited to specific plant species/genotypes ^3^. Until now, most genome-edited plants were generated through *Agrobacterium*-mediated transformation and are transgenic. Genome editing via transgenic approaches not only causes regulatory and public concerns ^4^, but also can generate new and off-target mutations in the next generation^5–7^. For annual crops such as rice, it is relatively easy to obtain transgene-free, gene-edited plants by genetic segregation via backcrossing or selfing ^8^. However, for perennials and vegetatively propagated plants, it is laborious and time-consuming to remove transgenes. Many crops lose traits of the parental cultivars via backcrossing, owing to their heterozygous nature as hybrids. Furthermore, in some plants, such as citrus and apple, the transgene cannot be removed through seed segregation once it is integrated into the plant genome because of their asexual reproduction nature through apomixis^9, 10^.

Even though genome editing via *Agrobacterium* results in transgenic plants, most T-DNAs used for carrying the Cas/gRNA do not integrate into the host chromosome, but are present in the nucleus, where they will be transcribed, leading to transient expression of the carried genes^11, 12^. The *Agrobacterium*-mediated transient expression was used for genome editing without transgene integration into plant genomes on several occasions ^13–16^. The main drawback of this approach identified in previous studies is that the majority of transformants are wild type, and most edited plants are mosaic/chimera or heterozygous and additional generations are needed to identify transgene-free and homozygous/biallelic mutants. In addition, previous genome editing through transient expression of Cas/gRNA constructs is usually performed without selection pressure, making it difficult, laborious and time-consuming to differentiate edited plants from unedited ones^17^.

In this study, we aimed to generate transgene-free genome-edited plants by employing T-DNA carrying CBE/gRNA-Cas12a/crRNA-GFP to co-edit the *ALS* gene, which encodes acetolactate synthase, and gene(s) of interest. Herbicides, such as chlorsulfuron, kills plants by acting as the inhibitors of acetolactate synthase. Mutation in the *ALS* genes using CBE confers resistance to herbicides such as chlorsulfuron in diverse plant species ^10, 15, 18–25^, thus providing a useful selection marker. The gene(s) of interest can be edited via Cas12a/crRNA, whereas GFP enables screening of putative transgene-free (GFP-negative) transformants. In this study, we have successfully used this co-editing strategy to efficiently generate transgene-free tomato, tobacco, potato, and citrus in the T0 generation for various genes. It is anticipated that this strategy will have broad applications in plant genetic improvements.

## Results

### Transgene-free genome editing of tomato in the first generation (T0) by co-editing of the *ALS* gene and gene of interest

To test whether we can achieve transgene-free genome editing in the T0 generation by co-editing of the *ALS* gene and gene of interest, we employed the model plant tomato (*Solanum lycopersicum*) owing to its high efficacy in plant transformation and genome editing^26^, availability of high-quality genome sequences^27^. We first investigated if we could obtain transgene-free, gene-edited tomato in the T0 generation by base-editing *SlALS1* (Solyc03g044330) alone. Previous studies suggested such a possibility, but the putative transgene-free plants were not confirmed by whole genome sequencing^10, 15, 28^. Here, we constructed the CBE-Cas12a-GFP-SlALS1 construct to edit the *SlALS1* gene using CBE to target the proline residue at position 186 (Pro186) (Fig. 1a). Our CBE-Cas12a-GFP construct also contains a GFP expression cassette for screening putative transgene-free regenerants, and the highly efficient, temperature-tolerant *ttLb*Cas12a ^29^ for editing gene of interest in downstream studies (Fig. 1a; Extended Data Figure S1). In accordance with the results reported by Veillet et al. ^15^, base editing of *SlALS1* enabled the generation of herbicide-resistant tomato transformants (Extended Data Figure S2). More than 20 chlorsulfuron-resistant lines without green fluorescence were obtained, suggesting putative transgene-free genome editing. Consistently, the *GFP* gene was not detected in 5 randomly selected GFP-negative lines with PCR (Fig. 1b). The targeted nucleotides of *SlALS1* gene by CBE were within the digestion site of the restriction enzyme *Sty*I (Fig. 1a), which enables identification of edited sequences by digestion. Editing of the targeted nucleotides completely abolished digestion by *Sty*I in one line, but partially abolished the digestion in four of the five tested lines (Fig. 1c), indicating homozygous/biallelic mutations in both alleles of *SlALS1* in line L2 and mutations in one allele only in the other four lines (L1, L3, L4, and L5). The mutations were confirmed by Sanger sequencing (Extended Data Figure S3). These results showed that editing Pro186 in either one or two *SlALS1* alleles enables herbicide resistance. To further confirm if the *SlALS1*-edited *GFP*-negative plants were indeed transgene-free, we conducted whole genome sequencing of the edited line 2 and confirmed the homozygous mutation of the *SlALS1* gene (Extended Data Figure S4). The sequence of the CBE-Cas12a-GFP-SlALS1 construct was not found in the genome of the *SlALS1*-edited, *GFP*-negative tomato plant. We also analyzed potential off-target genes. A total of 20 potential off-target sites with up to 4 mismatches to the target site were identified and none of them were edited (Extended Data Table S1), confirming the specificity of the base editing. These results show that we can obtain transgene-free, gene-edited plants in the first generation through base-editing of the *ALS* gene and selecting for chlorsulfuron-resistant and *GFP*-negative plants. This prompted us to further explore herbicide-assisted transgene-free genome editing in the T0 generation for gene(s) of interest.

**Fig. 1.**
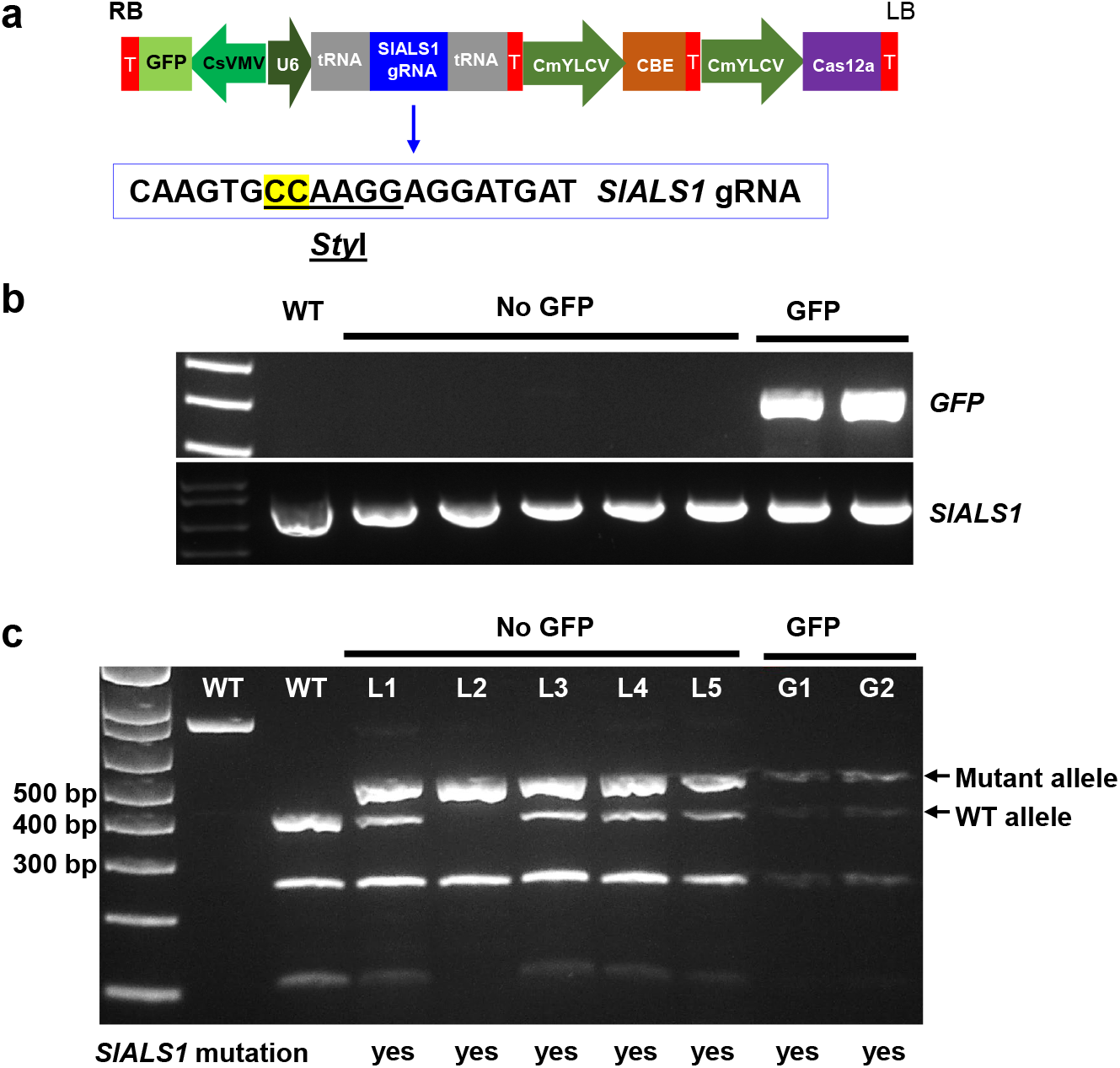
Establishment of herbicide-assisted transgene-free genome editing system. **a,** The CBE-Cas12a-GFP-SlALS1 construct used in the generation of transgene-free, *SlALS1*-edited tomato. The gRNA for *SlALS1* is boxed. The targeted nucleotides (CC) are highlighted in yellow. CsVMV, *Cassava vein mosaic virus* promoter; U6, citrus U6 promoter; CmYLCV, *Cestrum yellow leaf curling virus* promoter; CBE, cytosine base editor; T, terminator. For GFP, Nos terminator; for SlALS gRNA, poly (T) terminator; for CBE and Cas12a, HSP 18.2 terminator. RB, T-DNA right border; LB, T-DNA left border. **b,** PCR amplification of *GFP* in the regenerated chlorsulfuron-resistant tomato lines with or without green fluorescence. **c,** *SlALS1* gene genotyping of chlorsulfuron-resistant tomato regenerants through restriction enzyme digestion of PCR amplicons with *StyI*. PCR amplicons spanning the *SlALS1* gRNA region were subjected to restriction enzyme digestion with *StyI*. Editing of the targeted nucleotides abolishes the *Sty*I recognition site, resulting in resistance to *Sty*I digestion. Bottom text: *SlALS1* genotypes in the edited lines were confirmed by Sanger sequencing.

Co-editing of the *ALS* gene and gene(s) of interest has been suggested as a feasible approach to generate transgene-free plants^10, 15^. Next, we tested this hypothesis by co-editing *SlALS1* by CBE and *SlER* (Solyc08g061560) ^30^ by Cas12a in tomato using our CBE-Cas12a-GFP construct (Fig. 2a, Extended Data Figure S1). A total of 12 herbicide-resistant transformants without green fluorescence were selected for genotyping. Of the 12 herbicide-resistant transformants, the *SlALS1* gene was edited in all lines (Extended Data Figure S5). Five of these 12 herbicide-resistant plants were edited in *SlER*, with 3 being biallelic mutants and 2 being heterozygous mutants (Fig. 2b, Extended Data Figure S6). Consistent with the absence of green fluorescence, the 3 biallelic lines did not contain the *GFP* gene, as indicated by PCR, suggesting transgene-free (Fib. 2c). The 3 biallelic lines were further confirmed to be transgene-free by the absence of the *ttLbCas12a* sequence (Fib. 2c). The phenotypes of the *SlER* biallelic mutants included compact architecture, short petiole, densely clustered inflorescence, and enlarged SAM (Fig. 2d, e), in agreement with a previous report ^30^.To test the heritability of the mutation, seeds of the *sler-4* T0 plant were germinated, and the resulting seedlings were genotyped. The genotyping analysis found that mutation in the *SlER* gene was indeed heritable, with the progeny being either homozygous (inheriting one of two edited alleles) or biallelic (the same as their parent) (Extended Data Figure S7a). Additionally, PCR analysis confirmed that the seedlings did not contain *ttLbCas12a* or *GFP* (Extended Data Figure S7b), which was consistent with the absence of green fluorescence in the *sler-4* seeds (Extended Data Figure S8).

**Fig. 2.**
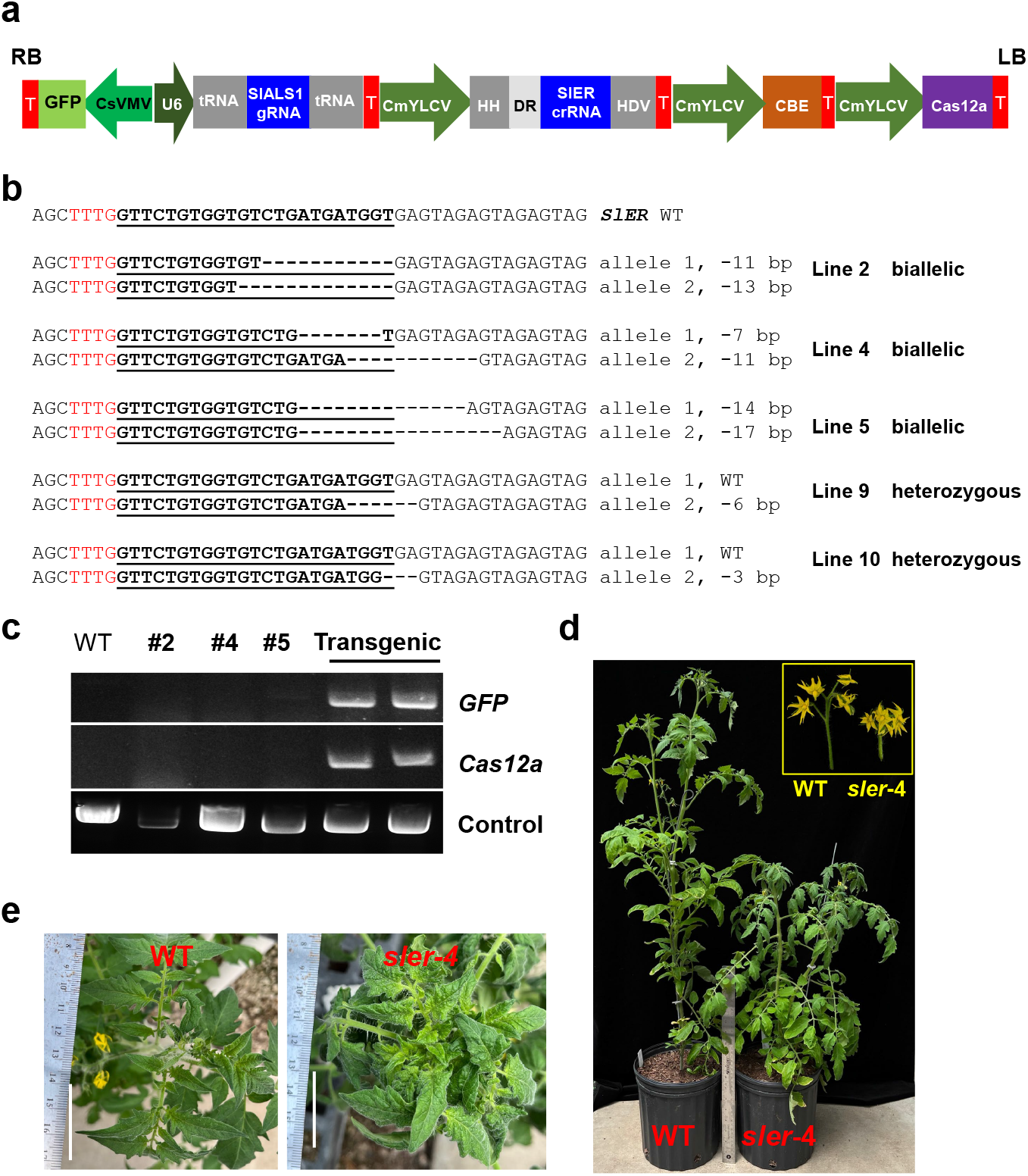
Transgene-free gene editing in the first generation (T0) in tomato. **a,** CBE-Cas12a-GFP-SlALS1-SlER construct used in the generation of transgene-free, *SlALS1*-edited, *SlER*-edited tomato. CsVMV, *Cassava vein mosaic virus* promoter; U6, U6 promoter; CmYLCV, *Cestrum yellow leaf curling virus* promoter; HH, ribozyme Hammerhead; DR, direct repeat; HDV, ribozyme hepatitis delta virus, CBE, cytosine base editor; T, terminator. For GFP, Nos terminator; for *SlALS* gRNA, poly (T) terminator; for *SlER* crRNA, poly (T) terminator followed by HSP 18.2 terminator; for CBE and Cas12a, HSP 18.2 terminator. RB, T-DNA right border; LB, T-DNA left border. **b,** The *SlER* genotypes of the edited lines without green fluorescence. **c,** PCR amplification of *GFP* and *Cas12a* from the biallelic mutants in b. **d, e,** Phenotypes of a representative transgene-free, *SlER*-edited line *sler-4*.

The co-editing strategy was also successful in generating transgene-free mutant lines for *SlRBL2* (Solyc09g010880) and *SlRbohD* (Solyc03g117980). We obtained 5 biallelic/homozygous mutants among 12 genotyped plants for *SlRBL2* (Extended Data Figure S9a). GFP observation and PCR analysis of the *GFP* gene and *Cas12a* gene (Extended Data Figure S9b) demonstrated that these 5 lines were transgene-free. For co-editing of *SlRbohD* with *SlALS1*, we tested one or two crRNAs (Fig. 3a, b). When one crRNA was used to target *SlRbohD*, only 1 biallelic mutant was generated (line 1) (Fig. 3c) among 12 non-GFP transformants. When two crRNAs targeting two different sites of *SlRbohD* were used (1 of these 2 crRNAs was the same as aforementioned), 25% biallelic/homozygous mutations were achieved (Fig. 3c, d, Extended Data Figure S10), suggesting that two crRNAs are more effective than one as reported previously ^31, 32^. GFP observation and PCR analysis of the *GFP* or *Cas12a* gene revealed that 3 lines, generated using either one or two crRNAs, were transgene-free (Fig. 3e).

**Fig. 3.**
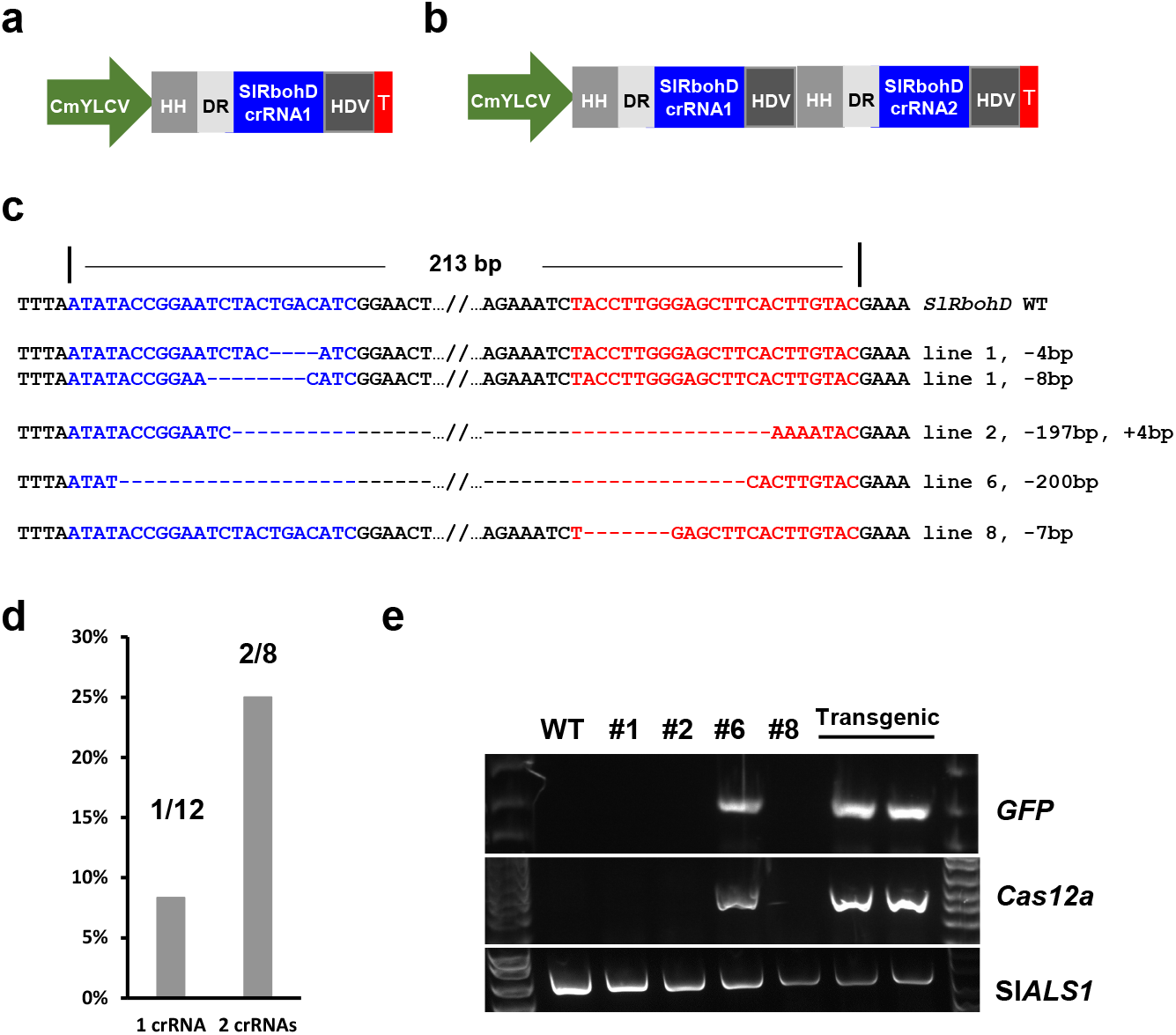
Efficient transgene-free gene editing of tomato in the T0 generation with 2 crRNAs. **a,** Construct scheme showing 1 crRNA targeting *SlRbohD*. Other parts of the construct are not shown. **b,** Construct scheme showing 2 crRNAs targeting *SlRbohD*. **c,** The *SlRbohD* genotypes of the transgene-free, homozygous/biallelic edited lines without green fluorescence. **d,** Comparison of rate of transgene-free, homozygous/biallelic mutants using 1 crRNA and 2 crRNAs. **e,** PCR amplification of *GFP* and *Cas12a* in the lines shown in c.

### Transgene-free, multiplex genome editing of tomato in the T0 generation

Next, we investigated whether we could achieve transgene-free, multiplex genome editing of tomato in the first generation. We performed co-editing of *SlEDS1* (Solyc06g071280) and *SlPAD4* (Solyc02g032850), with *SlALS1. EDS1* and *PAD4* are required for plant immunity^33–35^. Among 18 non-GFP regenerants, 6 lines contained biallelic/homozygous edits for both *SlEDS1* and *SlPAD4*, and 2 lines were biallelically edited in only *SlEDS1* but not *SlPAD4* (Fig. 4a, Supplementary Information File 1). The edited lines were transgene-free based on GFP observation and PCR analysis of the *GFP* and Cas*12a* genes (Fig. 4b). Similarly, we conducted multiplex gene editing of *SlDMR6* (Solyc03g080190) ^36^ and *SlINVINH1* (Solyc12g099200) with *SlALS1*. We obtained 3 biallelic/homozygous *SlDMR6/SlINVINH1* double mutants from 17 non-GFP transformants (Fig. 4c, Supplementary Information File 2). We also obtained 4 homozygous/biallelic *slinvinh1* single mutants (Supplementary Information File 2). These edited lines were transgene-free based on GFP observation and PCR analysis of the *GFP* and *Cas12a* genes (Fig. 4d). Taken together, we can achieve transgene-free, multiplex gene editing of tomato in the T0 generation efficiently.

**Fig. 4.**
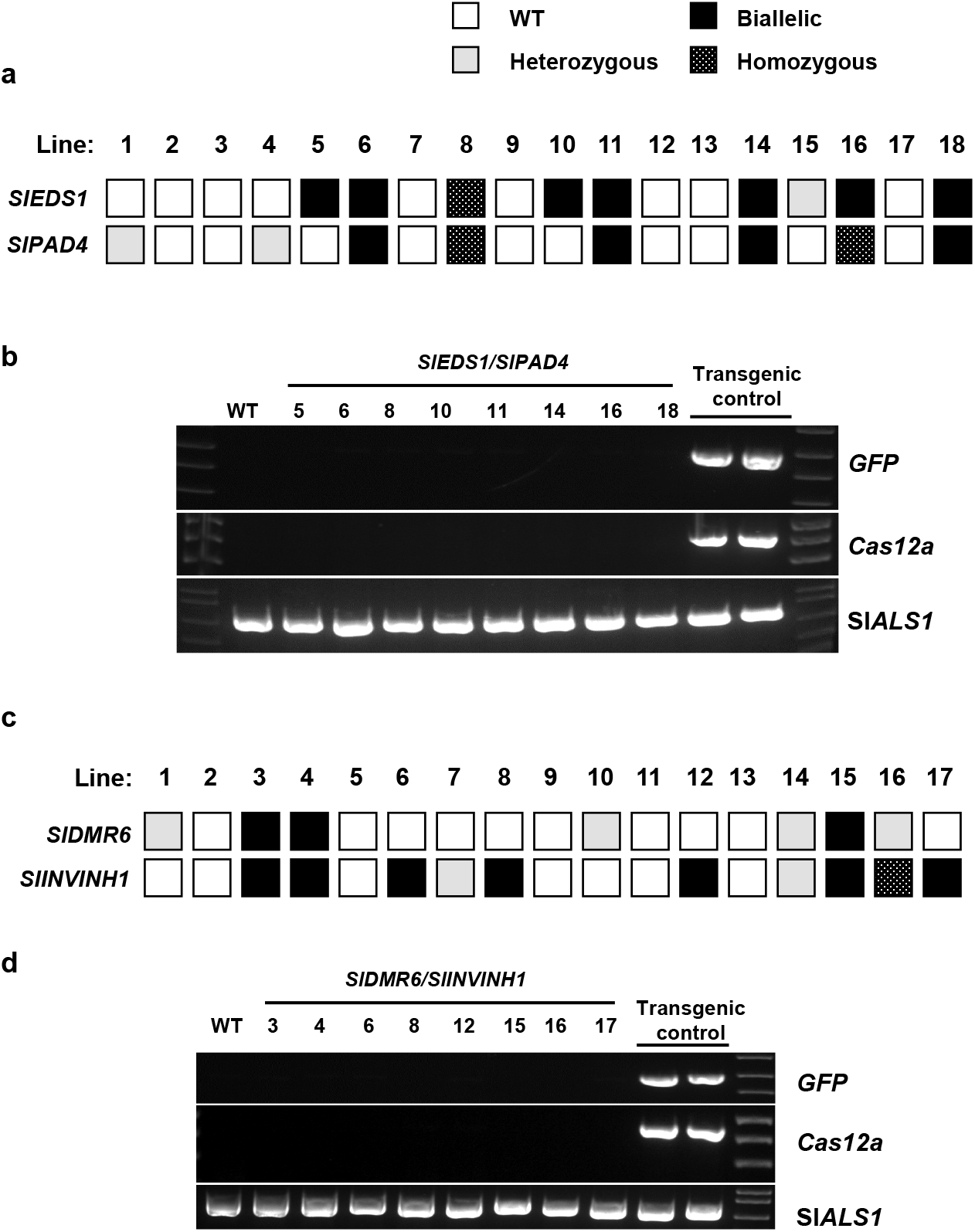
Transgene-free, multiplex gene editing of tomato in the first generation. **a,** Generation of transgene-free, biallelic/homozygous double mutants of tomato for *SlEDS1* and *SlPAD4*. **b,** PCR amplification of *GFP* and *Cas12a* from the edited *sleds1/slpad4* mutant lines from a. **c,** Generation of transgene-free, biallelic/homozygous double mutants for *SlDMR6* and *SlINVINH1*. **d,** PCR amplification of *GFP* and *Cas12a* from the edited *sldmr6/slinvinh1* mutant lines from c.

### Transgene-free genome editing of tobacco in the T0 generation

Next, we tested whether the co-editing strategy could be used to generate transgene-free plants in other plant species. We first investigated the model plant tobacco (*Nicotiana tabacum*) by co-editing *NtPDS* with *NtALS* (Fig. 5a). It is noteworthy that *N. tabacum* contains two *PDS* genes, *NtPDS1* and *NtPDS2*. Thus, we designed one crRNA targeting a conserved region of both genes. Over 20 chlorsulfuron-resistant tobacco plants showing albino phenotype were obtained (Fig. 5a). Among them, 7 albino plants did not display obvious green fluorescence (Fig. 5a). We further confirmed the absence of *GFP* and *Cas12a* in three albino, non-fluorescent lines by PCR (Fig. 5b, Extended Data Figure S11). As expected, the chlorsulfuron-resistant tobacco plant contained mutation in the *NtALS* gene, while the wild type *N. tabacum* did not contain the mutation in the *NtALS* gene (Fig. 5c, Extended Data Figure S12a). Genotyping of *NtPDS* in this line confirmed editing of both *NtPDS1* and *NtPDS2* genes, which was responsible for the albino phenotype (Fig. 5c, Extended Data Figure S12b).

**Fig.5:**
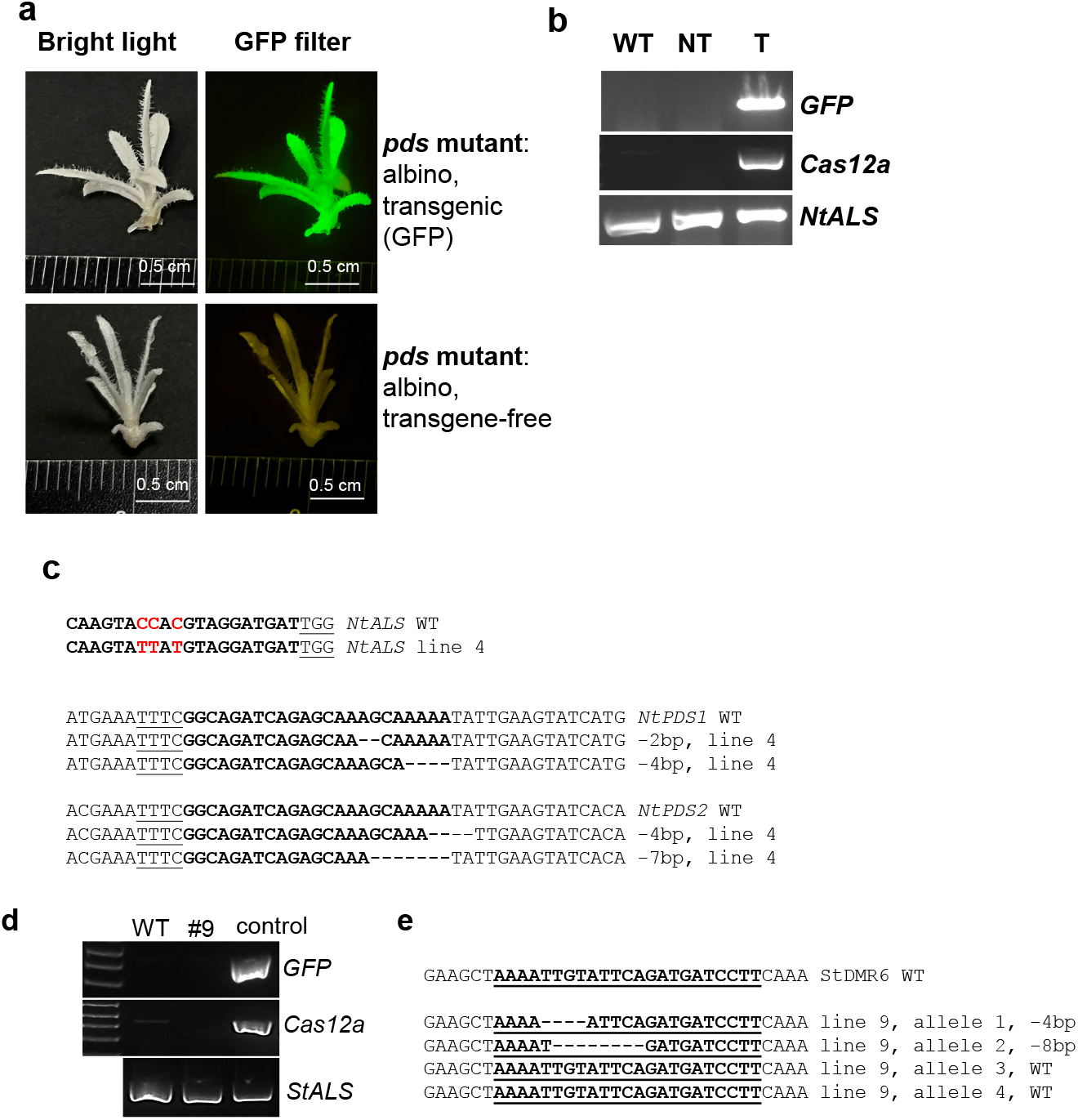
Transgene-free gene editing in the first generation (T0) in tobacco and potato. **a-c,** Co-editing of *NtALS* and *NtPDS* in *Nicotiana tabacum*. **a**, Albino phenotype with or without green fluorescence. Regenerants were selected on herbicide chlorsulfuron-containing media. Upper: transgenic albino tobacco plant; lower: transgene-free albino tobacco plant. **b,** Confirmation of transgene-free gene editing. PCR amplification of *GFP* and *Cas12a* in WT, non-transgenic (NT), and transgenic (T) plants. **c,** Genotypes of *NtALS, NtPDS* genes in a transgene-free, albino tobacco line from **a**. d & e, Transgene-free gene editing in potato**. d,** PCR amplification of *GFP* and *Cas12a* from a regenerated potato line 9 and control transgenic plant. **e,** Genotype of line 9 at *StDMR6*. crRNAs are underlined. 1 crRNA was used for *StDMR6* editing.

### Transgene-free genome editing of potato in the T0 generation

Furthermore, we investigated the feasibility of achieving transgene-free genome editing in potato, a vegetatively propagated crop with a tetraploid genome, using the co-editing strategy. We aimed to co-edit *StDMR6*, a disease susceptibility gene ^36, 37^, together with *StALS*, with a single crRNA that targets a conserved region in the first exon of four StDMR6 alleles. A total of 15 GFP-negative shoots regenerated from chlorsulfuron-containing media were genotyped and 10 were found to be wild-type (WT) at *StDMR6*, while 5 carried heterozygous edits. However, no tetra-allelic *StDMR6* mutants were observed, even in transgenic lines. The genotyping of a representative edited line revealed that it was transgene-free with 2 of the 4 *StDMR6* alleles edited (Fig. 5d, e, Extended Data Figure S13). These results suggest that our strategy can generate transgene-free, gene-edited potato in the T0 generation, but the generation of tetra-allelic mutants needs further optimization.

### Transgene-free genome editing of citrus in the T0 generation

Lastly, we aimed to achieve transgene-free genome editing of citrus in the T0 generation. Many tree plants, like citrus, have a long juvenile period, which makes it challenging to remove foreign DNA fragments when transgenic approaches are used for genome editing. We previously succeeded in obtaining transgene-free *ALS*-edited citrus through transient expression of CBE^10^. Here, we co-edited the TAL Effector Binding Element (EBE) region in the promoter of the citrus canker susceptibility gene *LOB1*^38–41^ with citrus *ALS* using our PBE-Cas12a-GFP-LOBP construct (Extended Data Figure S14). We utilized nCas9-PBE, a variant of CBE that has a unique 5 nucleotide editing window, resulting in targeting the proline residue at position 188 (Pro188) of *CsALS* only, which is equivalent to the proline residue at position 186 (Pro186) of *SlALS* ^42^. In the presence of chlorsulfuron, 107 pummelo (*Citrus maxima*) shoots were generated. Among them, 4 shoots were GFP-positive (Fig. 6a), and 103 shoots were GFP-negative (Fig. 6a). Based on genotyping of the *CsALS* and EBE_PthA4_-LOBP, and PCR analysis of the *nptII* gene (Fig. 6b), four transgene-free, EBE_PthA4_-LOBP-edited citrus lines were generated and subjected to downstream analyses. For the *CsALS* site, the four transgenic genome-edited and four transgene-free genome-edited lines contained homozygous/biallelic mutations (Extended Data Figures. S15 and S16). Intriguingly, among the four transgenic citrus genome-edited lines, for the EBE_PthA4_-LOBP site, Pum_GFP_1, Pum_GFP_2 and Pum_GFP_4 were chimeric, but without wild type sequences, and Pum_GFP_3 was wild type (Extended Data Fig. S15). Among the four transgene-free genome-edited lines, for the EBE_PthA4_-LOBP site, Pum_NoGFP_1, Pum_NoGFP_2, Pum_NoGFP_3 and Pum_NoGFP_4 contained biallelic, heterozygous, homozygous, and heterozygous mutations, respectively (Fig. 6c, Extended Data Fig. S16). As expected, biallelic/homozygous mutants and chimeric mutants without wild type sequence in the EBE_PthA4_-LOBP site demonstrated canker resistance and did not show any canker symptoms after inoculation with *Xcc*, regardless of being transgenic or transgene-free (Fig. 6d). Wild type pummelo showed typical canker symptoms, such as hyperplasia and hypertrophy (Fig. 6d). As a positive control, we inoculated wild type and genome-edited lines with XccΔpthA4:dLOB1.5. dLOB1.5 is a designed TALE, which binds to a different region from the target EBE_PthA4_-LOBP site in the promoter region of *CsLOB1*, thus activating *LOB1* expression to cause canker symptoms ^41^. Sanger sequencing results indicated that the dLOB1.5 binding sites were intact among the tested Pummelo plants (Extended Data Fig. S17). Consequently, XccΔpthA4:dLOB1.5 caused typical canker symptoms in wild type and all EBE_pthA4_-LOBP-edited lines (Fig. 6d). Taken together, the mutation of EBE_pthA4_-LOBP conferred Pummelo canker resistance, consistent with previous studies ^39, 41, 43^. Importantly, two transgene-free plants, Pum_NoGFP_1 and Pum_NoGFP_3, were resistant to *Xcc* infection (Fig. 6).

**Fig. 6:**
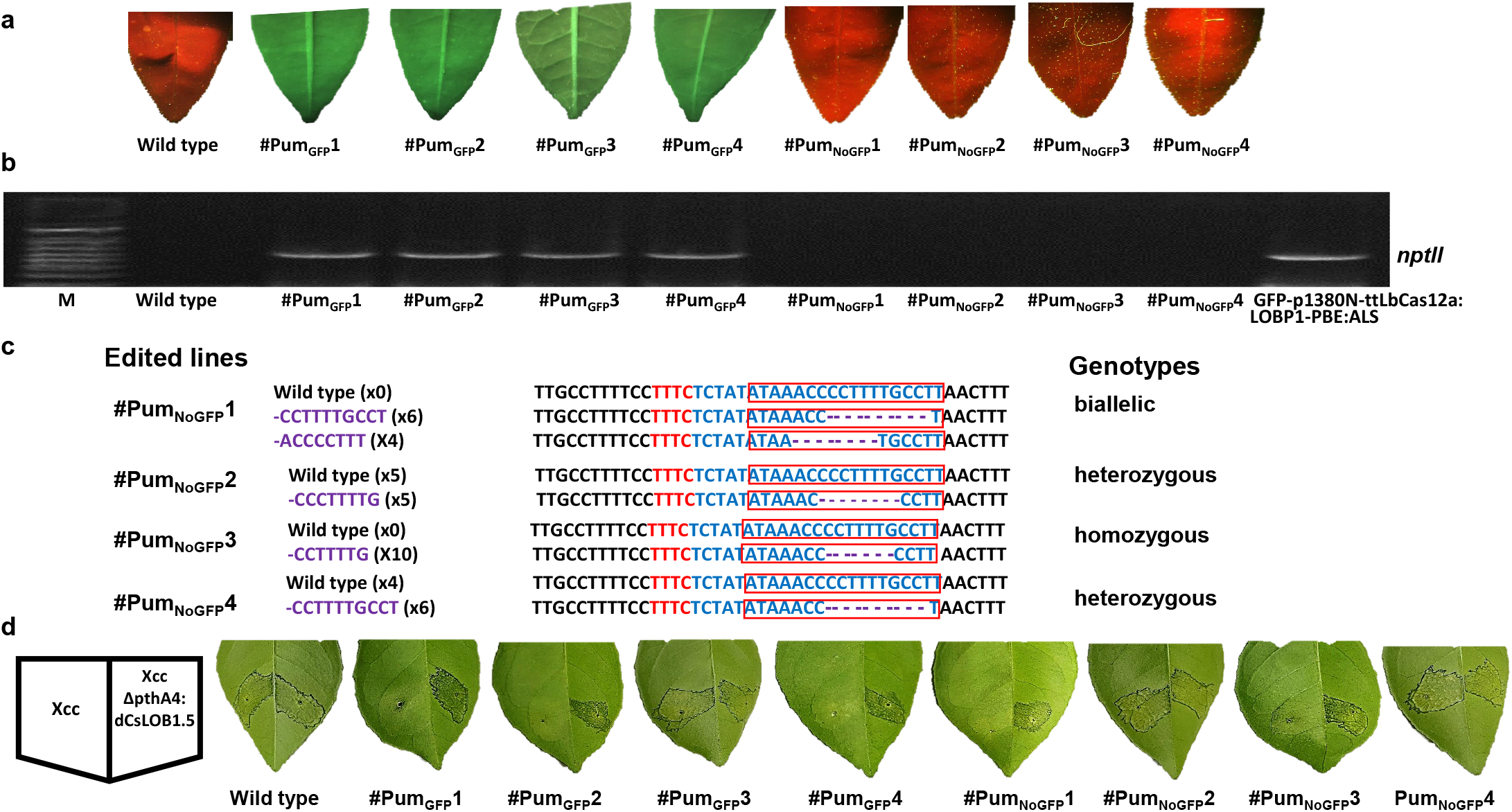
Transgene-free gene editing in the first generation (T0) in pummelo (*Citrus maxima*). **a,** GFP fluorescence was observed in transgenic Pummelo plants, whereas wild type and transgene-free plants did not exhibit any GFP signal. **b**, Using a pair of primers Npt-Seq-5 and 35T-3PCR, wild type, transgenic and transgene-free Pummelo plants were analyzed. The wild type Pummelo and plasmid GFP-p1380N-ttLbCas12a:LOBP1-EBE:ALS were used as controls. M, 1kb DNA ladder. **c**, Sanger sequencing analysis of GFP-negative lines by PCR amplification and cloning of *LOB1* promoter. **d**, Canker-resistance in the transgenic and transgene-free Pummelo plants. Five days post Xanthomonas citri subsp. citri (Xcc) inoculation, citrus canker symptoms were observed on wild type Pummelo, transgenic Pum_GFP_3, transgene-free Pum_NoGFP_1 and Pum_NoG_F_P_3, whereas no canker symptoms were observed on other LOBP-edited Pummelo plants, which could be attributed to 100% mutation rates in Pum_GFP_1, Pum_GFP_2, Pum_GFP_4, Pum_N_oGFP2 and Pum_N_oGFP4. As expected, *XccpthA4:Tn5(dCsLOB15)* caused canker symptoms on all plants. dCsLOB1.5 induces *LOB1* to cause canker symptoms by recognizing a different region from EBE_PthA4_-TII LOBP. GFP-positive lines: Pum_GFP_1 to Pum_GFP_4. GFP-negative lines: Pum_NoGFP_1 to Pum_NoGFP_4.

### Whole genome sequencing analysis of edited lines confirms transgene-free genome editing without off-target mutations

To further confirm whether the putative transgene-free genome-edited lines were indeed transgene-free, we conducted whole genome sequencing. For tomato, we sequenced six transgene-free lines that were edited for *SlRbohD* (#1, #2, #8), *SlER* (#4), *SlEDS1/SlPAD4* (#8, #18), as well as two transgenic control lines, EPGFP and *SlRbohD* (#6) (Extended Data Table S2). The sequencing coverage ranged from 28 × to 59 ×. Genomic analysis confirmed that the construct DNA was integrated into the genome of the transgenic control lines EPGFP and *SlRbohD* (#6) (Extended Data Table S2), as evidenced by the presence of construct reads (Extended Data Figure S18, Extended Data Table S2). In contrast, genomic analysis of the *SlRbohD* (#1, #2, #8), *SlER* (#4), *SlEDS1/SlPAD4* (#18) lines found no construct DNA (Extended Data Table S2). Intriguingly, *SlEDS1/SlPAD4* line 8 contained 281 reads matching construct sequences despite being GFP-negative and PCR-negative for GFP and Cas12a (Fig. 4b). Genomic analysis of the edited lines confirmed genome editing, as demonstrated by Sanger sequencing results. For instance, *SlRbohD* edited line #1 contained biallelic mutations of −4/-8, whereas #8 contained homozygous mutations of 7 bp deletion at *SlRbohD* (Extended Data Figures. S19-S20), consistent with previous Sanger sequencing results (Fig. 3c, Extended Data Figure S10). We searched for potential off-target sites of the crRNA targeting *SlRbohD, SlER, SlEDS1, SlPAD4, SlINVINH1*, and *SlRBL2* genes using the CRISPR P v2.0 program and Cas-OFFinder program. Whole genome sequencing analyses or Sanger sequencing of PCR amplicons of the homologous sites showed no off-target mutations (Extended Data Table S3). Similarly, whole genome sequencing analysis of the GFP-negative, *LOB1*-edited citrus line Pum_NoGFP_3 (Extended Data Table S2) found no construct DNA in its genome. Furthermore, whole genome sequencing analysis indicated that Pum_NoGFP_3 harbored heterozygous *CsALS* and homozygous mutant EBE_PthA4_-LOBP, which was consistent with Sanger sequencing results (Fig. 6c). In addition, off-targets were analyzed in Pum_NoGFP_3 based on whole genome sequencing data. In the case of mismatch number<=5 for crRNA, there were eight potential off-targets (Extended Data Figure. S21). The off-target sites were visualized using IGV software version 2.15.4 (Robinson et al., 2011), and no off-target mutations were identified.

## Discussion

In this study, we have developed a potent genome editing toolkit to generate transgene-free genome-edited plants in the T0 generation by co-editing the *ALS* gene and gene(s) of interest through *Agrobacterium*-mediated transient expression. It was successfully used for genome editing of tomato and tobacco (annuals), citrus (a perennial tree crop), and potato (a vegetatively propagated tetraploid crop). The biallelic/homozygous mutation rates for target genes among herbicide-resistant transformants in our study ranged from 8% to 50%, which is comparable to the genome editing efficacy using Cas/gRNA DNA, mRNA, or ribonucleoproteins ‘. The efficient identification of biallelic/homozygous mutants resulted from co-editing of the *ALS* gene, which provides a useful and practical selection marker as a gain-of-function against sulfonylurea herbicides. Importantly, precise editing by CBE targets the proline residue, for instance, Pro186 in tomato or Pro188 in citrus, which probably disrupts the recognition and binding of the herbicides without affecting ALS function ^44^. Consistently, *ALS* mutants at the proline residue did not show any phenotypical changes in our study. Natural mutations of the *ALS* gene are prevalent in plant species without affecting plant phenotypes (except herbicide resistance) and fitness ^45–49^. Thus, editing of the *ALS* gene as a selection marker will not negatively affect genetic improvement of crops and their commercialization.

The co-editing strategy has multiple advantages in generating transgene-free genome-edited plants: 1) The co-editing strategy ensures transgene-free genome editing by transient expression of Cas/gRNA and removing of stable transformants using GFP as an indicator. Transiently expressed Cas/gRNA eventually degrades, thus mimicking the approaches that use Cas/gRNA DNA in transgene-free genome editing via transformation of protoplasts^50^. Genotyping of construct components such as GPF, Cas12a, or *nptII* genes and whole genome sequencing of putative transgene-free lines indeed confirm the absence of foreign genes in 87.5% (7 of 8) putative transgene-free lines. Intriguingly, the *sleds1/slpad4* line 8 contained 281 reads matching construct sequences whereas the two transgenic lines contained 1830 and 2716 construct reads. This is consistent with previous reports that small DNA fragments from transformed plasmids are sometimes inserted at both on-target and off-target sites in host cells, even though at low frequencies ^51^. Consequently, whole genome sequencing is required to verify the edited lines to be transgene-free. “Transgene-free” is a prerequisite for commercialization of genetically modified organisms. Transgenic crops are under robust and strict regulations in different countries and regions, ^4^ and cause negative public reception, which impedes their commercialization despite superior traits. 2) Transgene-free lines without T-DNA eliminate the potential disruption of gene functions at the insertion site caused by T-DNA in *Agrobacterium*-mediated stable transformation ^52^. Thus, this approach is not only suitable for crop genetic improvements, but also provides advantages in genetic studies. 3) Off-target mutations are another critical factor for consideration during genetic improvement by genome editing. No off-target mutations were identified in our genome-edited lines. This probably results from the short functional time of Cas/gRNA during transient expression, as suggested by previous studies ^10, 53^. Similarly, transient expression of Cas/gRNA DNA, mRNA, and RNP in embryogenic protoplasts, calli, or immature embryo cells has been reported to generate transgene-free plants without causing off-target mutations^1, 2, 54^. 4) This co-editing strategy can produce transgene-free, gene-edited plants in the T0 generation. Generation of transgene-free genome-edited plants in the T0 generation bypasses the need to remove transgenes in future generations by genetic separation via backcrossing or selfing. The removal of transgenes is not feasible for many crops which are asexually propagated, or highly heterozygous, or have long juvenility, such as grape, citrus, potato and banana. Generation of transgene-free plants in the T0 generation significantly expedites the genetic improvement of crops. For example, transgene-free citrus was generated within 6 months using our approach. However, it takes approximately 20 years to generate new citrus varieties using traditional breeding approach ^55^. Lastly, our strategy is based on *Agrobacterium*-mediated delivery of CRISPR components into recipient plant tissues such as cotyledons, leaves, and epicotyls. Hence, it can be easily adopted because *Agrobacterium-mediated* transformation is one of the most widely used and convenient methods. Transformation of embryogenic protoplasts with Cas/gRNA RNP^56^, or DNA^50^ has successfully generated transgene-free, genome-edited plants. However, plant regeneration from protoplasts is technically challenging. Noticeably, the regenerated plants from protoplasts are prone to somaclonal variations and genome instability^57, 58^. In addition, regeneration from protoplasts is not accessible to many plant species, especially monocots. Another method to generate transgene-free edits is to transiently deliver plasmids, mRNA, or RNPs directly into callus cells or immature embryos through biolistic particle bombardment^3, 59, 60^. However, due to low efficiency and no selection, a huge amount of work must be done on tissue culture, regeneration, genotyping, and selection of edited plants from unedited plants. It is not surprising that the co-editing strategy may also aid in the enrichment and selection of edited protoplasts, callus cells or immature embryos achieved through means other than *Agrobacterium*.

In sum, we have developed an efficient transgene-free genome editing methodology based on *Agrobacterium*-mediated transient expression for plants. As *Agrobacterium*-mediated transformation works for many plant species, we anticipate that this approach has broad applications in genetic improvements and genetic studies of plants. It is particularly useful for perennials and vegetatively propagated plants to generate transgene-free, gene-edited plants in the T0 generation.

## Methods

### Making the CBE-Cas12a-GFP construct

The CBE plasmid with GFP ^10^ was digested with *Pme*I/*Rsr*II and the vector backbone was retained. The Cas12a-D156R^29^ fragment was recovered by digesting Hybrid-D156R-PDS-LOB1-A containing tt*Lb*Cas12a^29^ with *Pme*I/*Rsr*II. These two fragments were then ligated to form CBE-Cas12a-partial. CBE-Cas12a-partial was then digested with *Rsr*II. The other half of Cas12a was PCR-amplified using primers Cas12half-F1/Cas12half-R1 (Extended Data Table S4) with tt*Lb*Cas12a^29^ as the template. The amplicon was then In-fusion cloned with the *Rsr*II-digested CBE-Cas12a-partial to create CBE-Cas12a-GFP. The final construct was confirmed through Sanger sequencing.

### Making CBE-gRNA-Cas12a-crRNA-GFP constructs

CBE-Cas12a-GFP was digested with *Aar*I and ligated with annealed primers for *NtALS* or *CsNLS* or *SlALS1* or *StALS* (Extended Data Table S4) with compatible ends. A construct named PUC57-mini-crRNA was synthesized to drive crRNA expression (Extended Data Fig. S22). The crRNA array is flanked by ribozymes, Hammerhead (HH) and hepatitis delta virus (HDV), for precise processing^61^. Primers for single crRNA were annealed and ligated to *Bsm*BI-digested PUC57-mini-crRNA. For multiplexing, multiple HH-DR-HDV units were PCR amplified from the synthesized plasmid PUC57-HDV-HH-DR (Extended Data Fig. S22). The PCR products were ligated and cloned into *Bsm*BI-digested PUC57-mini-crRNA through GoldenGate cloning. The whole crRNA cassette, including the promoter and terminator, was PCR amplified using primers Mini-F1/Mini-R1 and cloned into the *Sbf*I-digested CBE-Cas12a-GFP-ALS constructs using In-Fusion cloning (Takara Bio). All constructs were confirmed by sequencing.

### Making the GFP-p1380N-ttLbCas12a:LOBP1-PBE:ALS construct

Using pUC-NosT-crRNA:LOBP as template^43^, the fragment containing AtU6-26 promoter, the coding sequence of hammerhead ribozyme (HH) and crRANA-LOBP1 was PCR-amplified using primers AtU6-5-*Xho*I (5’-AGGT<u>CTCGAG</u>TCGTTGAACAACGGAAACTCGA CTTGCC-3’) and CrRNA-LBDP1-phos (5’-phosphorylated-AAGGCAAAAGGGGTTTATAT AGAATCTACACTTAGTAGAAATTAga −3’), and the fragment containing the coding sequence of hepatitis delta virus ribozyme (HDV) and NosT was PCR-amplified using primers HDV-5-Phos (5’-phosphorylated-GGCCGGCATGGTCCCAGCCTCCTCGCT - 3’) and NosT-3-*Asc*I (5’-ACCTGGGCCC<u>GGCGCGCC</u>GATCTAGTAACATAGATGA-3’). *Xho*I-cut AtU6-26-HH-crRNA-LOBP1 and *Asc*I-cut HDV-NosT were inserted into *Xho*I-*Asc*I-cut pUC-NosT-MCS to build pUC-NosT-crRNA:LOBP1 through three-way ligation, in which the vector and two DNA fragments were ligated together in one step. pUC-NosT-MCS contained *Eco*RI-NosT-*Xho*I-*Asc*I-*Xba*I-*Pme*I for cloning, as described before ^62^. Subsequently, the *Eco*RI-NosT-crRNA:LOBP1-NosT-*Asc*I-*Xba*I-*Pme*I fragment was cloned into *Eco*RI-*Pme*I-cut GFP-p1380N-ttLbCas12a to generate GFP-p1380N-ttLbCas12a:LOBP1-*Asc*I-*Xba*I-*Pme*I (Extended Data Figure S14). GFP-p1380N-ttLbCas12a was constructed previously ^63^.

From vector CmYLCV-A3A-RAD51-nCas9 ^10^, the CmYLCV promoter was amplified using primer CmYLCV-5-*Hin*dIII*-Sb/*I-*Asc*I (5 ‘-AGGT<u>AAGCTTCCTGCAGGCGCGCC</u>AGATTTGCCTTTTCAATTTCAGAAAGA-3’) and CmYLCV-3-*Bam*HI (5’-AGGT<u>GGATCC</u>AGCTTAGCTCTTACCTGTTTTCGTCGT-3’). *Hind*III-*Bam*HI-cut CmYLCV was cloned into *Hin*dIII-*Bam*HI-cut pnCas9-PBE vector from Addgene (Addgene plasmid #98164) to build pCmYLCV-nCas9-PBE. To produce GFP-p1380N-CmYLCV-nCas9-PBE, the *Sbf*I-*Eco*RI-cut CmYLCV-nCas9-PBE fragment was ligated with the *Sbf*I-*Eco*RI-cut GFP-p1380N-Cas9 construct ^40^

From 35S-SpCas9p:DunLOBP ^43^, the AtU6-26 promoter was amplified again using AtU6-26-5-*Xho*I and AtU6-26-3-phos (5’-phosphorylated-AATCACTACTTCGACTCTAGCTGT-3’), and the sgRNA-ALSBE-NosT was PCR-amplified using sgRNA-ALSBE-P (5’-phosphorylated-GcaggtcccgcggaggatgatGTTTTAGAGCTAGAAATAGCAAGT-3’) and NosT-3-*Spe*I. Through three-way ligation, *Xho*I-cut AtU6-26 and *Spe*I-digested sgRNA-ALSBE-NosT were inserted into *Xho*I-*Xba*I-treated pUC-NosT-MCS to construct pUC-NosT-AtU6-26-sgRNA-ALSBE. Finally, the *Eco*RI-NosT-AtU6-26-sgRNA-ALSBE-NosT-*Pme*I fragment from pUC-NosT-AtU6-26-sgRNA-ALSBE were cloned into *Eco*RI-*Pme*I-cut GFP-p1380N-CmYLCV-nCas9-PBE to build GFP-p1380N-PBE:ALS (Extended Data Figure S14). The *Asc*I-*Pme*I-cut CmYLCV-nCas9-PBE:ALS fragment from GFP-p1380N-PBE:ALS was clone into *AscI-PmeI-cut* vector GFP-p1380N-ttLbCas12a:LOBP1-*Asc*I-*Xba*I-*Pme*I to form GFP-p1380N-ttLbCas12a:LOBP1-PBE:ALS (Extended Data Figure S14). All constructs were confirmed by sequencing.

### Plant transformation

The final constructs were transformed into either the *Agrobacterium* strain AGL1 (for tomato and potato) or EHA105 (for tobacco and citrus). For tomato transformation (cultivar Moneymaker), we followed the described protocol ^64^ with modifications. After co-cultivating the cotyledon explants on co-cultivation medium for three days, the explants were subcultured on a Murashige and Skoog (MS) regeneration medium with 2 mg/L zeatin riboside and 350 mg/L Timentin (for Agrobacterium elimination) at 30°C for 6 to 10 days. The explants were then subcultured on the same regeneration medium with 2 mg/L zeatin riboside, 350 mg/L Timentin, and 110 nM herbicide chlorsulfuron to select chlorsulfuron-resistant calli and shoots. The calli showing green fluorescence were discarded, while the non-fluorescent calli were kept as potential transgene-free, gene-edited transformants for further culture on chlorsulfuron-containing regeneration media. A similar protocol was used for tobacco transformation, using young sterile tobacco leaf discs as explants, and a regeneration medium containing 1 mg/L 6-Benzylaminopurine (BAP), 0.1 mg/L Naphthalene acetic acid (NAA), 350 mg/L Timentin, and 250 nM herbicide chlorsulfuron. For potato transformation, the tetraploid cultivar Atlanta plantlets were purchased from the University of Wisconsin and the University of Idaho. A similar protocol was used for potato transformation. Potato leaves were used as explants for transformation. The regeneration medium for potato transformation contains 1 mg/L Zeatin, 2 mg/L Gibberellic acid (GA), 350 mg/L Timentin, 100 nM herbicide chlorsulfuron. For citrus transformation, we followed the protocol we developed previously^10^. Plants were grown at room temperature (22 °C - 25 °C) with a 16-hour light/8-hour dark cycle. After rooting, the plants were transferred to a glasshouse.

### Canker symptom assay in citrus

Wild type, transgenic and transgene-free Pummelo plants were grown in a greenhouse at the Citrus Research and Education Center, University of Florida. Prior to *Xcc* treatment, all plants were trimmed to generate new shoots. Leaves of similar age were infiltrated with either *Xcc* or *Xcc*ΔpthA4:dLOB1.5 (5 × 10^8^ CFU/mL) using needleless syringes. At five days post inoculation (DPI), canker symptoms were observed and photographed.

### Microscopy analysis

An Omax camera was installed to a Zeiss Stemi SV11 dissecting microscope for photographing GFP fluorescence. Under illumination of the Stereo Microscope Fluorescence Adapter (NIGHTSEA), GFP fluorescence was visualized. Subsequently, the samples were photographed with the Omax Toupview software connected to the Omax camera.

### Genomic DNA extraction and genotyping

Genomic DNA was extracted from plant leaves with a cetyltrimethylammonium bromide (CTAB)-based genomic DNA extraction protocol we described previously^39, 65^. Detection of edits in the target genes was performed via PCR amplification of fragments spanning gRNAs or crRNAs, followed by cloning of PCR products into a cloning vector (Zero Blunt™ TOPO™ PCR Cloning Kit, Invitrogen) and Sanger sequencing. At least 10 clones for each gene from each plant were subjected to Sanger sequencing. Primers were designed for the detection of *GFP* fragment near the T-DNA right border, and *Cas12a* fragment near the T-DNA left border in the edited plant lines.

### Whole genome sequencing and data analysis

The 150-bp paired-end reads whole genome sequencing data were generated using the Illumina NovaSeq 6000 platform by Novogene. The raw reads were filtered using Fastp version 0.22.0 to remove low-quality reads. On average, more than 20.6 and 48.4Gb of high-quality data were generated for each citrus Pummelo and tomato plant sample, respectively. The high-quality paired-end short genomic reads were mapped to the reference genomes of citrus Pummelo (C. maxima) or tomato using Bowtie2 software version 2.2.6 ^66^. The mutations (single nucleotide polymorphisms, deletions, and insertions) in the gene-edited plant genomes were generated using the SAMtools package version 1.2 ^67^ and Deepvariant program version 1.4.0 ^68^. The mutations were filtered based on quality and sequence depth, and the target site mutations were visualized using IGV software version 2.15.4 ^69^. The off-target sites were predicted using CRISPR-P 2.0 ^70^ and the Cas-OFFinder program ^71^ and aligning target sequence with whole genome using blast program. Based on the mapping results, mutations of off-target sites were detected using the SAMtools package version 1.2 and deepvariant program version 1.4.0.

## Data availability

The raw reads of genome resequencing for pumelo plants were deposited in the NCBI Bioproject database under the accession number PRJNA931434. The reference genome of pumelo was downloaded from public citrus genome database CPBD: Citrus Pan-genome to Breeding Database (http://citrus.hzau.edu.cn/index.php). The raw reads of genome resequencing for tomato plants were deposited in the NCBI Bioproject database under the accession number PRJNA931572. The reference genome of tomato was downloaded from public tomato genome database of International Tomato Genome Sequencing Project https://solgenomics.net/organism/Solanum_lycopersicum/genome).

## Acknowledgements

We thank Wang lab members for constructive suggestions and insightful discussions. This project was supported by funding from Florida Citrus Initiative Program, Citrus Research and Development Foundation, U.S. Department of Agriculture National Institute of Food and Agriculture grants 2022-70029-38471, 2021-67013-34588, 2018-70016-27412 and 2016-70016-24833, FDACS Specialty Crop Block Grant Program (N. Wang).

## Author Contributions

X.H., H.J. and N.W. conceptualized and designed the experiments. X.H., H.J., and Y.W. performed the experiments. J.X. and J.W. performed bioinformatics. X.H., H.J. and N.W. wrote the manuscript with input from all co-authors.

## Competing interests

N. W., H. J. and X. H. filed a PCT patent application based on the results reported in this paper. All other authors declare no competing financial interests.

**Correspondence and requests** for materials should be addressed to N. Wang.

