## Supplementary Information for "Efficient transgene-free genome editing in plants in the T0 generation based on a co-editing strategy"

**Extended Data Table S1.** Off-target homologous sites with up to 4 mismatches to the target site in the *SlALS1* gene.

| Sequence | Off score | Number of mismatches | Region | Edited |
|---|---|---|---|---|
| CAAGTGCCGAGGAGGATGATTGG | 0.667 | 1 | CDS | No |
| TTAGTGCCAAGGAAAATGATAGG | 0.513 | 4 | Intergenic | No |
| GAGGTGACAAGGAAGATGATGGG | 0.314 | 4 | Intergenic | No |
| AAAGTTCCAAGCAGGATAATTGG | 0.244 | 4 | Intergenic | No |
| AAAGAGACAAGGAGGATGAAGGG | 0.211 | 4 | Intergenic | No |
| CAGGGGCTAAGCAGGATGATGGG | 0.181 | 4 | CDS | No |
| GAAGTGCCAAGAATGAAGATGGG | 0.121 | 4 | CDS | No |
| TTTGTGCCATGGAGGATGATTAG | 0.117 | 4 | Intergenic | No |
| CATGTGCCAAGGAGGATGAATAG | 0.103 | 2 | Intergenic | No |
| CAAGGGCCAAGTAAGATGTTGGG | 0.099 | 4 | intron | No |
| CATGTGCCATGGAGGATGAATAG | 0.091 | 3 | Intergenic | No |
| CAAGAGCAAAGGAGGATGATGAG | 0.084 | 2 | Intergenic | No |
| GATGTGCCATGGAGGATGAATAG | 0.083 | 4 | Intergenic | No |
| GATGTGCCATGGAGGATGAATAG | 0.083 | 4 | Intergenic | No |
| GATGTGCCATGGAGGATGAATAG | 0.083 | 4 | Intergenic | No |
| GATGTGCCATGGAGGATGAATAG | 0.083 | 4 | Intergenic | No |
| CTAGTGAAATGGAGGATGATTAG | 0.081 | 4 | Intergenic | No |
| CAAGTGCCAAGGAACAAAATTGG | 0.076 | 4 | CDS | No |
| CCACTGACAAGGAGAATGATTAG | 0.076 | 4 | Intergenic | No |
| TATGTGCCAAGGAGGATTAATAG | 0.069 | 4 | Intergenic | No |

**Extended Data Table S2. Summary of the whole genome sequencing data**

| Sample | Total raw data (Gb) | Total high quality data (Gb) | Sequencing coverage (×) | Construct sequence |
|---|---|---|---|---|
| *SlALS* edited line 2 | 49.12 | 48.63 | 59.53 | No |
| Transgenic control line EPGFP | 43.36 | 42.78 | 52.36 | Yes (1830 reads) |
| *SlRbohD* edited line 1 | 31.52 | 31.14 | 38.11 | No |
| *SlRbohD* edited line 2 | 25.45 | 25.10 | 30.72 | No |
| *SlRbohD* edited line 6 | 22.28 | 21.96 | 26.88 | Yes (2716 reads) |
| *SlRbohD* edited line 8 | 46.49 | 45.87 | 56.14 | No |
| *SlER* edited line 4 | 22.91 | 22.61 | 27.67 | No |
| *SlEDS1/SlPAD4* edited line 8 | 25.07 | 24.80 | 30.36 | Yes (281 reads) |
| *SlEDS1/SlPAD4* edited line 18 | 26.76 | 26.44 | 32.36 | No |
| $Pum_{NoGFP}3$ | 20.68 | 20.23 | 58.47 | No |

**Extended Data Table S3. Off-target homologous sites to the target site.**

| **Targets** | **Sequence** | **Number of mismatches** | **Off-target mutations** |
|---|---|---|---|
| For *SlER* off-target | TTTG ATTCTGTTGTGTGTGTTGATGGT | 4 | No |
| For *SlER* off-target | TTTG ATTCTGTTGCGTCTGTTGATGGT | 4 | No |
| For *SlEDS1* off-target | TTTA GAGAGAAAGAGATCATACCCACA | 4 | No |
| For *SlINVINH1* off-target | TTTG GTGTATTTTTGCATGTTGTTTCC | 4 | No |
| For *SlINVINH1* off-target | TTTC GAATGTTCTTGAATGTTGTTTCT | 4 | No |
| For *SlINVINH1* off-target | TTTG GTGTTTTCTTGCATGTATTCTTT | 4 | No |
| For *SlRBL2* off-target | TTTC TTTTCAGCCCTTGAGGGAAAATC | 4 | No |

**Extended Data Table S4. Primers**

| Primer | Sequence | Purpose |
|---|---|---|
| Cas12half-F1 | ACAAAAACTTTTCGCGGACCGATG | Half Cas12a-D156R amplification to make CBE-Cas12a |
| Cas12half-R1 | CTATGTCCTGATAGCGGTCAGCCACCTTATCTTTAATCATATTCC | Half Cas12a-D156R amplification to make CBE-Cas12a |
| NtALS-F1 | TGCACAAGTACCACGTAGGATGAT | For NtALS base editing |
| NtALS-R1 | AAACATCATCCTACGTGGTACTTG | For NtALS base editing |
| SlALS1-F1 | TGCACAAGTGCCAAGGAGGATGAT | For SlALS1 base editing |
| SlALS1-R1 | AAACATCATCCTCCTTGGCACTTG | For SlALS1 base editing |
| CsALS-F | ATGGTCTCGTGCACAGGTCCCTCGGAGGATGATGTTTAAGAGCTATGCTGGAAACA | For CsALS base editing |
| CsALS-R | TaGGTCTCCAAACATCATCCTCCGCGGGACCTGTGCACCAGCCGGGAATCGAAC | For CsALS base editing |
| StALS1-F | TGCACAAGTGCCGAGGAGGATGAT | For StALS1 base editing |
| StALS1-R | AAACATCATCCTCCTCGGCACTTG | For StALS1 base editing |
| NtALSgt-F1 | TTTTCCGACCAGCGCCGTTGA | For NtALS genotyping |
| NtALSgt-R1 | TTCCACAAAACGCCTTAACTCCT | For NtALS genotyping |
| SlALS1-F | GTGGTTTTGCCGTTGCCAAT | For SlALS1 genotyping |
| SlALS1-R | CAGCTCCTCACTTGATTGCG | For SlALS1 genotyping |
| CsALSgt-F2 | CGCCACCAACCTGGTCAGCG | For CsALS genotyping |
| CsALSgt-R2 | CCTGGCCGCCCAGATGTTGC | For CsALS genotyping |
| NtPDScr-F1 | TAGAtGGCAGATCAGAGCAAAGCAAAAA | For NtPDS gene editing |
| NtPDScr-R1 | GGCCTTTTTGCTTTGCTCTGATCTGCCA | For NtPDS gene editing |
| CsPDScr-F1 | TAGAtAGGTAACTGAAGCTTGAGGATAT | For CsPDS gene editing |
| CsPDScr-R1 | GGCCATATCCTCAAGCTTCAGTTACCTA | For CsPDS gene editing |
| NtPDSgt-F2 | CTATATCCGTTCATGTGTTCTTTA | For NtPDS genotyping |
| NtPDSgt-R2 | ATAGGAGATCTTTGCAAGGGCC | For NtPDS genotyping |
| LOBcr-F1 | TAGAtCTTTCTCTATATAAACCCCTTTT | For LOB1 gene editing |
| LOBcr-R1 | GGCCAAAAGGGGTTTATATAGAGAAAGA | For LOB1 gene editing |

| | | |
|---|---|---|
| SlERcr-F1 | TAGAtGTTCTGTGGTGTCTGATGATGGT | For SlER gene editing |
| SlERcr-R1 | GGCCACCATCATCAGACACCACAGAACA | For SlER gene editing |
| SlERmd-F1 | AGACTTGAGTCTACCATGGCATC | For SlER genotyping |
| SlERmd-R1 | GACTTCTTAATTTCCAACAATGCAG | For SlER genotyping |
| SlRBL2cr-F1 | TAGAtGTTTCAGCCATTCAGTGAAAATC | For SlRBL2 gene editing |
| SlRBL2cr-R1 | GGCCGATTTTCACTGAATGGCTGAAACA | For SlRBL2 gene editing |
| SlRBL2md-F1 | AACTCAGTATACCCAGTAGAGCCA | For SlRBL2 genotyping |
| SlRBL2md-R1 | TCAATTCACTTCATTGATCACTAGC | For SlRBL2 genotyping |
| SlRbohDcr-F1 | TAGAtATATACCGGAATCTACTGACATC | For SlRbohD gene editing, 1 crRNA |
| SlRbohDcr-R1 | GGCCGATGTCAGTAGATTCCGGTATATA | For SlRbohD gene editing, 1 crRNA |
| SlRbohDcr-F2 | TtcgtctctTAGATATATACCGGAATCTACTGACATCGGCCGGCATGGTCCCAGC | For SlRbohD gene editing, 2 crRNAs |
| SlRbohDcr-R2 | aaCGTCTCAGGCCTACCTTGGGAGCTTCACTTGTACATCTACACTTAGTAGAAATTAGAC | For SlRbohD gene editing, 2 crRNAs |
| SlRbohDmd-F1 | GATGCAAAATTCGGAAAATCATCACC | For SlRbohD genotyping |
| SlRbohDmd-R1 | ATGAGCAGCAGCTGATTTATTCCTG | For SlRbohD genotyping |
| SlEDS1cr-F1 | TtcgtctctTAGATGAGAGAAAGAGATCAACACCACTGGCCGGCATGGTCCCAGC | For SlEDS1 and SlPAD4 gene editing |
| SlPAD4cr-R1 | aaCGTCTCAGGCCGCTTATGTGGGATTCTCCGGCGTATCTACACTTAGTAGAAATTAGAC | For SlEDS1 and SlPAD4 gene editing |
| SlEDS1md-F1 | CTCAATCTTTCTTCAAGGATTCTTG | For SlEDS1 genotyping |
| SlEDS1md-R1 | CTCATTTTTAAGTGAAGATTTGTCC | For SlEDS1 genotyping |
| SlPAD4md-F1 | TTCGAGTCTAGTGAAACTTTAGCAG | For SlPAD4 genotyping |
| SlPAD4md-R1 | AGCAACAACTGAGAGTGAATAAATTG | For SlPAD4 genotyping |
| SlDMR6cr-F1 | TtcgtctctTAGATACAATCGACCACTTCCGATAGACGGCCGGCATGGTCCCAGC | For SlDMR6 and SlInvI gene editing |
| SlINVIcr-R1 | aaCGTCTCAGGCCAGAGACAACATGCAAGAACACACATCTACACTTAGTAGAAATTAGAC | For SlDMR6 and SlInvI gene editing |
| SlDMR6md-F1 | AACCAAAGTTATTTCTAGCGGAATC | For SlDMR6 genotyping |
| SlDMR6md-R1 | TCATGGTCTTTGAAGGATCATCTG | For SlDMR6 genotyping |
| SlInvmd-F1 | CCTAATAATGTTTCTTGCTATGTTGC | For SlINV genotyping |
| SlInvmd-R1 | ATAAAATCCTAATTCATCATAGTCGAG | For SlINV genotyping |

| | | |
|---|---|---|
| SlSOBIRcr-F1 | TtcgtctctTAGATCTGGCAGAATACCTCAATCCTTGGGCCGGCATGGTCCCAGC | For SlSobir1 and SlSobir2 gene editing (2 crRNAs for each gene) |
| SlSOBIRcr-R1 | ttCGTCTCAACCTAGTACAGTTGTAGTACCAGATCTACACTTAGTAGAAATTAGACG | For SlSobir1 and SlSobir2 gene editing (2 crRNAs for each gene) |
| SlSOBIRcr-F2 | TtcgtctctAGGTGGCCGGCATGGTCCCAGC | For SlSobir1 and SlSobir2 gene editing (2 crRNAs for each gene) |
| SlSOBIRcr-R2 | ttCGTCTCAGGAACTTTGTCTTCTGCTATTGGATCTACACTTAGTAGAAATTAGACG | For SlSobir1 and SlSobir2 gene editing (2 crRNAs for each gene) |
| SlSOBIRcr-F3 | TtcgtctctTTCCGGCCGGCATGGTCCCAGC | For SlSobir1 and SlSobir2 gene editing (2 crRNAs for each gene) |
| SlSOBIRcr-R3 | aaCGTCTCAGGCCGCATTTATCAGCAGATTTGAATCATCTACACTTAGTAGAAATTAGACG | For SlSobir1 and SlSobir2 gene editing (2 crRNAs for each gene) |
| SlSobirmd-F1 | AATATCGTGCGAGCGTAGACCG | For SlSobir1 genotyping |
| SlSobirmd-R1 | AGCTGGAGCTGGAACCGAAGC | For SlSobir1 genotyping |
| SlSobirmd-F2 | ACCAACTGAAATAGTTGACTCCAG | For SlSobir2 genotyping |
| SlSobirmd-R2 | GTTAAGCCTGGATCATTTCTGATC | For SlSobir2 genotyping |
| StDMR6cr-F1 | TAGAtAAGGATCATCTGAATACAATTTT | For StDMR6 gene editing, 1 crRNA |
| StDMR6cr-R1 | GGCCAAAATTGTATTCAGATGATCCTTA | For StDMR6 gene editing, 1 crRNA |
| StDMR6-F3 | TtcgtctctTAGATGGTAATTAATCATGGTGTACCAAGGCCGGCATGGTCCCAGC | For StDMR6 gene editing, 3 crRNAs |
| StDMR6-R3 | ttCGTCTCAGAATCCACCACTCTACTCTCCCTATCTACACTTAGTAGAAATTAGACG | For StDMR6 gene editing, 3 crRNAs |
| StDMR6-F4 | TtcgtctctATTCGGCCGGCATGGTCCCAGC | For StDMR6 gene editing, 3 crRNAs |
| StDMR6-R4 | aaCGTCTCAGGCCAAAATTGTATTCAGATGATCCTTATCTACACTTAGTAGAAATTAGACG | For StDMR6 gene editing, 3 crRNAs |
| StDMR6md-F3 | CCATGGAAACGAAAGTTATTTCC | For StDMR6 genotyping |
| StDMR6md-F4 | TTAATTATATTTATTTGCTACGCCAT | For StDMR6 genotyping |
| StDMR6md-R1 | GATTAGAAGGCCATTCAGGAGCA | For StDMR6 genotyping |
| Mini-F1 | ATAAAACAATATTATCCCTGGCAGACATACTGTCCCACAAA | For PCR from PUC57-mini-crRNA |
| Mini-R1 | TCTGGCGCGCCGATCGCCGATCTAGTAACATAGATGACACCG | For PCR from PUC57-mini-crRNA |
| Cas12gt-F1 | CTAAGAAGAATAACGTCTTTGATTGG | Detection of Cas12a in the edited plants |
| Cas12gt-R1 | TGCTTTACGGATGTCTGAGCATAC | Detection of Cas12a in the edited plants |
| GFPgt-F2 | TTTCACTTCTCCTATCATTATCCTC | Detection of GFP in the edited plants |
| GFPgt-R2 | CATGGCGCTCTTGAAGAAGTCG | Detection of GFP in the edited plants |

| | | |
|---|---|---|
| SlERoff1-F1 | TGATGGGCCAAATGGAGCAATTTG | For tomato off-target detection |
| SlERoff1-R1 | TGAAATTACCGAATAATGTCAATATCC | For tomato off-target detection |
| SlERoff2-F1 | CAATTCACACTTGCTAAATCCCAAC | For tomato off-target detection |
| SlERoff2-R1 | TTTAGGTTAGAACTGGGAATTCCTC | For tomato off-target detection |
| SlINVoff1-F1 | GTCAGATCCAAATTCATTGTTATCCA | For tomato off-target detection |
| SlINVoff1-R1 | CACATTCCTTTAAGGGATGTATCCA | For tomato off-target detection |
| SlINVoff2-F1 | GATTTCTCTTCTTAAATTCACTTCGA | For tomato off-target detection |
| SlINVoff2-R1 | CGTTCAAGATTCGATAAGATTGTCG | For tomato off-target detection |
| SlINVoff3-F1 | CCAAAGTTGTCTCCACTTAGTGAC | For tomato off-target detection |
| SlINVoff3-R1 | CATGAAAGTTTCCTTTCATCCAGC | For tomato off-target detection |
| SlEDS1off1-F1 | AAGAAGTTTACAACTAATATGTCACCA | For tomato off-target detection |
| SlEDS1off1-R1 | ACAGCCAGAGTGCAATAGACTTGA | For tomato off-target detection |
| SlRBL2off1-F1 | AATCATGCTTCTTCCTATGCTGTC | For tomato off-target detection |
| SlRBL2off1-R1 | ATAACTAATTGTACCTCCTATACTTCA | For tomato off-target detection |

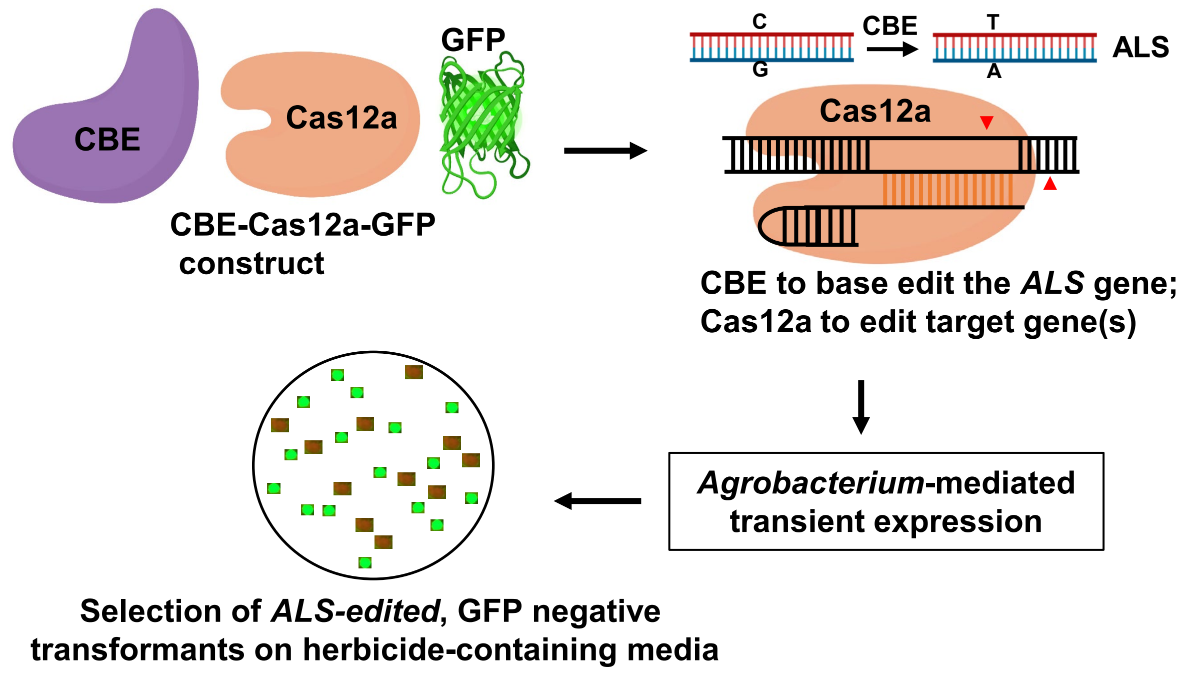


**Extended Data Figure S1. Generation of transgene-free, gene-edited plants in the first generation using the CBE-Cas12a-GFP construct.** The CBE-Cas12a-GFP construct consists of CBE, Cas12a, and GFP, each driven by its own promoter. CBE and its gRNA are used to base edit the *ALS* gene to confer resistance to herbicide chlorsulfuron; Cas12a and its crRNA are used for target gene editing; GPF is used for selecting transgene-free transformants. The construct is introduced into plants through *Agrobacterium*-mediated transient expression. Transformants are screened on media containing chlorsulfuron. Putative transgene-free, gene-edited plants are kept if they lack green fluorescence. Plants with fluorescence are discarded. Regenerants without GFP are subjected to genotyping and further analysis.

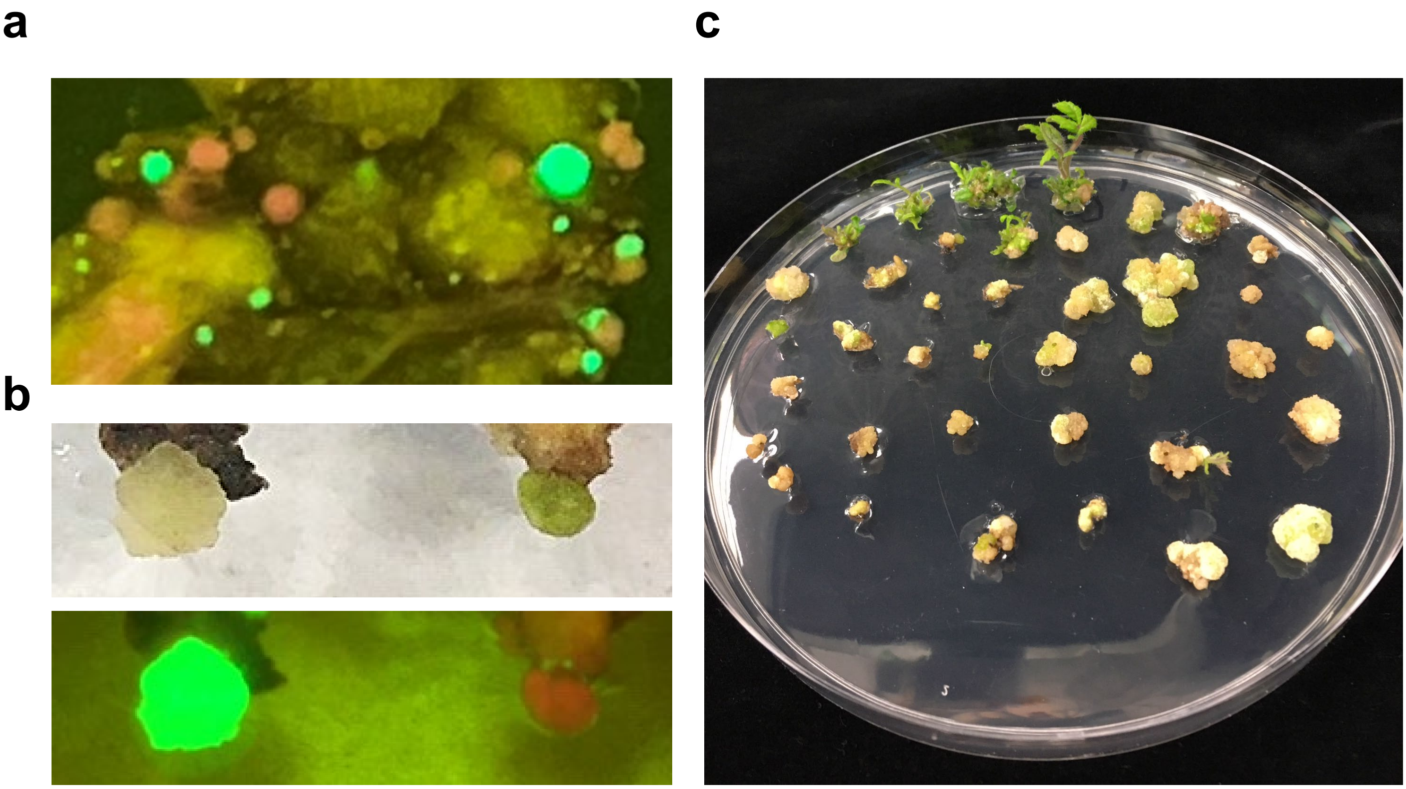


**Extended Data Figure S2. Tomato calli and shoots selected on media containing chlorsulfuron herbicide**. **a,** Tomato calli were grown on the medium containing 110 nM chlorsulfuron. **b,** Close-up view of calli from the selection medium shown in a. **c,** Tomato calli and shoots selected on medium containing 110 nM chlorsulfuron.

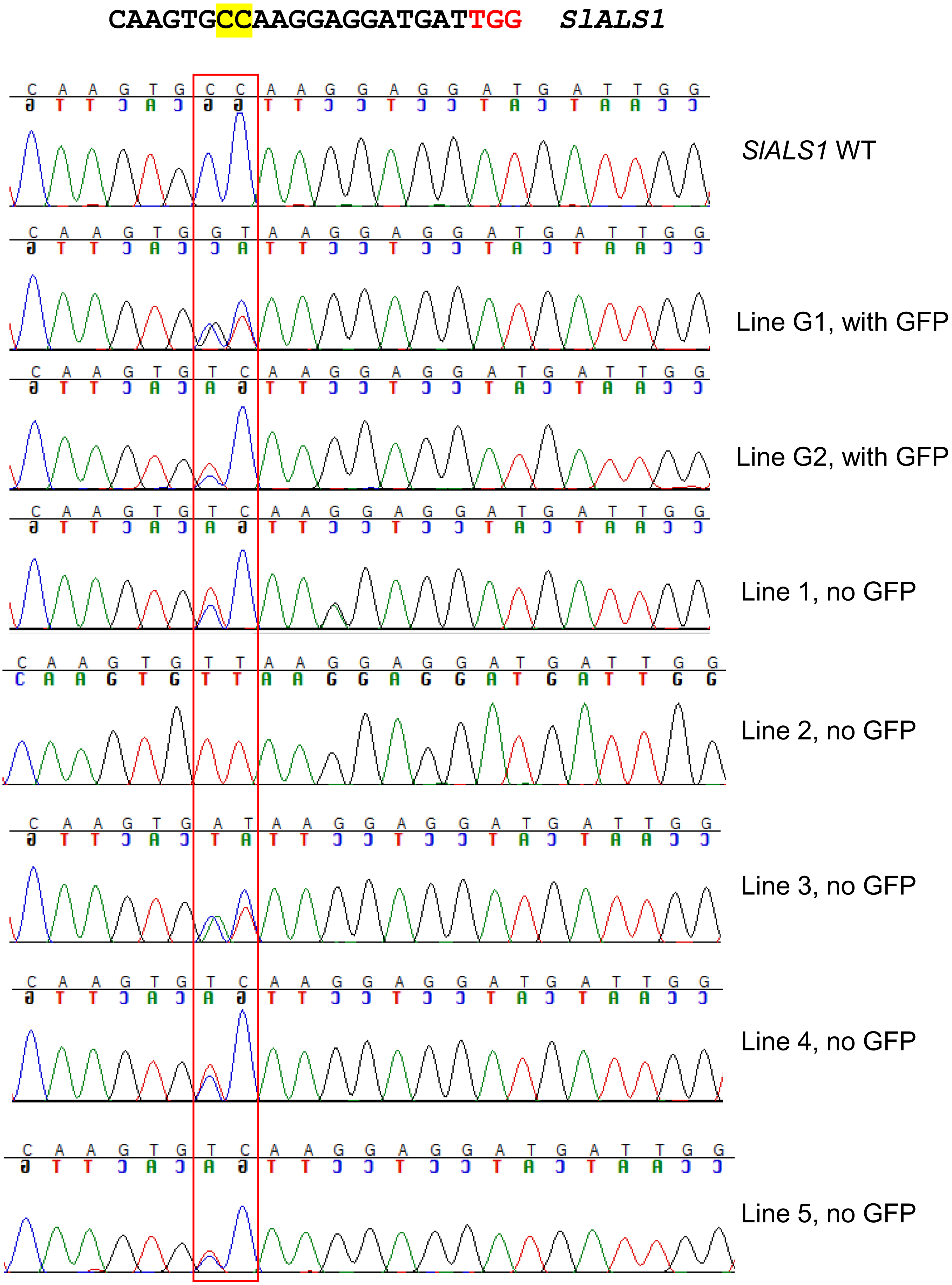


**Extended Data Figure S3. *SlALS1* genotyping of chlorsulfuron-resistant tomato plant lines by Sanger sequencing.** Direct sequencing of the PCR products was conducted. Direct sequencing indicates both alleles were edited in line 2, whereas only one allele was edited in line 1, 3, 4 and 5.

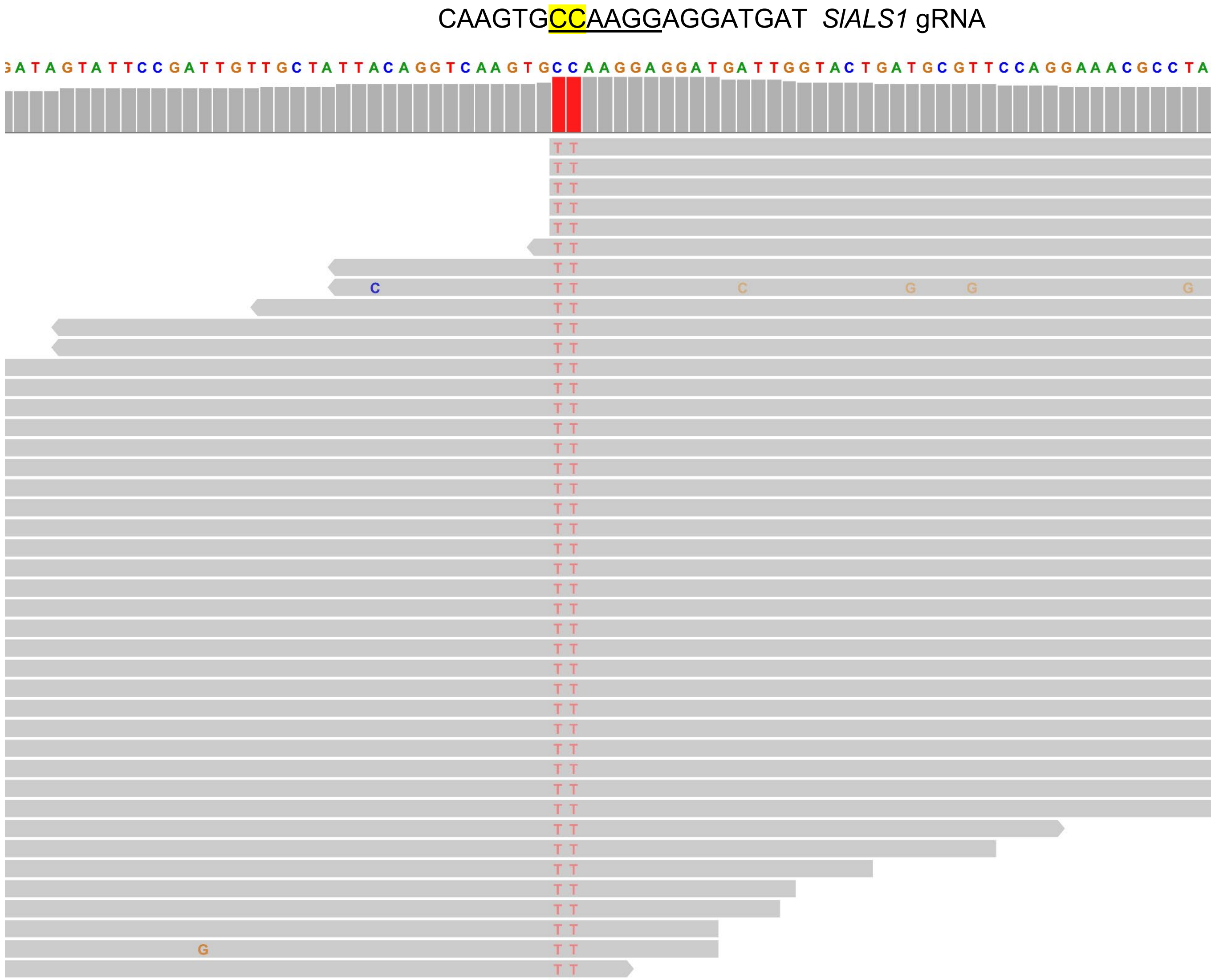


**Extended Data Figure S4. Whole genome sequencing confirms base editing of *SlALS1* in the *SlALS1-edited*, GFP negative tomato line L2.**

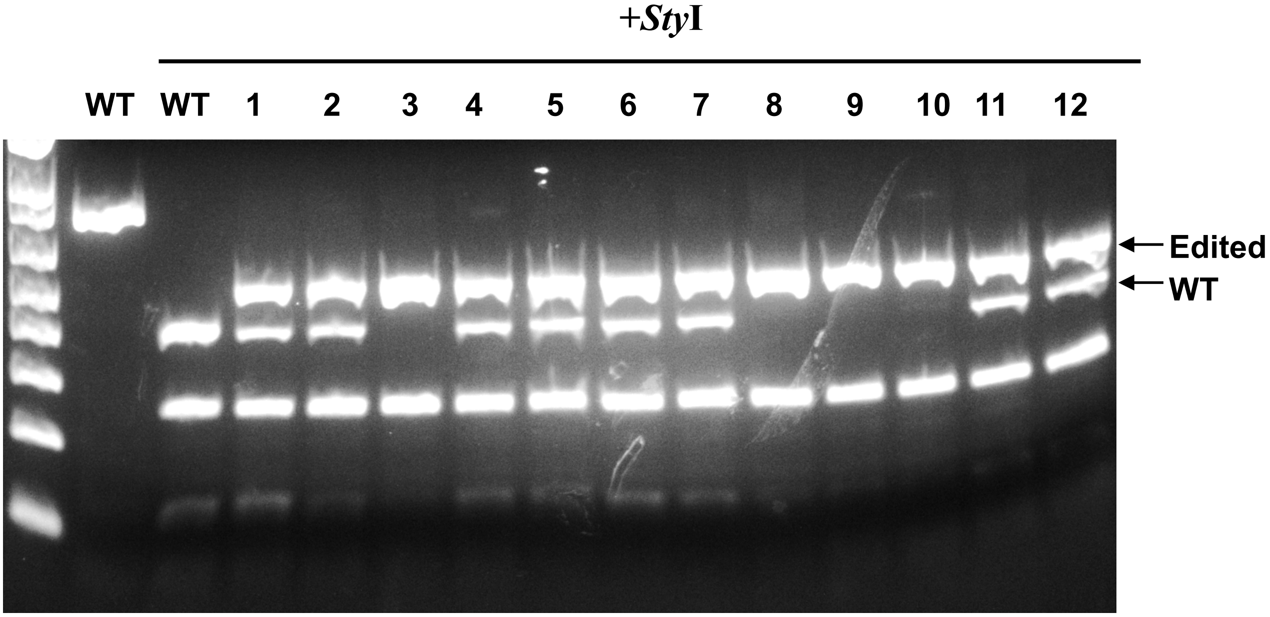


**Extended Data Figure S5. *SlALS* gene genotyping of chlorsulfuron-resistant, non-GFP tomato regenerants generated after CBE-Cas12a-GFP-SlALS1-SlER construct transformation.** PCR amplicons spanning the *SlALS1* gRNA region were subjected to restriction enzyme digestion with *Sty*I. Editing of the targeted nucleotides abolishes the *Sty*I recognition site, resulting in resistance to *Sty*I digestion.

### Line 2 biallelic

CTTCAGCTTTG**GTTCTGTGGTGTCTGATGATGGT**GAGTAGAGTAGAGTAG *SlER* WT
CTTCAGCTTTG**GTTCTGTGGTGT-----------**GAGTAGAGTAGAGTAG allele 1, -11 bp
CTTCAGCTTTG**GTTCTGTGGT-------------**GAGTAGAGTAGAGTAG allele 2, -13 bp

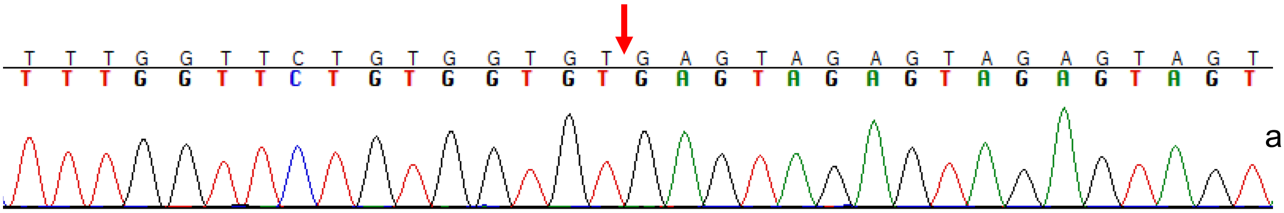


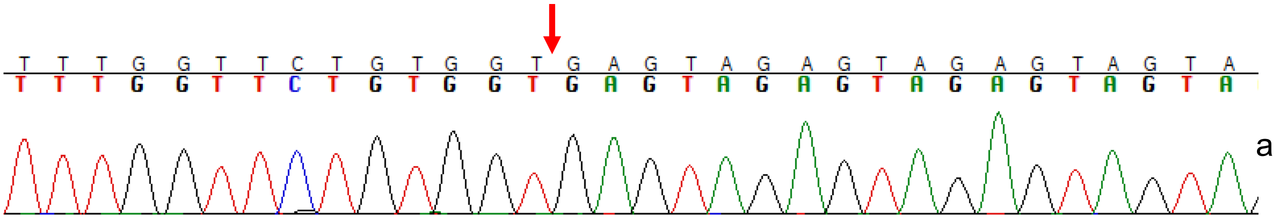


### Line 4 biallelic

CTTCAGCTTTG**GTTCTGTGGTGTCTGATGATGGT**GAGTAGAGTAGAGTAG *SlER* WT
CTTCAGCTTTG**GTTCTGTGGTGTCTG-------T**GAGTAGAGTAGAGTAGTAG allele 1, -7 bp
CTTCAGCTTTG**GTTCTGTGGTGTCTGATGA----**-------GTAGAGTAGTAG allele 2, -11 bp

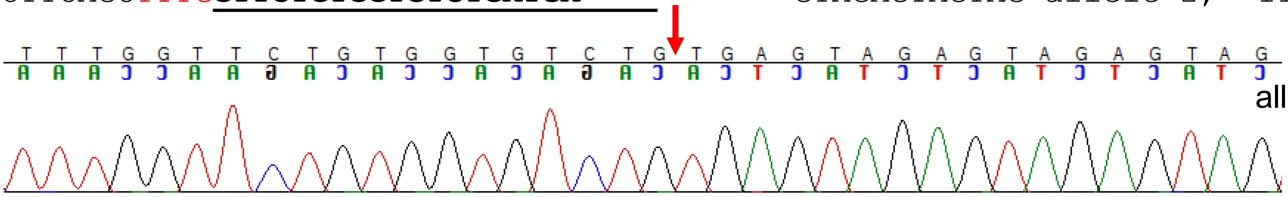


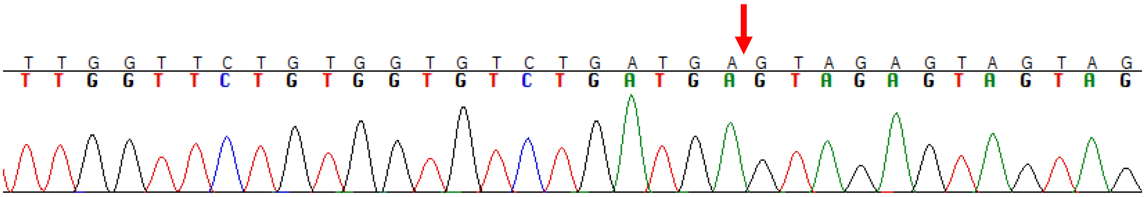


### Line 5 biallelic

CTTCAGCTTTG**GTTCTGTGGTGTCTGATGATGGT**GAGTAGAGTAGAGTAGTAG *SlER* WT
CTTCAGCTTTG**GTTCTGTGGTGTCTG--------**------AGTAGAGTAGTAG allele 1, -14 bp
CTTCAGCTTTG**GTTCTGTGGTGTCTG--------**---------AGAGTAGTAG allele 2, -17 bp

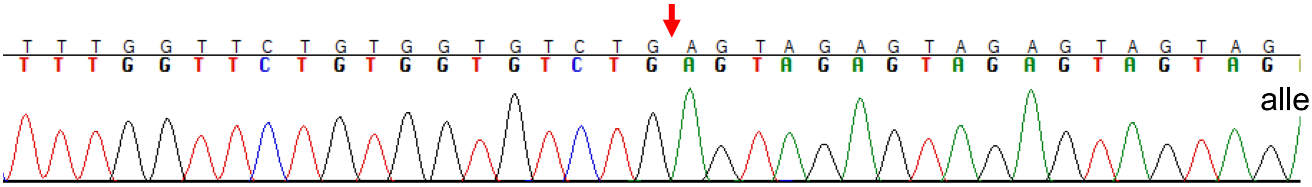


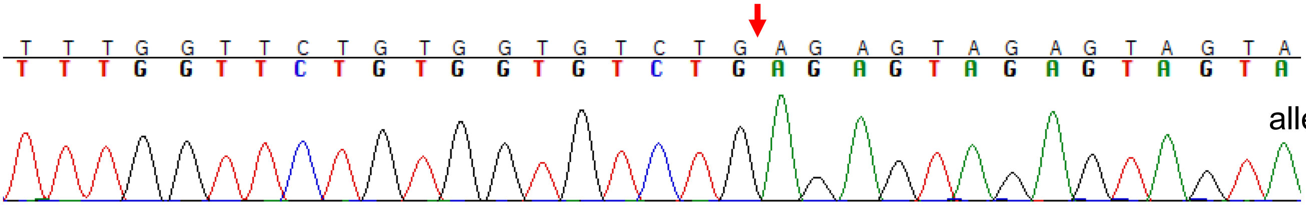


**Extended Data Figure S6. Chromatograms for *SlER* genotypes of tomato GFP-negative *sler* biallelic mutants.** At least 16 clones for each gene from each plant were subjected to Sanger sequencing.

**a**

AGCTTTG**GTTCTGTGGTGTCTG-------T**GAGTAGAGTAGAGTAG allele 1, -7 bp
AGCTTTG**GTTCTGTGGTGTCTGATGA---**---------GTAGAGTAG allele 2, -11 bp
Plant 1, biallelic

AGCTTTG**GTTCTGTGGTGTCTG-------T**GAGTAGAGTAGAGTAG allele 1, -7 bp
Plant 2, homozygous

AGCTTTG**GTTCTGTGGTGTCTG-------T**GAGTAGAGTAGAGTAG allele 1, -7 bp
AGCTTTG**GTTCTGTGGTGTCTGATGA---**---------GTAGAGTAG allele 2, -11 bp
Plant 3, biallelic

AGCTTTG**GTTCTGTGGTGTCTG-------T**GAGTAGAGTAGAGTAG allele 1, -7 bp
Plant 4, homozygous

AGCTTTG**GTTCTGTGGTGTCTG-------T**GAGTAGAGTAGAGTAG allele 1, -7 bp
AGCTTTG**GTTCTGTGGTGTCTGATGA---**---------GTAGAGTAG allele 2, -11 bp
Plant 5, biallelic

AGCTTTG**GTTCTGTGGTGTCTG-------T**GAGTAGAGTAGAGTAG allele 1, -7 bp
AGCTTTG**GTTCTGTGGTGTCTGATGA---**---------GTAGAGTAG allele 2, -11 bp
Plant 6, biallelic

AGCTTTG**GTTCTGTGGTGTCTG-------T**GAGTAGAGTAGAGTAG allele 1, -7 bp
AGCTTTG**GTTCTGTGGTGTCTGATGA---**---------GTAGAGTAG allele 2, -11 bp
Plant 7, biallelic

AGCTTTG**GTTCTGTGGTGTCTG-------T**GAGTAGAGTAGAGTAG allele 1, -7 bp
AGCTTTG**GTTCTGTGGTGTCTGATGA---**---------GTAGAGTAG allele 2, -11 bp
Plant 8, biallelic

AGCTTTG**GTTCTGTGGTGTCTG-------T**GAGTAGAGTAGAGTAG allele 1, -7 bp
AGCTTTG**GTTCTGTGGTGTCTGATGA---**---------GTAGAGTAG allele 2, -11 bp
Plant 9, biallelic

AGCTTTG**GTTCTGTGGTGTCTG-------T**GAGTAGAGTAGAGTAG allele 1, -7 bp
AGCTTTG**GTTCTGTGGTGTCTGATGA---**---------GTAGAGTAG allele 2, -11 bp
Plant 10, biallelic

AGCTTTG**GTTCTGTGGTGTCTGATGA---**---------GTAGAGTAG allele 2, -11 bp
Plant 11, homozygous

AGCTTTG**GTTCTGTGGTGTCTG-------T**GAGTAGAGTAGAGTAG allele 1, -7 bp
AGCTTTG**GTTCTGTGGTGTCTGATGA---**---------GTAGAGTAG allele 2, -11 bp
Plant 12, biallelic

AGCTTTG**GTTCTGTGGTGTCTG-------T**GAGTAGAGTAGAGTAG allele 1, -7 bp
Plant 13, homozygous

AGCTTTG**GTTCTGTGGTGTCTGATGA---**---------GTAGAGTAG allele 2, -11 bp
Plant 14, homozygous

**b**

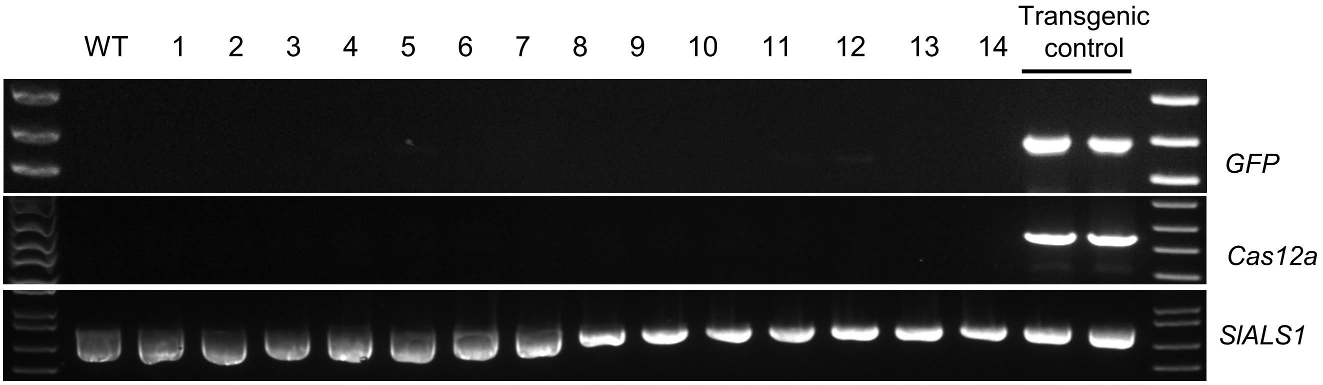


**Extended Data Figure S7. Heritability test. a,** *SlER* genotypes of the progenies of *sler-4*. Seeds of the *sler-4* line were germinated, and the seedlings were genotyped at *SlER*. **b,** PCR amplification of *GFP* and *Cas12a* for lines listed in a. At least 16 clones for each gene from each plant were subjected to Sanger sequencing.

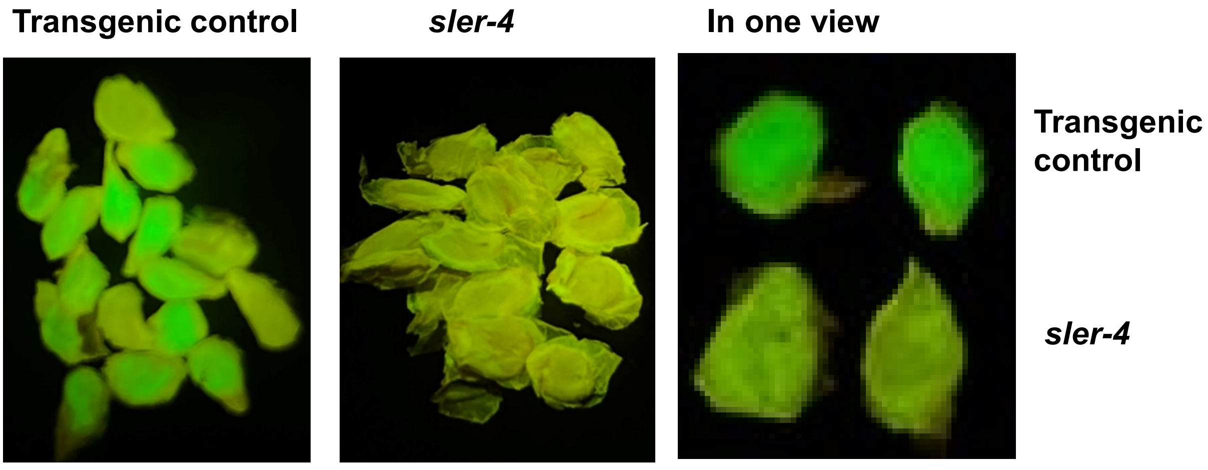


**Extended Data Figure S8. Detection of green fluorescence in *sler-4* and transgenic control seeds under a GFP filter.**

**a**

GGGTAGGTTTTCGTTTCAGCCATTCAGTGAAAATCCTCTTCTTGGGCCTTCTTCTACCA WT *SlRBL2*

**Line 4**
GGGTAGGTTTTCGTTTCAGCCATTCAG----AATCCTCTTCTTGGGCCTTCTTCTACCA allele 1, -4 bp
GGGTAGGTTTTCGTTTCAGC-------------TCCTCTTCTTGGGCCTTCTTCTACCA allele 2, -13 bp

**Line 8**
GGGTAGGTTTTCGTTTCAGCCATTCAGT-----------------GCCTTCTTCTACCA homozygous, -17 bp

**Line 9**
GGGTAGGTTTTCGTTTCAGCCATTC-------------TTCTTGGGCCTTCTTCTACCA allele 1, -13 bp
GGGTAGGTTTTCGTTTCAGCCATTCAGT---------CTTCTTGGGCCTTCTTCTACCA allele 2, -9 bp

**Line 11**
GGGTAGGTTTTCGTTTCAGCCATTCA--------CCTCTTCTTGGGCCTTCTTCTACCA allele 1, -8 bp
GGGTAGGTTTTCGTTTCAGCCATTC-------------TTCTTGGGCCTTCTTCTACCA allele 2, -13 bp

**Line 12**
GGGTAGGTTTTCGTTTCAGCCATTCA--------CCTCTTCTTGGGCCTTCTTCTACCA homozygous, -8 bp

**b**

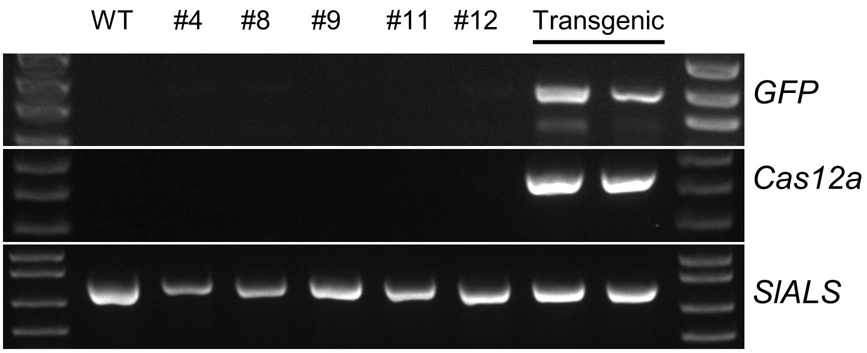


**Extended Data Figure S9. Transgene-free editing of *SlRBL2* in the T0 generation. a,** Genotypes of homozygous/biallelic mutants of *SlRBL2* (Solyc09g010880) that were without green fluorescence. **b**, Detection of *GFP* and *Cas12a* by PCR. Transgenic lines were used as a positive control.

GGAGGAAAAAAGAGCGCGAGGTTTAATATACCGGAATCTACTGACATCGGAACTAGTGCAGGAGCCGGCGCCAAGTCCAATGA
CGATGCTTATGTGGAAATCACTCTCGATGTTCGTGAAGACTCTGTGGCTGTCCACAGTGTGAAAACTGCCGGCGGTGCTGACG
TGGAAGATCCGGAGTTGGCTTTATTGGCTAAAGGTTTGGAGAAGAAATCTACCTTGGGAGCTTCACTTGTACGAAATGCTTCT
TCGAGAATCCGGCAA SlRbohD WT

line 1, biallelic

GAGCGCGAGGTTTAATATACCGGAATCTACTGACATCGGAACTAGTGCAGGAGCCGG *SlRbohD* WT
GAGCGCGAGGTTTAATATACCGGAATCTAC----ATCGGAACTAGTGCAGGAGCCGG line 1, -4bp
GAGCGCGAGGTTTAATATACCGGAA--------CATCGGAACTAGTGCAGGAGCCGG line 1, -8bp

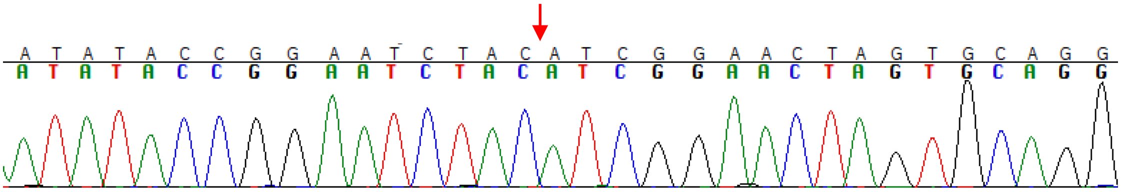

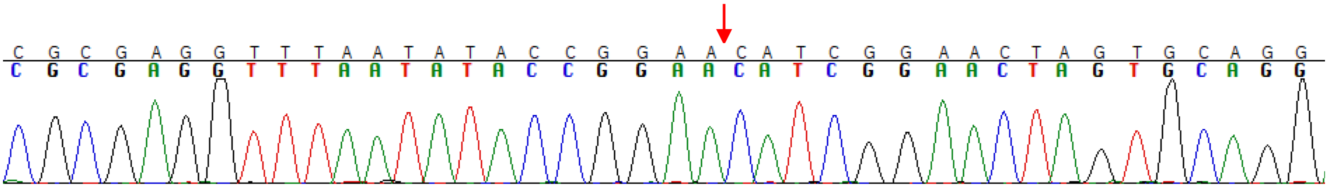

line 2, homozygous

AAAGAGCGCGAGGTTTAATATACCGGAATC-----------//------AAAATACGAAATGCTTCTTCGA -197bp, +4bp

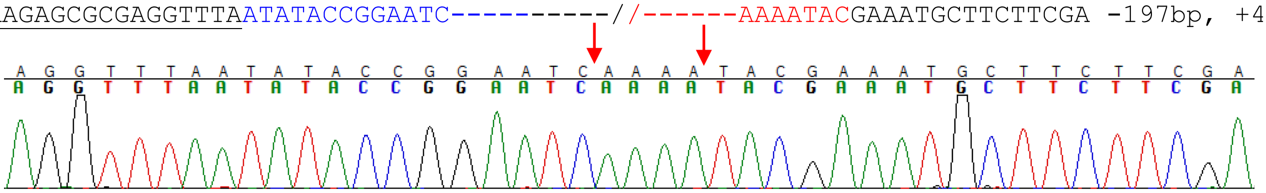

line 6, homozygous

AGCGCGAGGTTTAATAT-----------------//------CACTTGTACGAAATGCTTCTTCG -200 bp, homozygous

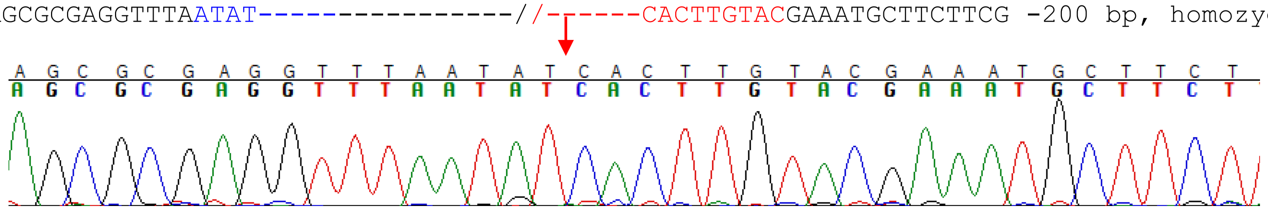

line 8, homozygous

CTAAAGGTTTGGAGAAGAAATCTACCTTGGGAGCTTCACTTGTACGAAATGCTTCTTCGAGAATCCGGCA SlRbhoD WT
CTAAAGGTTTGGAGAAGAAATCT-------GAGCTTCACTTGTACGAAATGCTTCTTCGAGAATCCGGCA line 8, -7bp

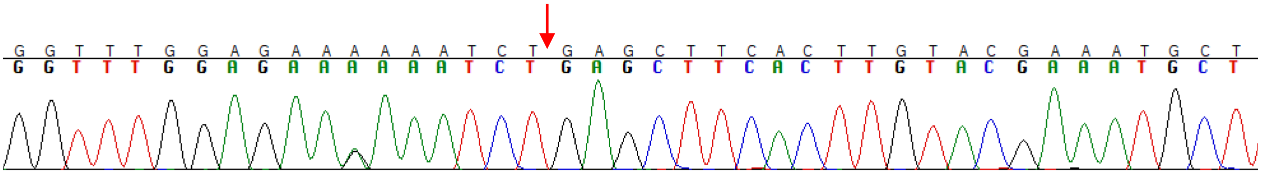

**Extended Data Figure S10.** Chromatograms for *SlRbohD* genotypes of tomato biallelic/homozygous mutants.

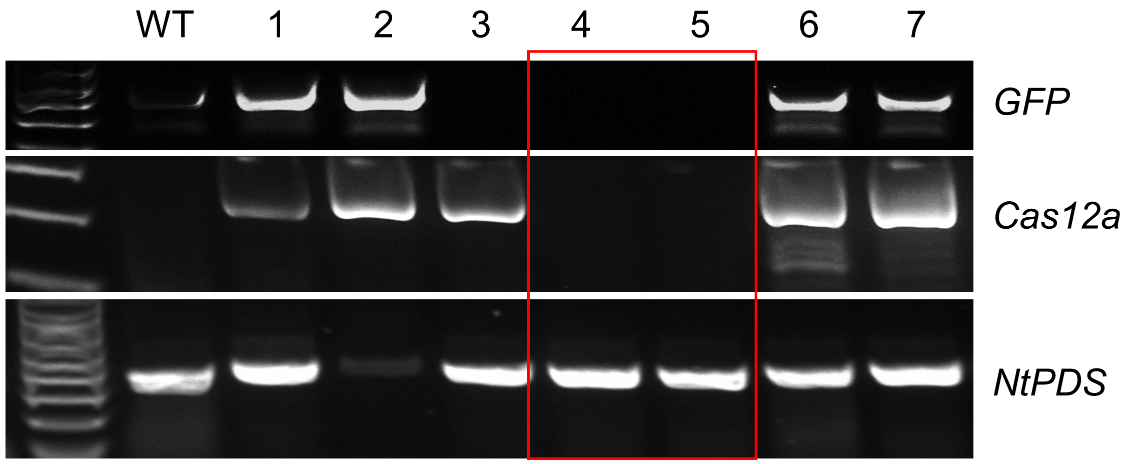


**Extended Data Figure S11. Confirmation of transgene-free, albino tobacco lines.** Detection of transgenes (*GFP* and *Cas12a*) in albino tobacco lines.

**a**

CAAGTACCACGTAGGATGATT**GG** *NtALS* WT
CAAGTATTATGTAGGATGATT**GG** *NtALS* mutant, transgene-free

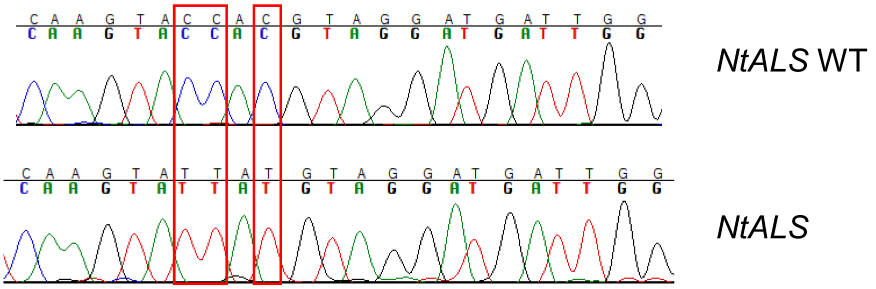


**b**

ATGAAATTTC**GGCAGATCAGAGCAAAGCAAAAA**TATT *NtPDS1* WT
ATGAAATTTC**GGCAGATCAGAGCAA--CAAAAA**TATT -2bp, line 4
ATGAAATTTC**GGCAGATCAGAGCAAAGCA----**TATT -4bp, line 4

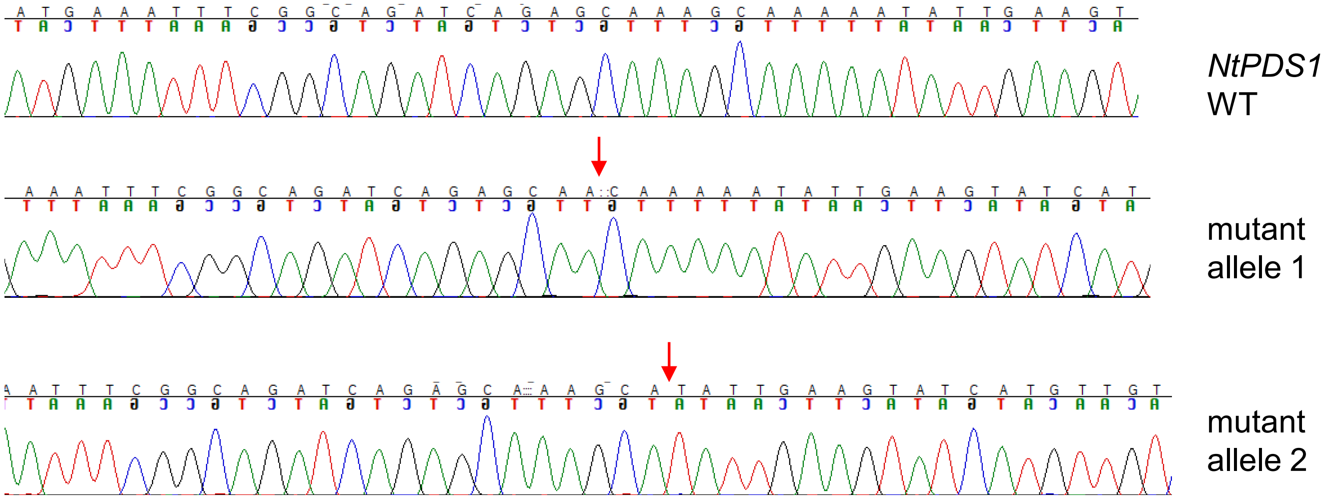


ACGAAATTTC**GGCAGATCAGAGCAAAGCAAAAA**TATTGAAGTATCACA NtPDS2 WT
ACGAAATTTC**GGCAGATCAGAGCAAAGCAAA----**TTGAAGTATCACA -4bp, line 4
ACGAAATTTC**GGCAGATCAGAGCAAA-------**TATTGAAGTATCACA -7bp, line 4

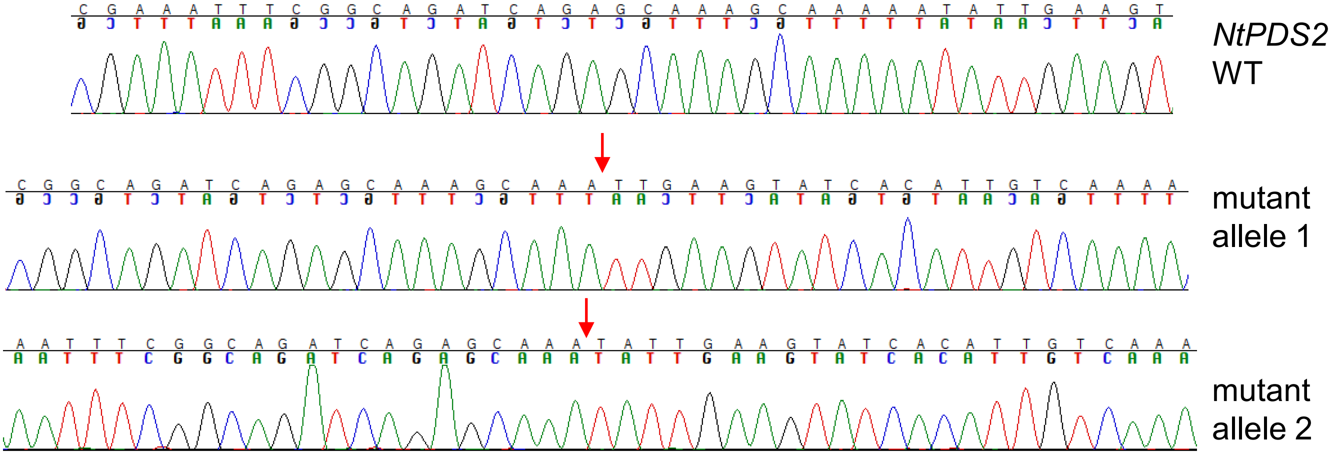


**Extended Data Figure S12. Genotyping of tobacco (*Nicotiana tabacum*) co-edited for *NtPDS* with *NtALS*. a,** Chromatogram for *NtALS* genotype of tobacco line 4. **b,** Chromatogram for *NtPDS genes* genotypes of tobacco line 4.

**a**

AGAAGAGAAGCT**AAAATTGTATTCAGATGATCCTT**CAAAGACCATGAG StDMR6 WT

AGAAGAGAAGCT**AAAA----ATTCAGATGATCCTT**CAAAGACCATGAG line 9, allele 1, -4bp
AGAAGAGAAGCT**AAAAT--------GATGATCCTT**CAAAGACCATGAG line 9, allele 2, -8bp
AGAAGAGAAGCT**AAAATTGTATTCAGATGATCCTT**CAAAGACCATGAG line 9, allele 3, WT
AGAAGAGAAGCT**AAAATTGTATTCAGATGATCCTT**CAAAGACCATGAG line 9, allele 4, WT

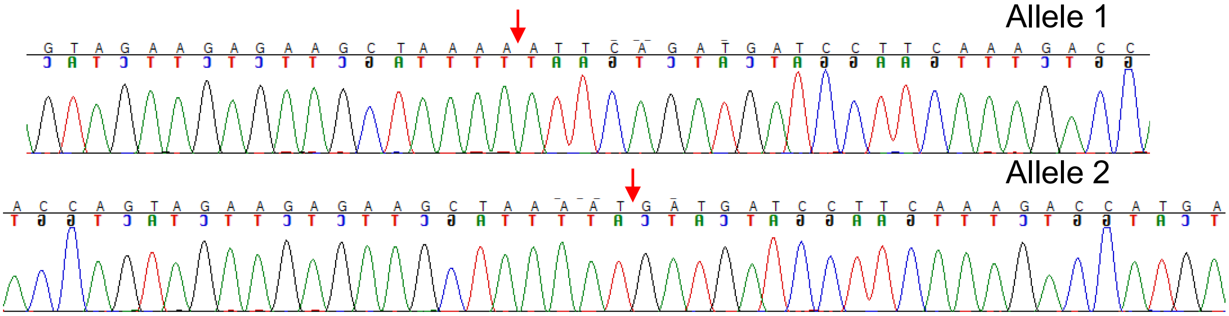


**Extended Data Figure S13. Genotyping of potato co-edited for *StDMR6* with *StALS.***
The chromatogram for *StDMR6* genotypes of potato mutant line #9 generated with 1 crRNA was shown. Only mutant alleles were shown.

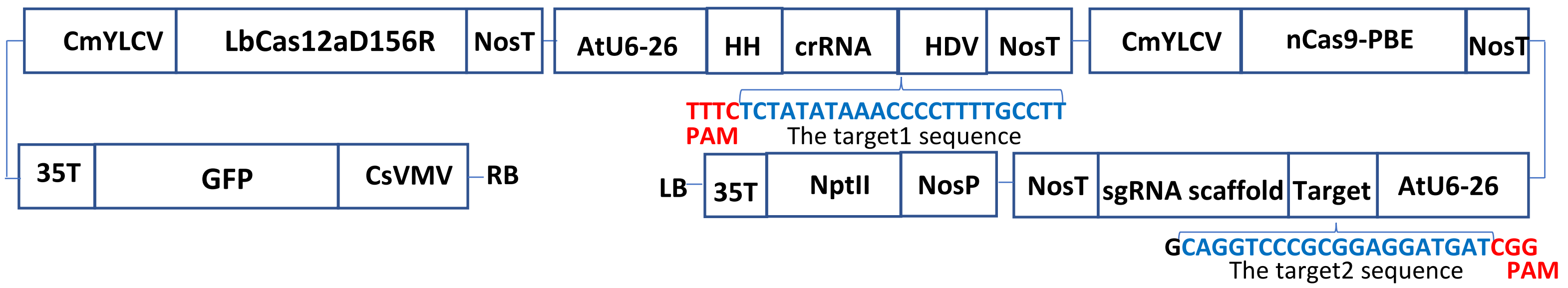


**Extended Data Figure S14.** Schematic representation of the binary vector GFP-p1380N-ttLbCas12a:LOBP1-PBE:ALS. LB and RB, the left and right borders of the T-DNA region; CsVMV, the cassava vein mosaic virus promoter; GFP, green fluorescent protein; 35T, the cauliflower mosaic virus 35S terminator; CmYLCV, the cestrum yellow leaf curling virus promoter; NosP and NosT, the nopaline synthase gene promoter and its terminator; ttLbCas12a, temperature-tolerant LbCas12a containing the single mutation D156R; AtU6-26, *Arabidopsis* U6-26 promoter; target1, the 23 nucleotides of Type II LOBP highlighted by blue, is located downstream of protospacer-adjacent motif (PAM); HH, the coding sequence of hammerhead ribozyme; HDV, the coding sequence of hepatitis delta virus ribozyme; nCas9-PBE, a plant base editor composed of rat cytidine deaminase APOBEC1, Cas9-D10A nickase (nCas9) and uracil glycosylase inhibitor (UGI); AtU6-26, *Arabidopsis* U6-26 promoter; target2, the 20 nucleotides of CsALS highlighted by blue, is located upstream of protospacer-adjacent motif (PAM); NptII, the coding sequence of neomycin phosphotransferase II.

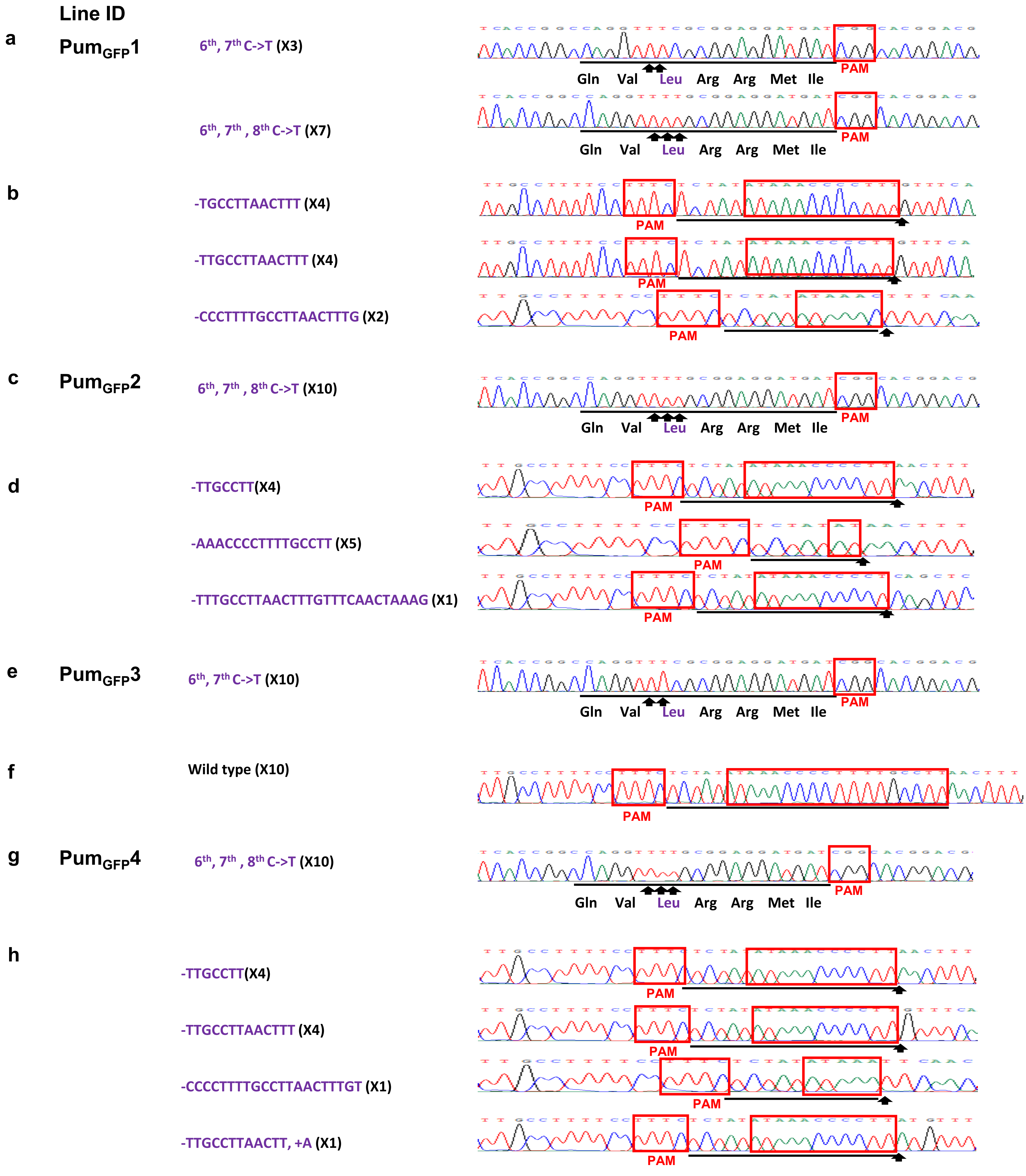


**Extended Data Figure S15.** Sanger sequencing analysis of transgenic pummelo (*Citrus maxima*) lines co-edited for *CsALS* and the effector binding elements in the promoter of the citrus canker susceptibility gene *CsLOB1* (EBE$_{PthA4}$-LOBP ). a, For *CsALS* of #Pum$_{GFP}$1, three of them are 6$^{th}$, 7$^{th}$ C->T conversions and seven of them are 6$^{th}$, 7$^{th}$, 8$^{th}$ C->T changes among 10 colonies sequenced. b, For EBE$_{PthA4}$-LOBP of #Pum$_{NoGFP}$1, there are four TGCCTTAACTTT, four TTGCCTTAACTTT and two CCCTTTTGCCTTAACTTTG deletion among 10 colonies sequenced. c, For *CsALS* of #Pum$_{GFP}$2, base editing occurred at 6$^{th}$, 7$^{th}$, 8$^{th}$ C->T among 10 colonies sequenced. d, For EBE$_{PthA4}$-LOBP of #Pum$_{NoGFP}$2, there are four TTGCCTT, five AAACCCCTTTTGCCTT and one TTTGCCTTAACTTTGTTTCAACTAAAG deletion among 10 colonies sequenced. e, For *CsALS* of #Pum$_{GFP}$3, all of them are 6$^{th}$, 7$^{th}$ C->T changes among 10 colonies sequenced. f, For EBE$_{PthA4}$-LOBP of #Pum$_{GFP}$3, all of them are wild type among 10 colonies sequenced. g, For *CsALS* of #Pum$_{GFP}$4, all of them are 6$^{th}$, 7$^{th}$ , 8$^{th}$ C->T conversions among 10 colonies sequenced. h, For EBE$_{PthA4}$-LOBP of #Pum$_{NoGFP}$4, there are four TTGCCTT, four TTGCCTTAACTTT, one CCCCTTTTGCCTTAACTTTGT and one TTGCCTTAACTT (containing one A insertion) deletion among 10 colonies sequenced.

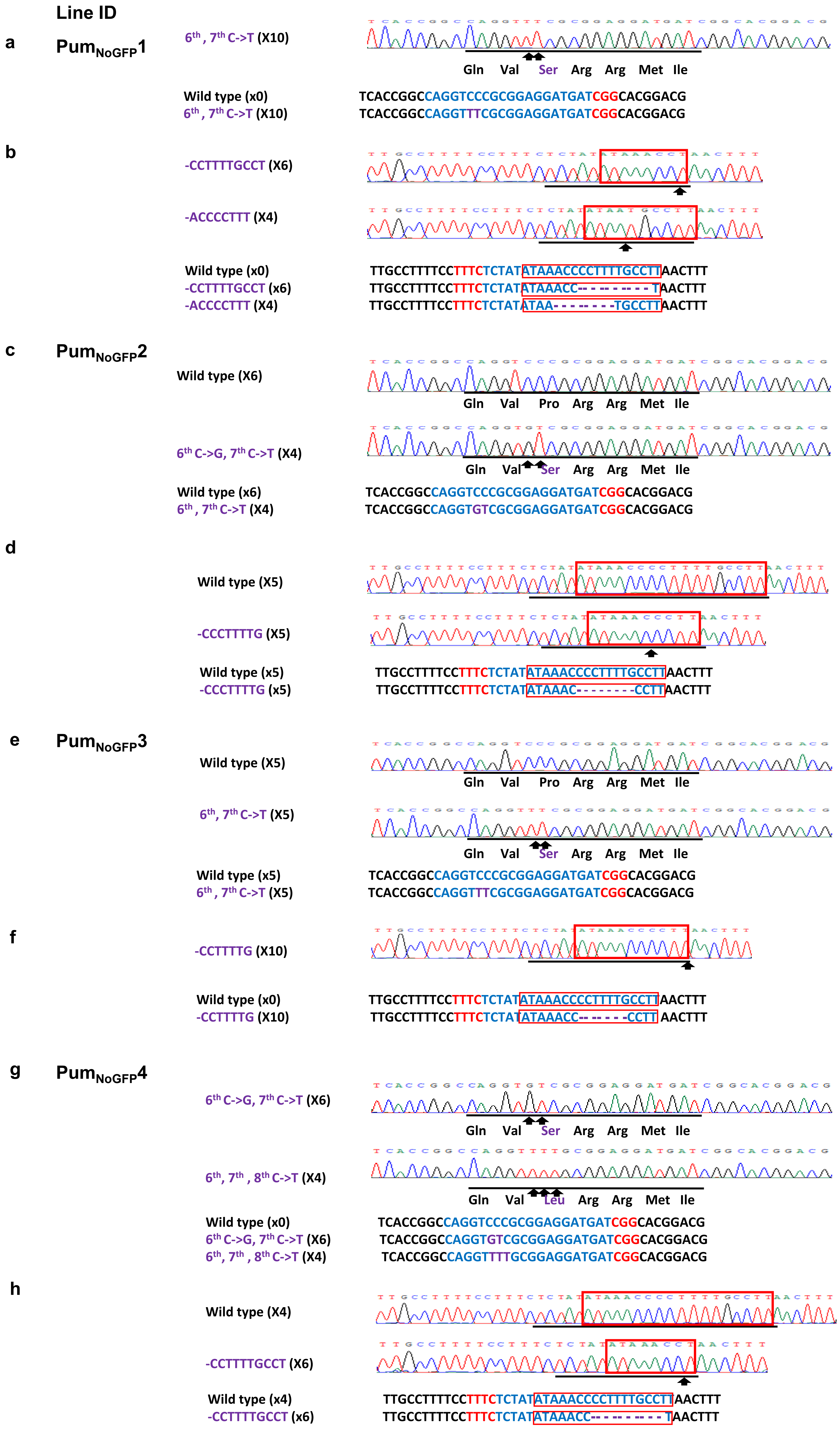


**Extended Data Figure S16.** Sanger sequencing analysis of transgene-free pummelo (*Citrus maxima*) lines co-edited for *CsALS* and the effector binding elements in the promoter of the citrus canker susceptibility gene *CsLOB1* (EBE$_{PthA4}$-LOBP ). a, For *CsALS* of #Pum$_{NoGFP}$1, all of them are 6$^{th}$, 7$^{th}$ C->T conversions among 10 colonies sequenced. b, For EBE$_{PthA4}$-LOBP of #Pum$_{NoGFP}$1, six of them are CCTTTTGCCT deletion, and four of them are ACCCCTTT deletion among 10 colonies sequenced. c, For *CsALS* of #Pum$_{NoGFP}$2, four colonies contain base editing (6$^{th}$ C->G, 7$^{th}$ C->T) among 10 colonies sequenced. d, For EBE$_{PthA4}$-LOBP of #Pum$_{NoGFP}$2, five of them are CCCTTTTG deletion among 10 colonies sequenced. e, For *CsALS* of #Pum$_{NoGFP}$3, five of them are 6$^{th}$, 7$^{th}$ C->T changes among 10 colonies sequenced. f, For EBE$_{PthA4}$-LOBP of #Pum$_{NoGFP}$3, all of them are CCTTTTG deletion among 10 colonies sequenced. g, For *CsALS* of #Pum$_{NoGFP}$4, six of them are 6$^{th}$ C->G, 7$^{th}$ C->T changes, and four of them are 6$^{th}$, 7$^{th}$ , 8$^{th}$ C->T conversions among 10 colonies sequenced. h, For EBE$_{PthA4}$-LOBP of #Pum$_{NoGFP}$4, six of them are CCTTTTGCCT deletion among 10 colonies sequenced.

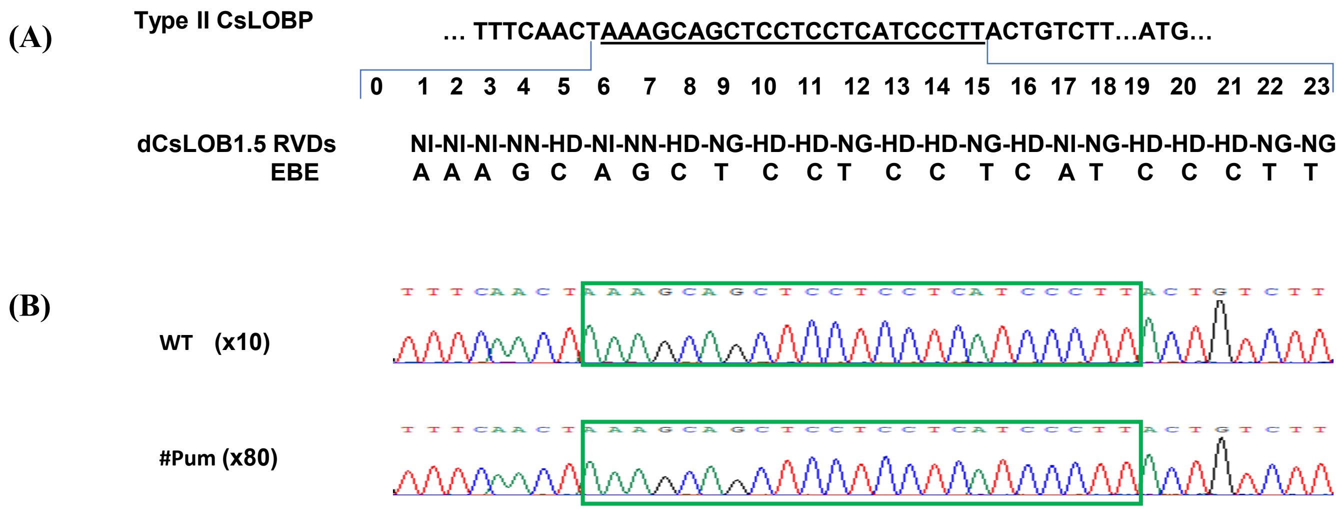


**Extended Data Figure S17.** dCsLOB1.5 and the representative chromatograms of dCsLOB1.5 targeting sequence in Pummelo. a, dCsLOB1.5 is an artificial dTALE, which specifically recognizes AAAGCAGCTCCTCCTCATCCCTT downstream of $EBE_{PthA4}$-LOBP by 7 bp. b, Representative chromatograms of dCsLOB1.5-binding sequence, which is the same in wild type, transgenic and transgene-free Pummelo plants. The dCsLOB1.5-binding sequence is highlighted by green rectangles.

GFP in the genome of sample EPGFP (transgenic control)

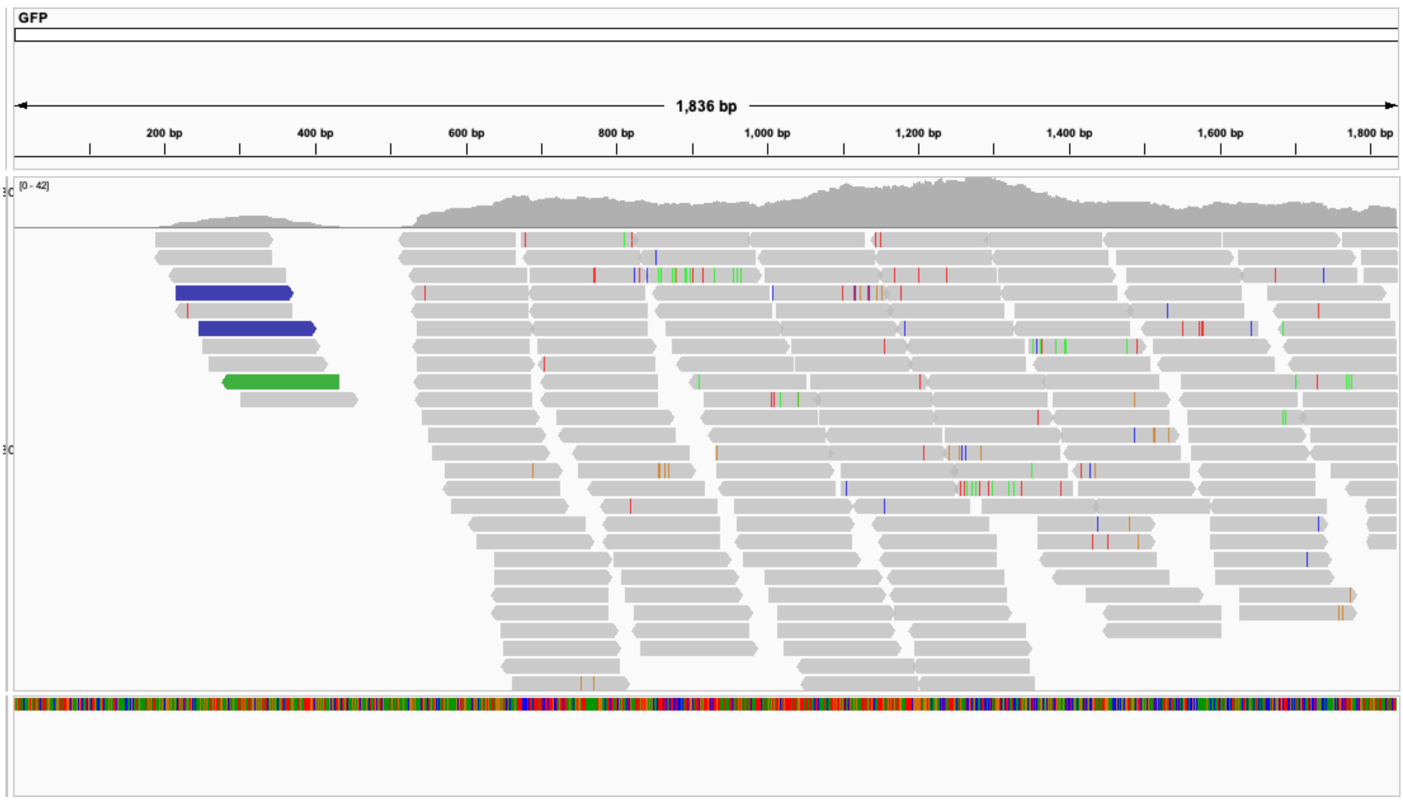


CsU6 promoter in the genome of sample EPGFP (transgenic control)

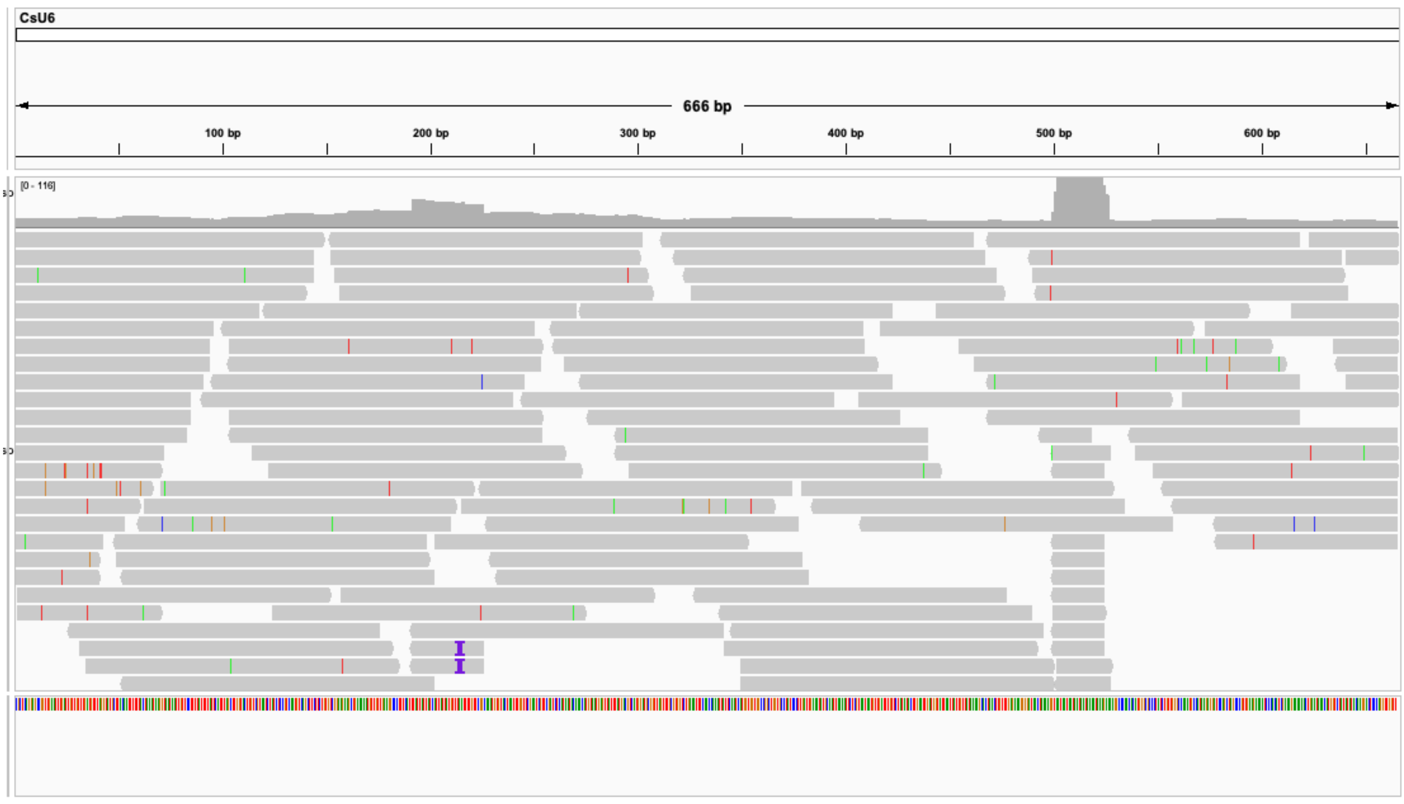


Cas12a in the genome of sample EPGFP (transgenic control)

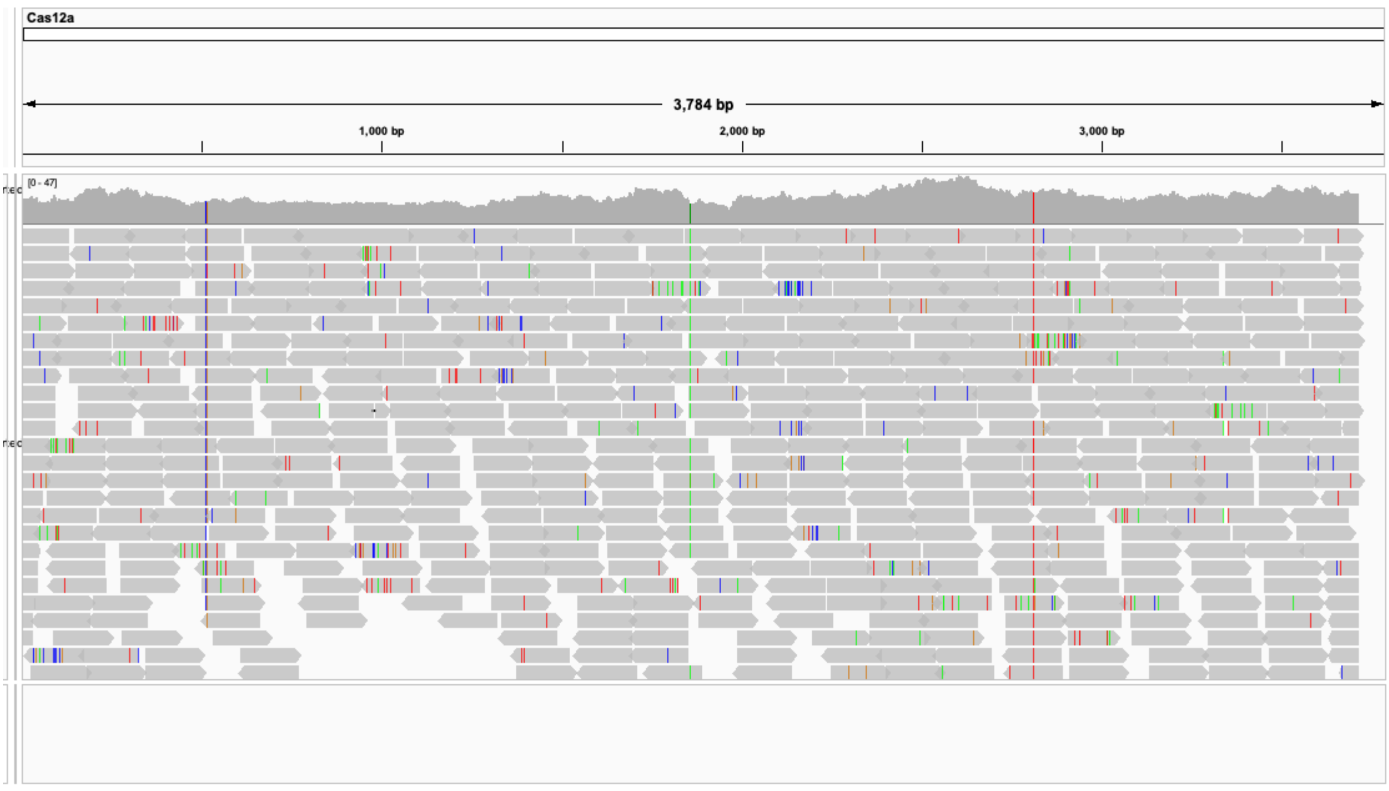


CBE in the genome of sample EPGFP (transgenic control)

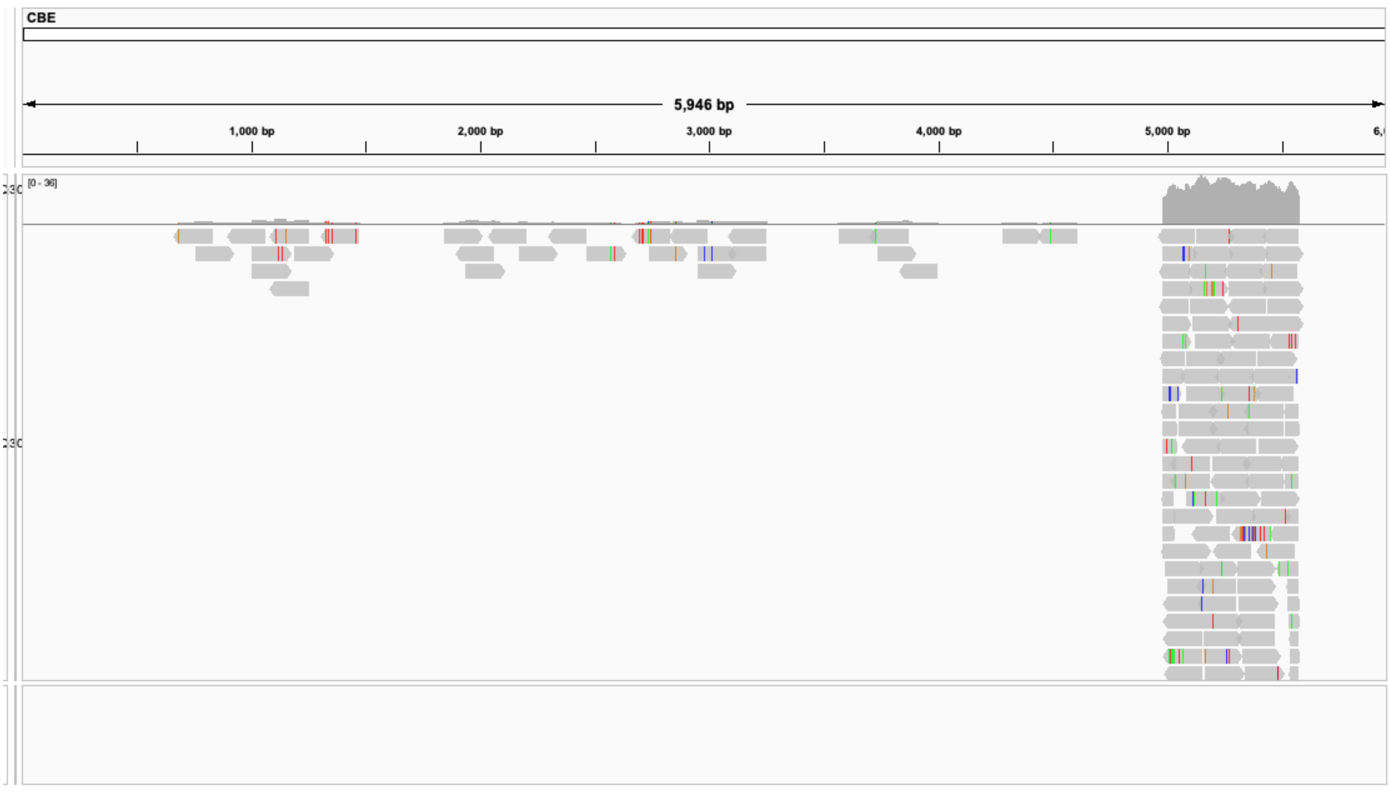


**Extended Data Figure S18.** Mapping of construct DNA in the transgenic tomato line EPGFP using whole genome sequencing data.

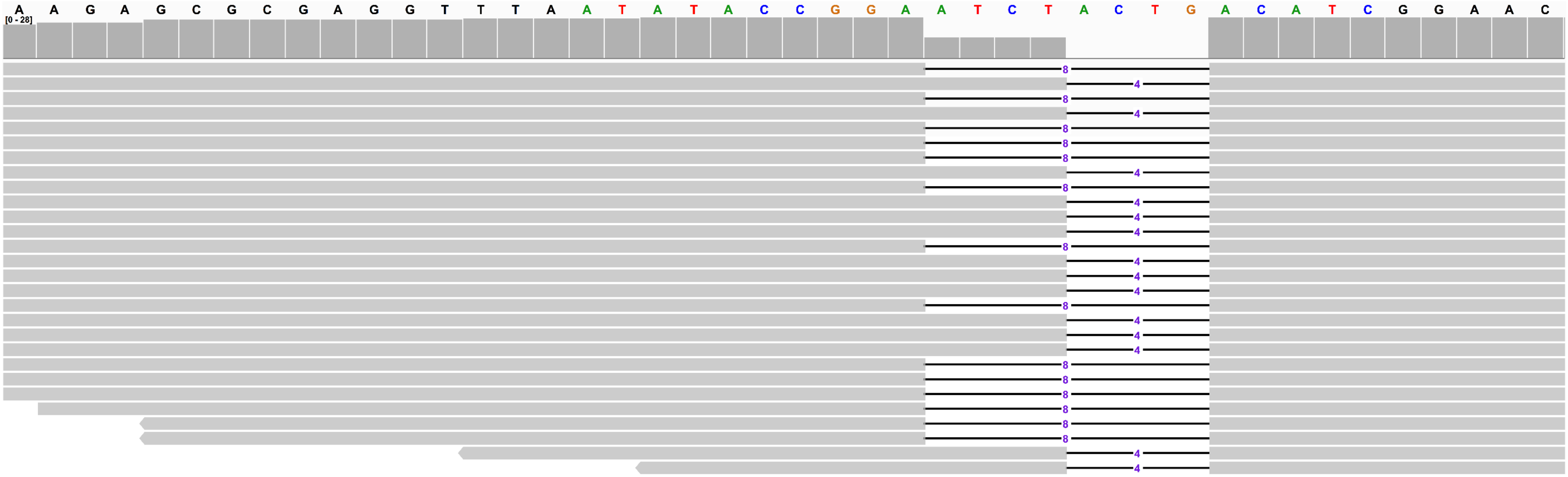


**Extended Data Figure S19.** Mapping of *SlRbohD* in the *slrbohd* line 1 using whole genome sequencing data. The nucleotides of target site were highlighted by colors other than black. There were two types of deletions of target site, including type I (-8 bp deletion) and Type II (-4 bp deletion), which were shown by horizontal bar chart. The by vertical bar chart showed the sequence depth for each base.

AAAGGTTTGGAGAAGAAATCTACCTTGGGAGCTTCACTTGTACGAAATGCTTC *SlRbohD* WT

AAAGGTTTGGAGAAGAAATCT-------GAGCTTCACTTGTACGAAATGCTTC line 8, -7bp

***SlRbohd* in line 8**

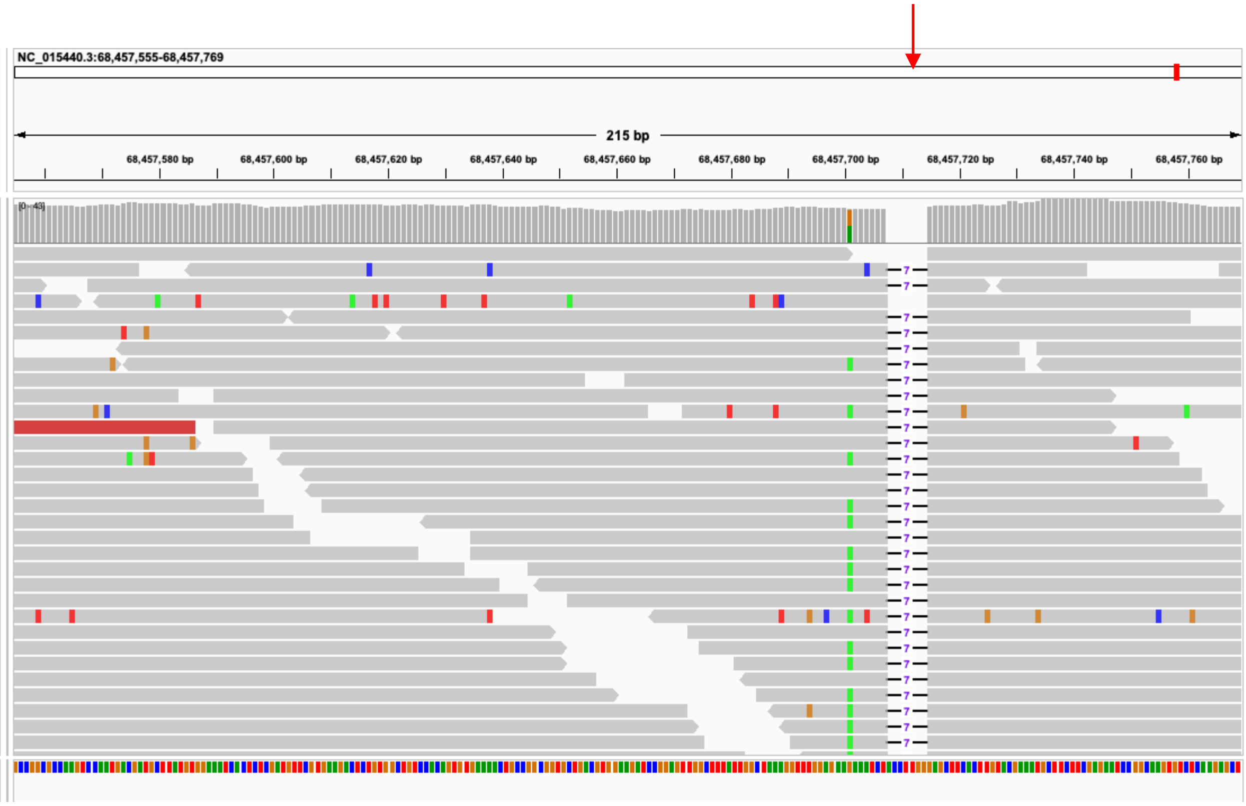


**Extended Data Figure S20.** Mapping of *SlRbohD* in the *slrbohd-8* line using whole genome sequencing data.

URL of this page: http://www.rgenome.net/cas-offinder/result?hash=e64e267b51b9d53d628b14f51a6001d2

| Job ID | Title | Submit Date | End Date | Status |
|---|---|---|---|---|
| 566940 | Untitled | Jan. 23, 2023, 8:40 p.m. | Jan. 23, 2023, 8:40 p.m. | Finished! Download result (or old-style result) |

### Summary

| Target Sequence | Bulge Type | Bulge Size | Mismatch | Number of Found Targets |
|---|---|---|---|---|
| TTTVTCTATATAAACCCCTTTTGCCTT | X | 0 | 0 | 1 |
| TTTVTCTATATAAACCCCTTTTGCCTT | X | 0 | 4 | 1 |
| TTTVTCTATATAAACCCCTTTTGCCTT | X | 0 | 5 | 7 |

### Details

Bulge Type | DNA bulge | RNA bulge | Mismatch | Filter | Download filtered result

| Bulge Type | Target | Chromosome | Position | Direction | Mismatches | Bulge Size |
|---|---|---|---|---|---|---|
| X | crRNA: TTTVTCTATATAAACCCCTTTTGCCTT<br>DNA: TTTATCTAgtgAAAaCCCTTTTGCCTT | NC_023046.1 | 22039254 | + | 4 | 0 |
| X | crRNA: TTTVTCTATATAAACCCCTTTTGCCTT<br>DNA: TTTCTaTATATAAtCCCCTTcgGCCaT | NC_023046.1 | 24273597 | - | 5 | 0 |
| X | crRNA: TTTVTCTATATAAACCCCTTTTGCCTT<br>DNA: TTTCTCaATAaAAAagCCTTTgGCCTT | NC_023047.1 | 11310893 | + | 5 | 0 |
| X | crRNA: TTTVTCTATATAAACCCCTTTTGCCTT<br>DNA: TTTCTtTAaATAAACCaCTTTgcCCTT | NC_023048.1 | 16426331 | + | 5 | 0 |
| X | crRNA: TTTVTCTATATAAACCCCTTTTGCCTT<br>DNA: TTTCTtTAaATAAACCaCTTTgcCCTT | NC_023048.1 | 16427432 | + | 5 | 0 |
| X | crRNA: TTTVTCTATATAAACCCCTTTTGCCTT<br>DNA: TTTATtTAaATAtACCCCaTTTGtCTT | NC_023050.1 | 9869312 | + | 5 | 0 |
| X | crRNA: TTTVTCTATATAAACCCCTTTTGCCTT<br>DNA: TTTGTCcATATAAACCCtTTTTtttTT | NC_023052.1 | 2436441 | - | 5 | 0 |
| X | crRNA: TTTVTCTATATAAACCCCTTTTGCCTT<br>DNA: TTTCTCTATATAAACCCCTTTTGCCTT | NC_023052.1 | 28359392 | - | 0 | 0 |
| X | crRNA: TTTVTCTATATAAACCCCTTTTGCCTT<br>DNA: TTTCTCaATtTAAACCgtTgTTGCCTT | NW_006257128.1 | 313631 | - | 5 | 0 |

**Extended Data Figure S21.** Eight potential off-targets identified for the crRNA targeting $EBE_{PthA4}$-LOBP. Off-target analysis was conducted using a web software (http://www.rgenome.net/cas-offinder/). Eight potential off-targets were identified with up to 5 bp mismatches with the targeting crRNA.

**PUC57-mini-crRNA sequence**

CmYLCV promoter,**HH**,direct repeat,**HDV,** HSP18.2 terminator, Nos terminator

TGGCAGACATACTGTCCCACAAATGAAGATGGAATCTGTAAAAGAAAACGCGTGAAATAATGCG
TCTGACAAAGGTTAGGTCGGCTGCCTTTAATCAATACCAAAGTGGTCCCTACCACGATGGAAAA
ACTGTGCAGTCGGTTTGGCTTTTTCTGACGAACAAATAAGATTCGTGGCCGACAGGTGGGGGTC
CACCATGTGAAGGCATCTTCAGACTCCAATAATGGAGCAATGACGTAAGGGCTTACGAAATAAG
TAAGGGTAGTTTGGGAAATGTCCACTCACCCGTCAGTCTATAAATACTTAGCCCCTCCCTCATT
GTTAAGGGAGCAAAATCTCAGAGAGATAGTCCTAGAGAGAGAAAGAGAGCAAGTAGCCTAGAAG
TAGTCAAGGCGGCGAAGTATTCAGGCACGTGGCCAGGAAGAAGAAAAGCCAAGACGACGAAAAC
AGGTAAGAGCTAAGCTTCTGCAG**AAATTACTGATGAGTCCGTGAGGACGAAACGAGTAAGCTCG**
**TC**TAATTTCTACTAAGTGTAGATGAGACGGAGCTCAGTCTGACCGCGGCGTCTCT**GGCCGGCAT**
**GGTCCCAGCCTCCTCGCTGGCGCCGGCTGGGCAACATGCTTCGGCATGGCGAATGGGAC**TTTTT
TTTATATGAAGATGAAGATGAAATATTTGGTGTGTCAAATAAAAAGCTTGTGTGCTTAAGTTTG
TGTTTTTTTCTTGGCTTGTTGTGTTATGAATTTGTGGCTTTTTCTAATATTAAATGAATGTAAG
ATCACATTATAATGAATAAACAAATGTTTCTATAATCCATTGTGAATGTTTTGTTGGATCTCTT
CTGCAGCATATAACTACTGTATGTGCTATGGTATGGACTATGGAATATGATTAAAGATAAGAGC
CTCGAATTTCCCCGATCGTTCAAACATTTGGCAATAAAGTTTCTTAAGATTGAATCCTGTTGCC
GGTCTTGCGATGATTATCATATAATTTCTGTTGAATTACGTTAAGCATGTAATAATTAACATGT
AATGCATGACGTTATTTATGAGATGGGTTTTTATGATTAGAGTCCCGCAATTATACATTTAATA
CGCGATAGAAAACAAAATATAGCGCGCAAACTAGGATAAATTATCGCGCGCGGTGTCATCTATG
TTACTAGATCGATATCATTGG

**PUC57-HDV-HH-DR**

**HDV, HH**, direct repeat (DR),

**GGCCGGCATGGTCCCAGCCTCCTCGCTGGCGCCGGCTGGGCAACATGCTTCGGCATGGCGAATG**
**GGACAAATTACTGATGAGTCCGTGAGGACGAAACGAGTAAGCTCGTC**TAATTTCTACTAAGTGT
AGAT

**Extended Data Fig. S22.** PUC57-mini-crRNA and PUC57-HDV-HH-DR sequences were shown.

Supplementary Information File 1. Chromatograms for *SlRbohD* and *SlPPAD4* edited GFP-negative biallelic/homozygous tomato mutants.

Target: *SlEDS1* and *SlPPAD4*
Line 5, *SlEDS1* biallelic, *SlPAD4* WT

```
GGTACAGTAACACTTCTTTTGGAGAGAAAGAGATCAACACCACTCTGTTTCCATCTTTGAGAAGTG  SlEDS1 WT
GGTACAGTAACACTTCTTTTGGAGAGAAAGAGATCAAC--CACTCTGTTTCCATCTTTGAGAAGTG  line 5, allele 1, -2bp
GGTACAGTAA----------------------------------CTGTTTCCATCTTTGAGAAGTG  line 5, allele 2, -34bp
```

*sleds1*, line 5, allele 1

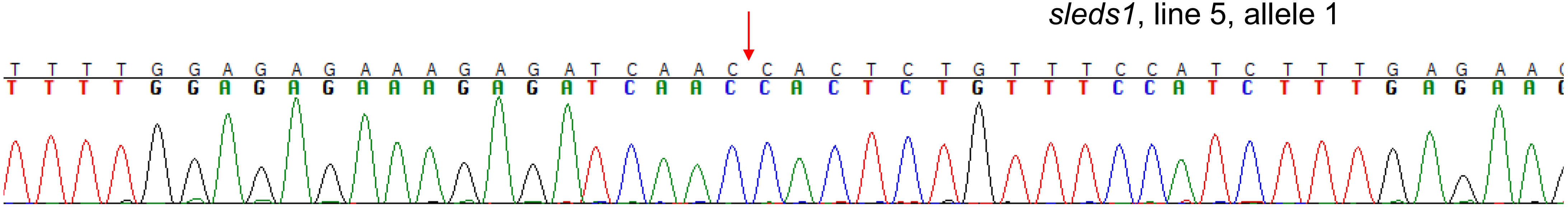


*sleds1*, line 5, allele 2

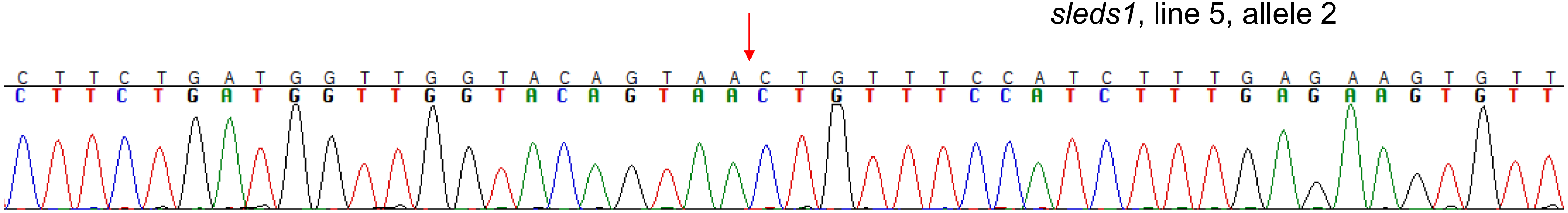

Line 6, *SlEDS1* biallelic, *SlPAD4* biallelic

```
GGTACAGTAACACTTCTTTTGGAGAGAAAGAGATCAACACCACTCTGTTTCCATCTTTGAGAAGTG  SlEDS1 WT
GGTACAGTAACACTTCTTTTGGAGAGAAAGAGATCAAC-----TCTGTTTCCATCTTTGAGAAGTG  line 6, allele 1, -5bp
GGTACAGTAACACTTCTTTTGGAGAGAAAGAGATC----------TGTTTCCATCTTTGAGAAGTG  line 6, allele 2, -10bp
```

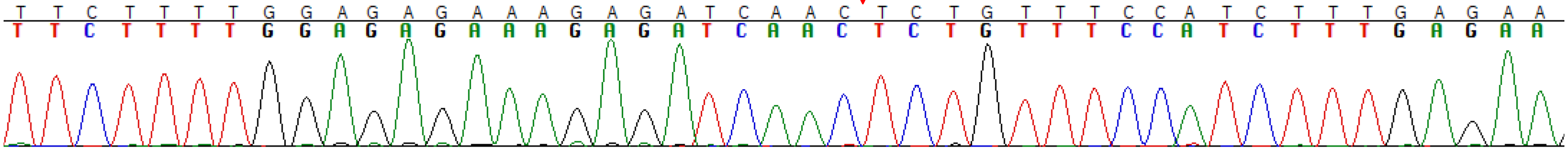

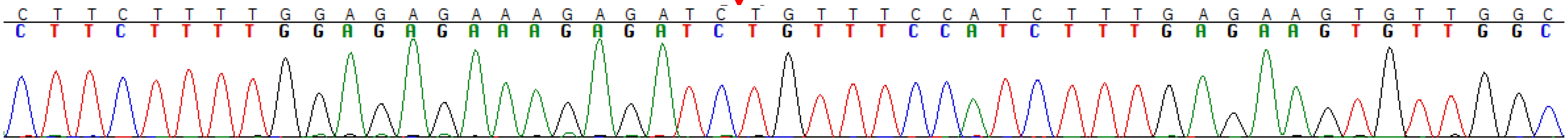

```
CAATTTCGCCGTCAATCGAGTTGGTGAGACAGCTTATGTGGGATTCTCCGGCGTGAAATTGGGCGCCGGAGTGGA SlPAD4 WT
CAATTTCGCCGTCAATCGAGTTGGTGAGACA--------GGGATTCTCCGGCGTGAAATTGGGCGCCGGAGTGGA line 6, allele 1, -8bp
CAATTTCGCCGTCAATCGAGTTGGTGAGACAG-----GTGGGATTCTCCGGCGTGAAATTGGGCGCCGGAGTGGA line 6, allele 2, -5bp
```

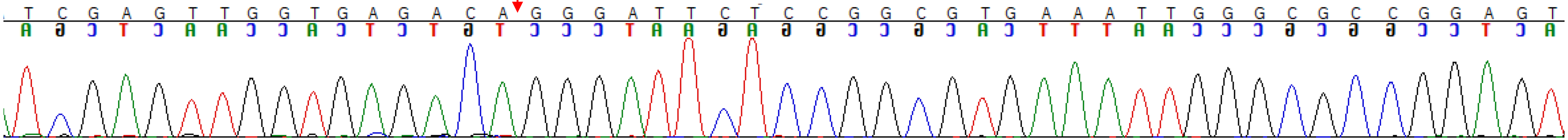

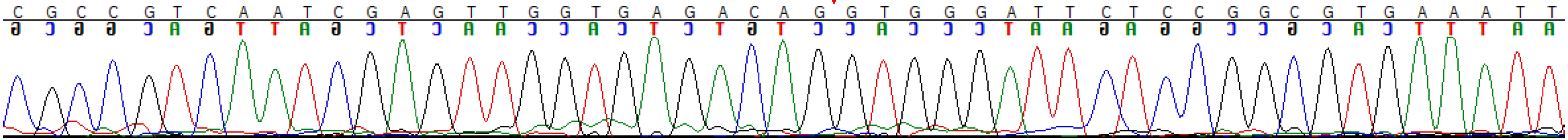

Line 8, *SlEDS1* homozygous, *SlPAD4* homozygous

GGTACAGTAACACTTCTTTTGGAGAGAAAGAGATCAACACCACTCTGTTTCCATCTTTGAGAAGTG *SlEDS1* WT
GGTACAGTAACACTTCTTTTGGAGAGAAAGAGATCAAC-----TCTGTTTCCATCTTTGAGAAGTG line 8, -5bp, homozygous

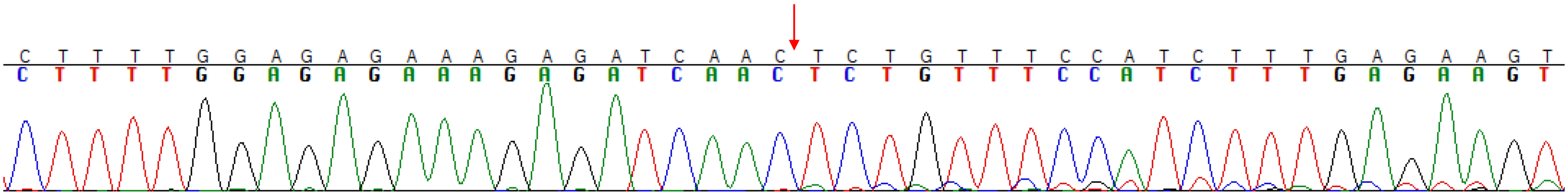

TTTCGCCGTCAATCGAGTTGGTGAGACAGCTTATGTGGGATTCTCCGGCGTGAAATTGGGCGCCGGAGTGGACCAA *SlPAD4* WT
TTTCGCCGTCAATCGAGTTGGTGAGACAG--------GGATTCTCCGGCGTGAAATTGGGCGCCGGAGTGGACCAA line 8, -8bp, homozygous

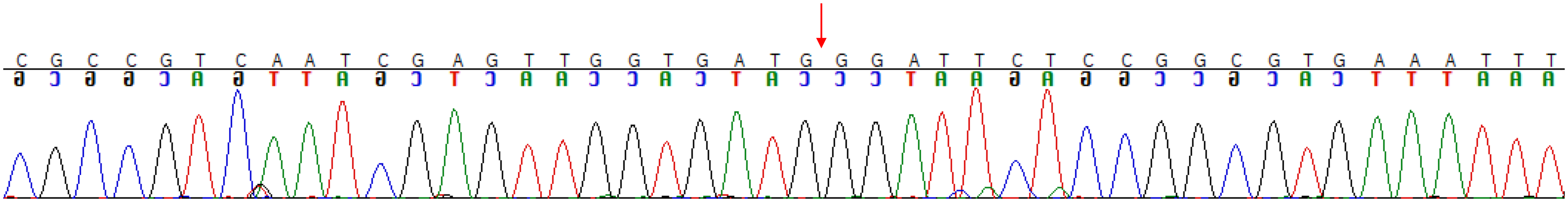

Line 10, *SlEDS1* biallelic, SlPAD4 WT

```
GGTACAGTAACACTTCTTTTGGAGAGAAAGAGATCAACACCACTCTGTTTCCATCTTTGAGAAGTG   SlEDS1 WT
GGTACAGTAACACTTCTTTTGGAGAGAAAGAGATCAACAC-----TGTTTCCATCTTTGAGAAGTG   line 10, -5bp
GGTACAGTAACACTTCTTTTGGAGAGAAAGAGATCAACACC---CTGTTTCCATCTTTGAGAAGTG   line 10, -3bp
```

*sleds1*, line 10, allele 1

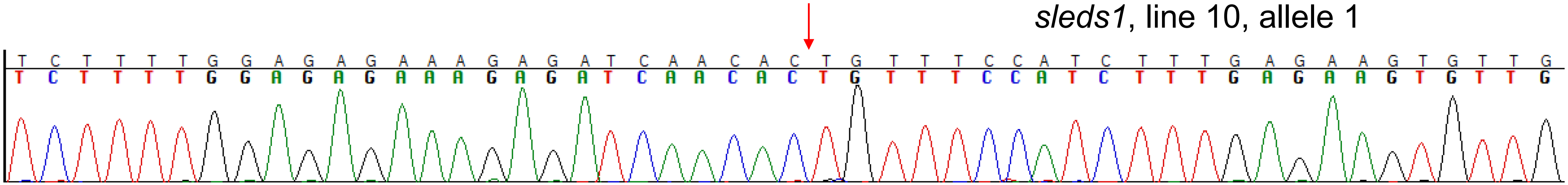


*sleds1*, line 10, allele 2

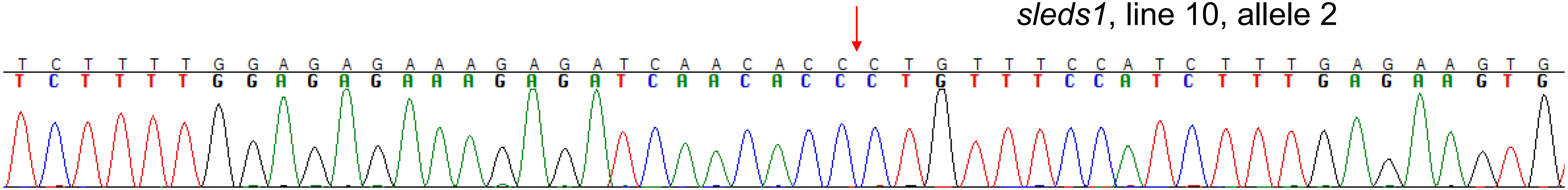

### Line 11, *SlEDS1* biallelic, *SlPAD4* biallelic

GGTACAGTAACACTTCTTTTGGAGAGAAAGAGATCAACACCACTCTGTTTCCATCTTTGAGAAGTG *SlEDS1* WT
GGTACAGTAACACTTCTTTTGGAGAGAAAGA-------------CTGTTTCCATCTTTGAGAAGTG line 11 allele 1, -13bp
GGTACAGTAACACTTCTTTTGGAGAGAAAGAGATC-----CACTCTGTTTCCATCTTTGAGAAGTG line 11 allele 2, -5bp

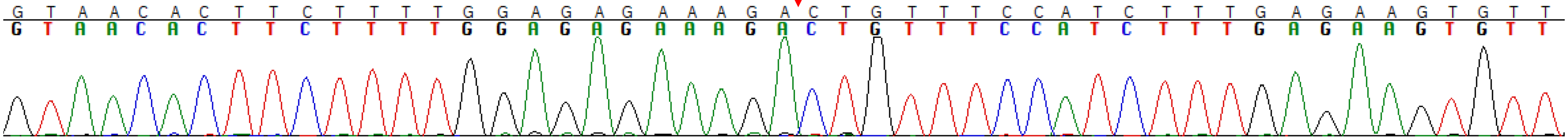


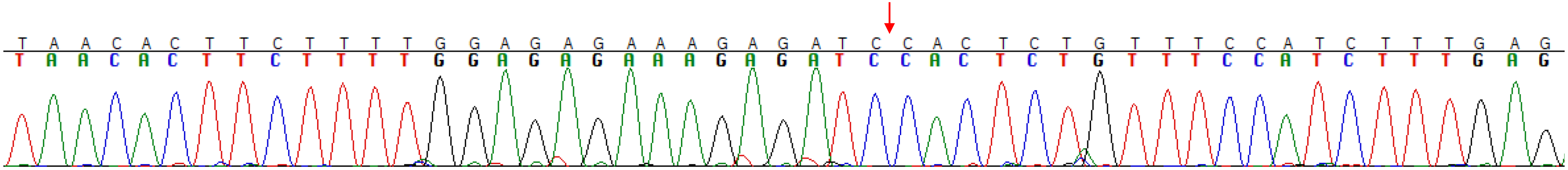


TTTCGCCGTCAATCGAGTTGGTGAGACAGCTTATGTGGGATTCTCCGGCGTGAAATTGGGCGCCGGAGTGGACCAA *SlPAD4* WT
TTTCGCCGTCAATCGAGTTGGTGAGACAG------------TCTCCGGCGTGAAATTGGGCGCCGGAGTGGACCAA line 11, -12bp
TTTCGCCGTCAATCGAGTTGGTGAGA---------TGGGATTCTCCGGCGTGAAATTGGGCGCCGGAGTGGACCAA line 11, -9bp

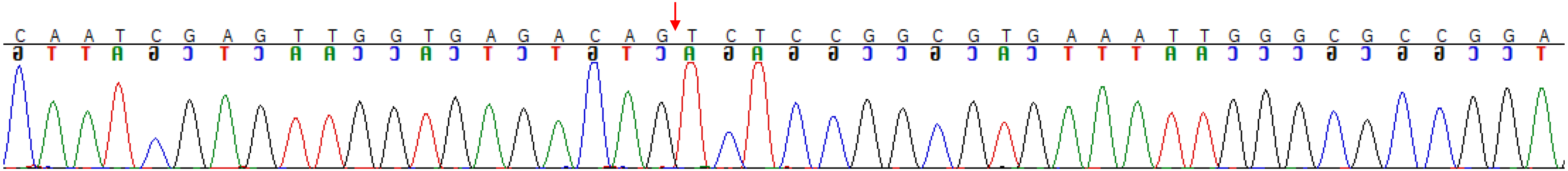


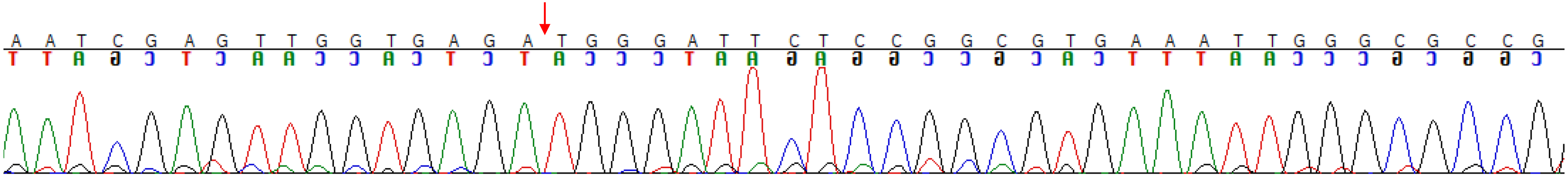

Line 14, *SlEDS1* biallelic, *SlPAD4* biallelic

GGTACAGTAACACTTCTTTTGGAGAGAAAGAGATCAACACCACTCTGTTTCCATCTTTGAGAAGTG *SlEDS1* WT
GGTACAGTAACACTTCTTTTGGAGAGAAAGAGATC------------------TCTTTGAGAAGTG line 14 allele 1, -18bp
GGTACAGTAACACTTCTTTTGGAGAGAAAGAGATCAAC-----TCTGTTTCCATCTTTGAGAAGTG line 14 allele 2, -5bp

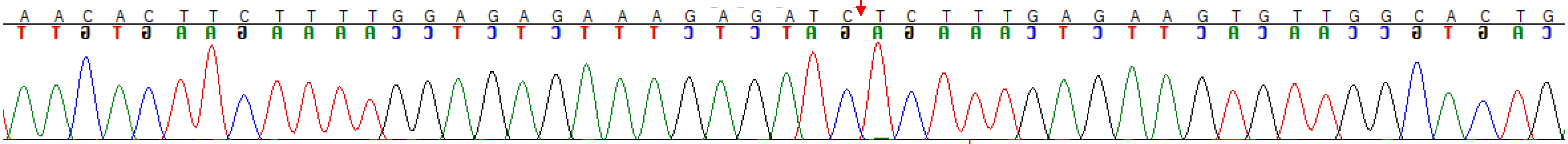

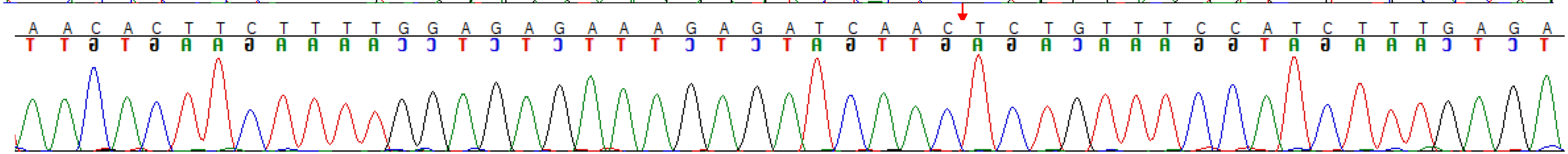

TTTCGCCGTCAATCGAGTTGGTGAGACAGCTTATGTGGGATTCTCCGGCGTGAAATTGGGCGCCGGAGTGGACCAA *SlPAD4* WT
TTTCGCCGTCAATCGAGTT---------------------TTCTCCGGCGTGAAATTGGGCGCCGGAGTGGACCAA line 14, -21bp
TTTCGCCGTCAATCGAGTTGGTGAGACA-----TGTGGGATTCTCCGGCGTGAAATTGGGCGCCGGAGTGGACCAA line 14, -5bp

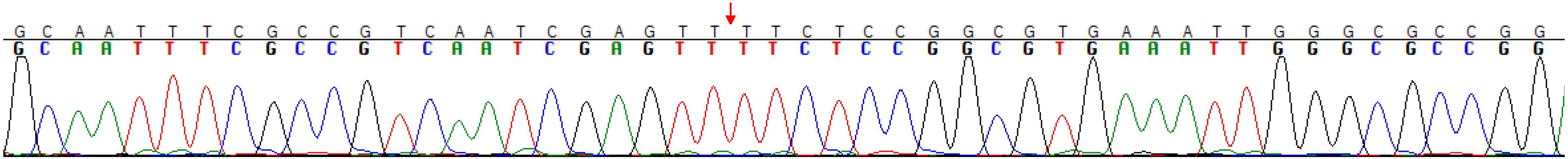

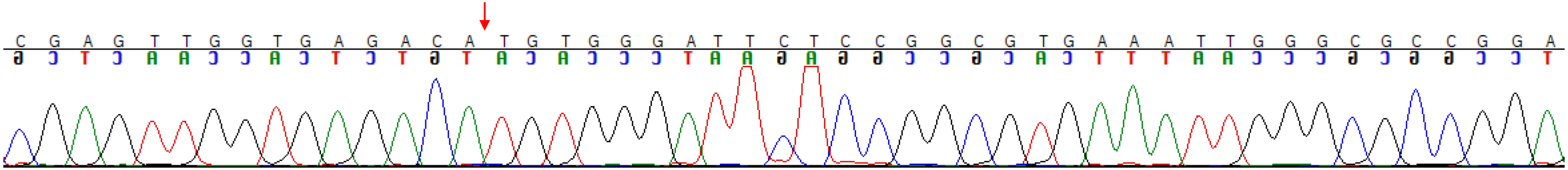

Line 16, *SlEDS1* biallelic, *SlPAD4* homozygous

GGTACAGTAACACTTCTTTTGGAGAGAAAGAGATCAACACCACTCTGTTTCCATCTTTGAGAAGTG *SlEDS1* WT
GGTACAGTAACACTTCTTTTGGAGAGAAAGAGATCAACAC-------TTTCCATCTTTGAGAAGTG line 16, -7bp
GGTACAGTAACACTTCTTTTGGAGAGAAAGAGATC------ACTCTGTTTCCATCTTTGAGAAGTG line 16, -6bp

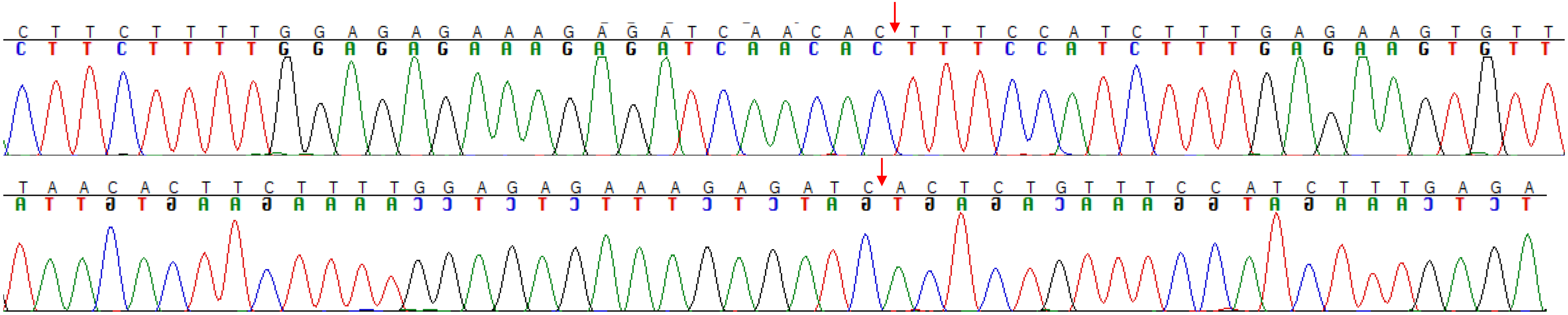

TTTCGCCGTCAATCGAGTTGGTGAGACAGCTTATGTGGGATTCTCCGGCGTGAAATTGGGCGCCGGAGTGGACCAA *SlPAD4* WT
TTTCGCCGTCAATCGAGTTGGTGAGACAGCTTA----GGATTCTCCGGCGTGAAATTGGGCGCCGGAGTGGACCAA line 16, -4bp, homozygous

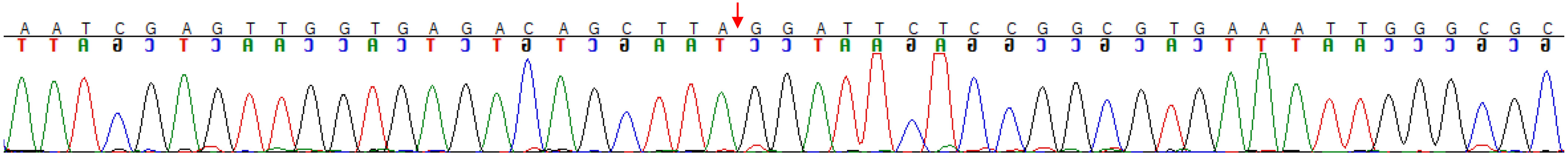

Line 18, *SlEDS1* biallelic, *SlPAD4* biallelic

```
GGTACAGTAACACTTCTTTTGGAGAGAAAGAGATCAACACCACTCTGTTTCCATCTTTGAGAAGTG  SlEDS1 WT
GGTACAGTAACACTTCTTTTGGAGAGAAAGAGATCA-------TCTGTTTCCATCTTTGAGAAGTG  line 18 allele 1, -7bp
GGTACAGTAACACTTCTTTTGGAGAGAAAGAGATC----------TGTTTCCATCTTTGAGAAGTG  line 18 allele 2, -10bp
```

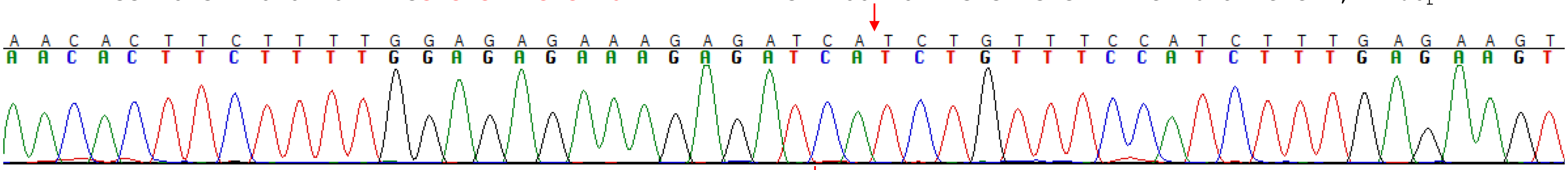

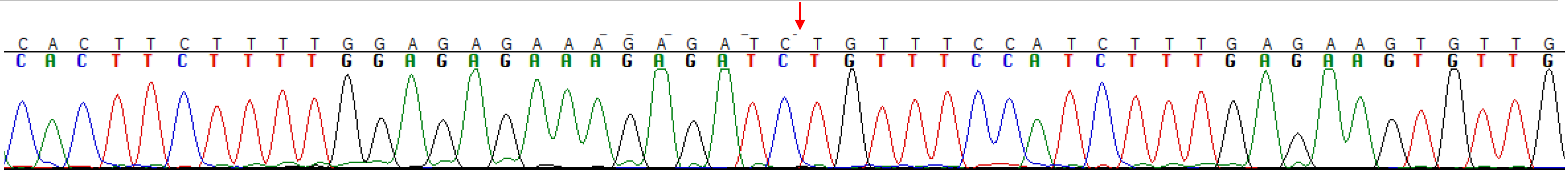

```
TCGGTGGGTTGCAATTTCGCCGTCAATCGAGTTGGTGAGACAGCTTATGTGGGATTCTCCGGCGTGAAATTGGGCGCCGGAGTGGACCAAAG SlPAD4 WT
TCGGTGGGTTGCAATTTCGCCGTC-----------------------------------CGGCGTGAAATTGGGCGCCGGAGTGGACCAAAG line 18, -35bp
TCGGTGGGTTGCAATTTCGCCGTCAATCGAGTTGGTGAGACAGC------GGGATTCTCCGGCGTGAAATTGGGCGCCGGAGTGGACCAAAG line 18, -6bp
```

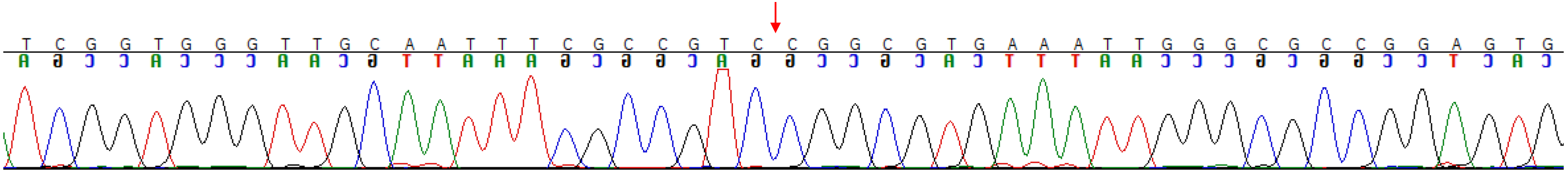

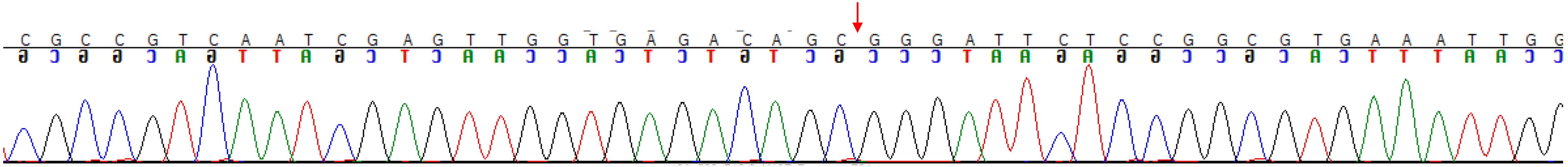

Supplementary Information File 2. Chromatograms for *SlDMR6* and *SlINVINH1* edited GFP-negative biallelic/homozygous tomato mutants.

Target: *SlDMR6* and *SlINVINH1*

line3, *SlDMR6* biallelic, *SlINVINH1* biallelic

ACCCGAATCCGATAGACCAC**GTCTATCGGAAGTGGTCGATTGT**GAAAATGTTCCAATAATTGACTTAAGTTGCGGAGATCAAGCT SlDMR6 WT

ACCCGAATCCGATAGACCAC**G-------GAAGTGGT-------**---------------------------TGCGGAGATCAAGCT line 3,allele 1,-41bp

ACCCGAATCCGATAGACCAC**G---ATA-GAAGTGGTCGATTGT**GAAAATGTTCCAATAATTGACTTAAGTTGCGGAGATCAAGCT line 3,allele 2, -7bp,+3bp

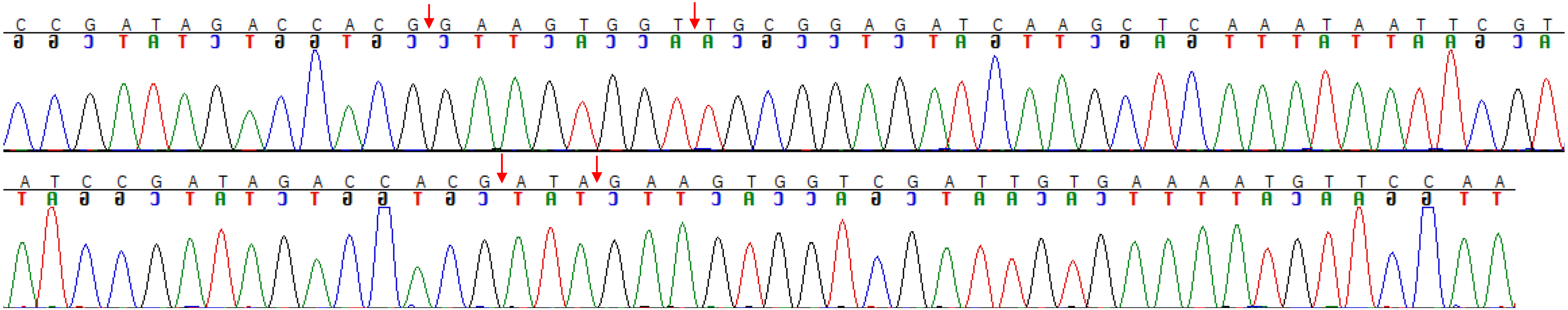

TAGTAACAAGTGGGAATAATAATCTAGT**AGAGACAACATGCAAGAACACAC**CAAATTATAATTTGTGTGTGAAAACTTTGTCT *SlINVINH1* WT

TAGTAACAAGTGGGAATA----------**---------ATGCAAGAACACAC**CAAATTATAATTTGTGTGTGAAAACTTTGTCT line 3,allele 1,-19bp

TAGTAACAAGTGGGAATAATAATCTAGT**AGA----------AAGAACACAC**CAAATTATAATTTGTGTGTGAAAACTTTGTCT line 3,allele 2,-10bp

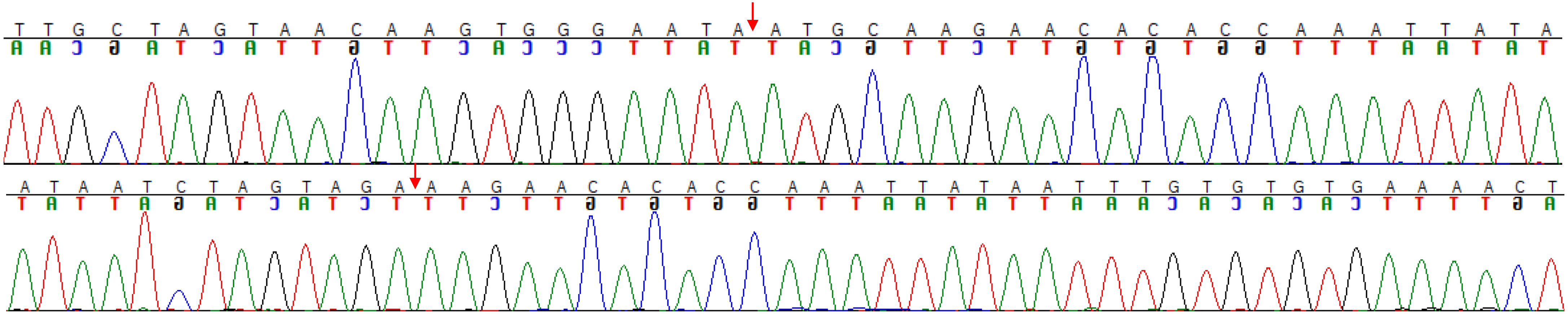

Line 4, *SlDMR6* biallelic, *SlINVINH1* biallelic

ACCCGAATCCGATAGACCAC**GTCTATCGGAAGTGGTCGATTGT**GAAAATGTTCCAATAATTGACTTAAGTTGCGGAGATCAAGCT SlDMR6 WT
ACCCGAATCCGATAGACCAC**G-------------GTCGATTGT**GAAAATGTTCCAATAATTGACTTAAGTTGCGGAGATCAAGCT line 4, allele 1, -13bp
ACCCGAATCCGATAGACCA-**--------GAAGTGGTCGATTGT**GAAAATGTTCCAATAATTGACTTAAGTTGCGGAGATCAAGCT line 4, allele 2, -9bp

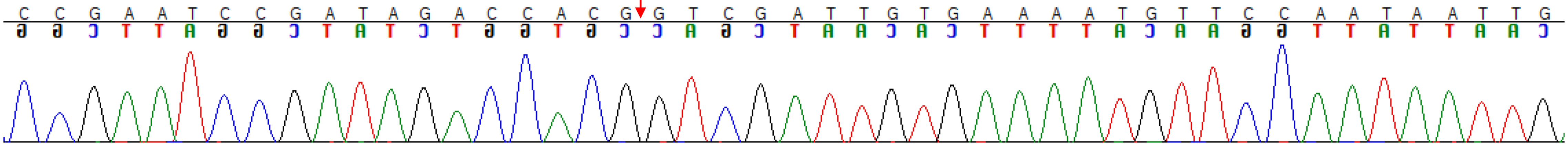


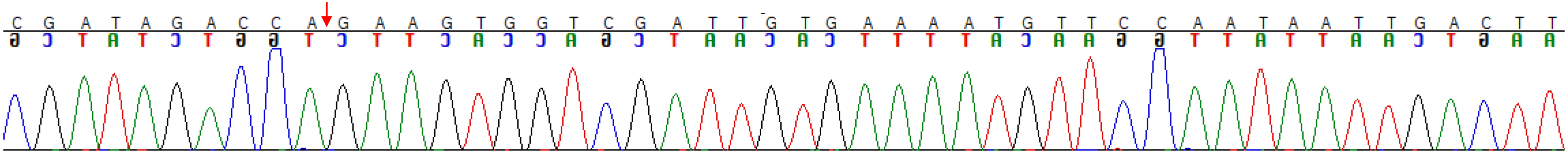


TAGTAACAAGTGGGAATAATAATCTAGT**AGAGACAACATGCAAGAACACAC**CAAATTATAATTTGTGTGTGAAAACTTTGTCT *SlINVINH1* WT
TAGTAACAAGTGGGAATAATAATCTAGT**A--------ATGCAAGAACACAC**CAAATTATAATTTGTGTGTGAAAACTTTGTCT line 4,allele 1,-8bp
TAGTAACAAGTGGGAATAATAATCTA--**---------ATGCAAGAACACAC**CAAATTATAATTTGTGTGTGAAAACTTTGTCT line 4,allele 1,-11bp

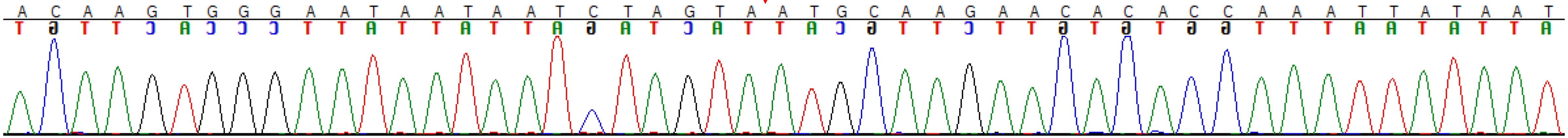


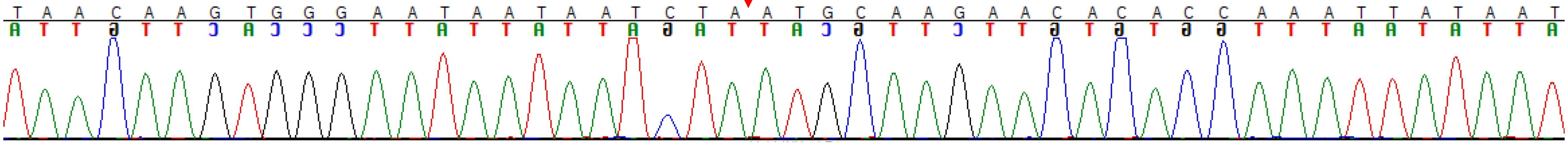

Line 6, *SlDMR6* WT, *SlINVINH1* biallelic

TAGTAACAAGTGGGAATAATAATCTAGT**AGAGACAACATGCAAGAACACAC**CAAATTATAATTTGTGTGTGAAAACTTTGTCT *SlINVINH1* WT
TAGTAACAAGTGGGAATAATAATCTAGT--------**CATGCAAGAACACAC**CAAATTATAATTTGTGTGTGAAAACTTTGTCT line 6,allele 1,-8bp
TAGTAACAAGTGGGAATAATAATCTAGT--**AGACAACATGCAAGAACACAC**CAAATTATAATTTGTGTGTGAAAACTTTGTCT line 6,allele 2,-2bp

*slinvinh1*, line 6, Allele 1

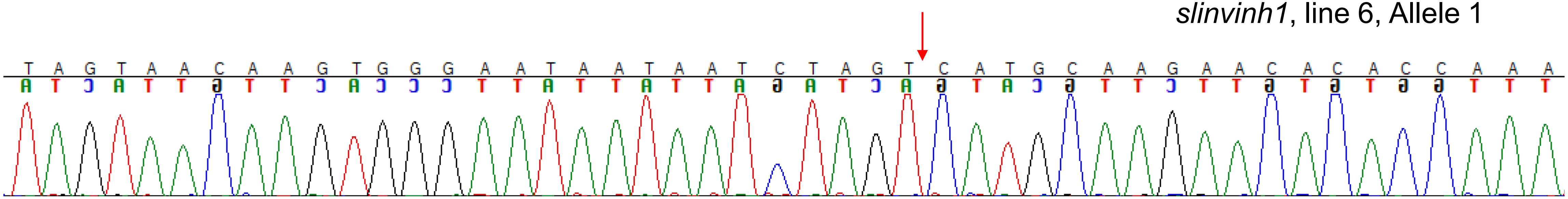


*slinvinh1,* line 6, Allele 2

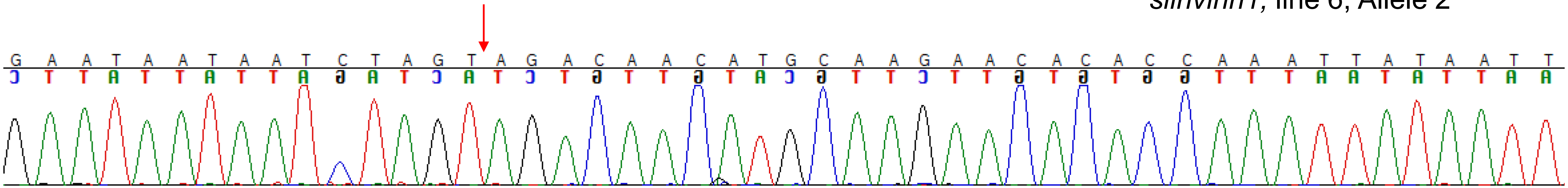

Line 12, *SlDMR6* WT, *SlINVINH1* biallelic

TAGTAACAAGTGGGAATAATAATCTAGT**AGAGACAACATGCAAGAACACAC**CAAATTATAATTTGTGTGTGAAAACTTTGTCT *SlINVINH1* WT
TAGTAACAAGTGGGAATAATAATCTAGT**AGA-------TGCAAGAACACAC**CAAATTATAATTTGTGTGTGAAAACTTTGTCT line 12,allele 1,-7bp
TAGTAACAAGTGGGAATAATAATCTAGT**AGAGACA---TGCAAGAACACAC**CAAATTATAATTTGTGTGTGAAAACTTTGTCT line 12,allele 2,-3bp

*slinvinh1*, line 12, Allele 1

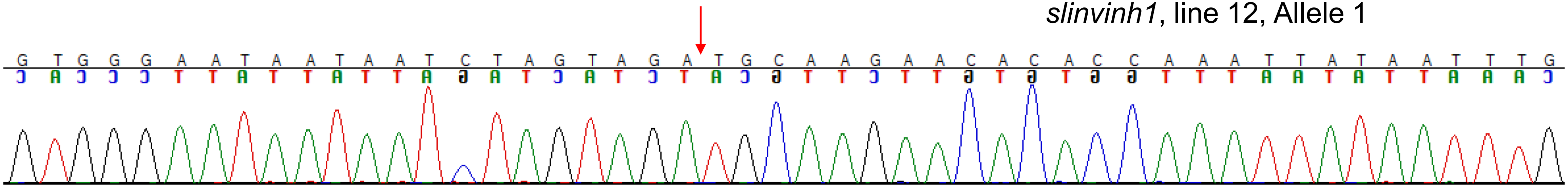


*slinvinh1*, line 12, Allele 2

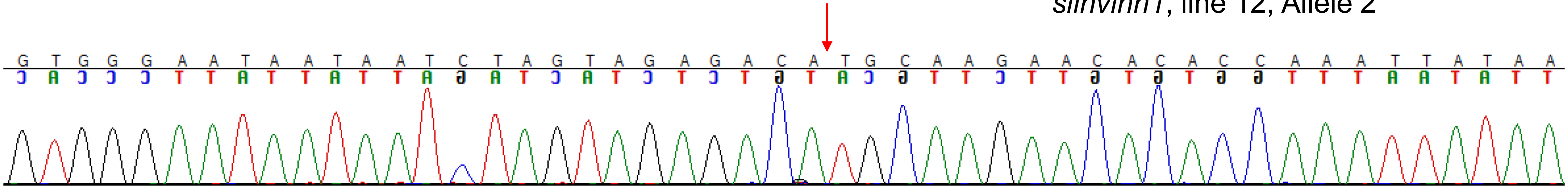

Line 15, *SlDMR6* biallelic, *SlINVINH1* biallelic

ACCCGAATCCGATAGACCAC**GTCTATCGGAAGTGGTCGATTGT**GAAAATGTTCCAATAATTGACTTAAGTTGCGGAGATCAAGCT SlDMR6 WT
ACCCGAATCCGATAGACCAC**GT-------AAGTGGTCGATTGT**GAAAATGTTCCAATAATTGACTTAAGTTGCGGAGATCAAGCT line 15, allele 1, -7bp
ACCCGAATCCGATAGACCA-**----------AGTGGTCGATTGT**GAAAATGTTCCAATAATTGACTTAAGTTGCGGAGATCAAGCT line 15, allele 2, -11bp

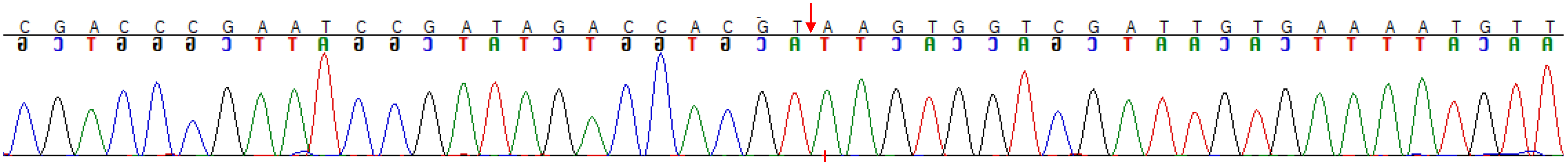

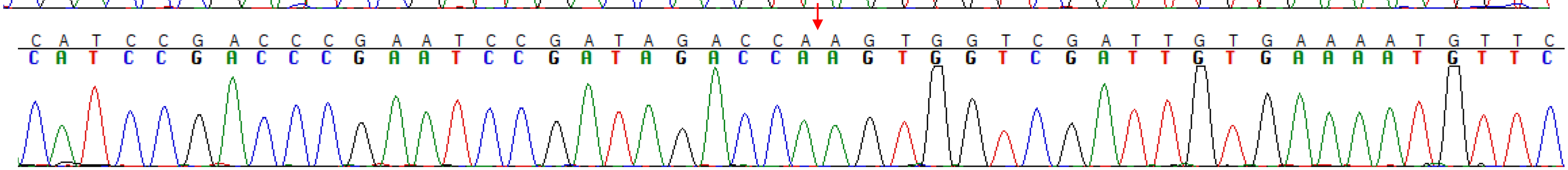

TAGTAACAAGTGGGAATAATAATCTAGT**AGAGACAACATGCAAGAACACAC**CAAATTATAATTTGTGTGTGAAAACTTTGTCT *SlINVINH1* WT
TAGTAACAAGTGGGAATAATAATCTAGT**-------ACATGCAAGAACACAC**CAAATTATAATTTGTGTGTGAAAACTTTGTCT line 15,allele 1,-7bp
TAGTAACAAGTGGGAATAATAATCTAGT**AGAGACA---TGCAAGAACACAC**CAAATTATAATTTGTGTGTGAAAACTTTGTCT line 15,allele 2,-3bp

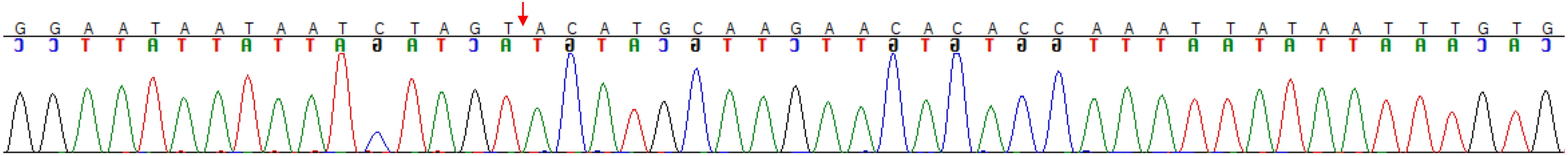

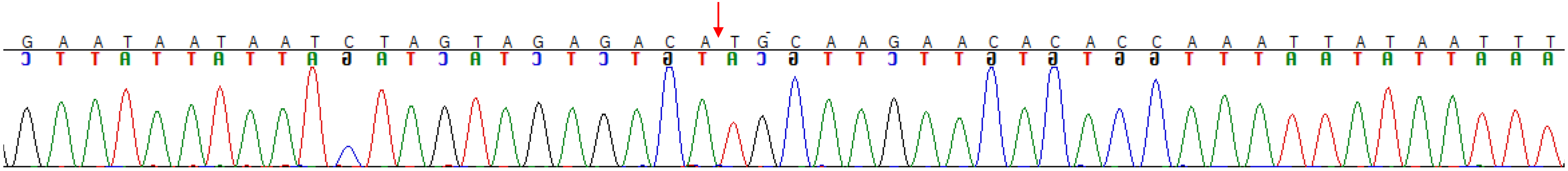

Line 16, *SlDMR6* heterozygous, *SlINVINH1* homozygous

```
ACCCGAATCCGATAGACCACGTCTATCGGAAGTGGTCGATTGTGAAAATGTTCCAATAATTGACTTAAGTTGCGGAGATCAAGCT SlDMR6 WT
ACCCGAATCCGATAGACCACGTCTATCGGAAGTGGTCGATTGTGAAAATGTTCCAATAATTGACTTAAGTTGCGGAGATCAAGCT line 16, allele 1, WT
ACCCGAATCCGATAGACCA-----------AGTGGTCGATTGTGAAAATGTTCCAATAATTGACTTAAGTTGCGGAGATCAAGCT line 16, allele 2, -11bp
```

*sldrm6* , line 16, Allele 2

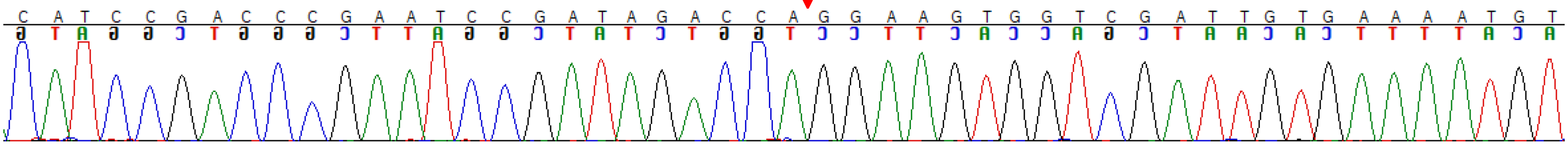


```
TAGTAACAAGTGGGAATAATAATCTAGTAGAGACAACATGCAAGAACACACCAAATTATAATTTGTGTGTGAAAACTTTGTCT SlINVINH1 WT
TAGTAACAAGTGGGAATAAT--------------AACATGCAAGAACACACCAAATTATAATTTGTGTGTGAAAACTTTGTCT line 16, -14bp, homozygous
```

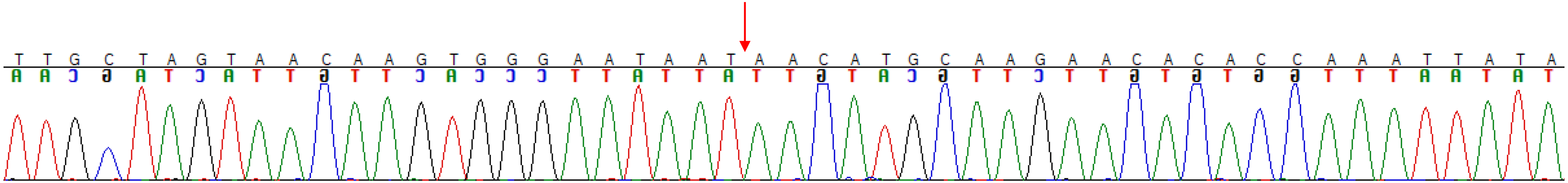

Line 17, *SlDMR6* WT, *SlINVINH1* biallelic

TAGTAACAAGTGGGAATAATAATCTAGT**AGAGACAACATGCAAGAACACAC**CAAATTATAATTTGTGTGTGAAAACTTTGTCT *SlINVINH1* WT
TAGTAACAAGTGGGAA-----------------**CAACATGCAAGAACACAC**CAAATTATAATTTGTGTGTGAAAACTTTGTCT line 17, -17bp
TAGTAACAAGTGGGAATAATAATCTAGT----------**TGCAAGAACACAC**CAAATTATAATTTGTGTGTGAAAACTTTGTCT line 17, -10bp

*slinvinh1* , line 17, Allele 1

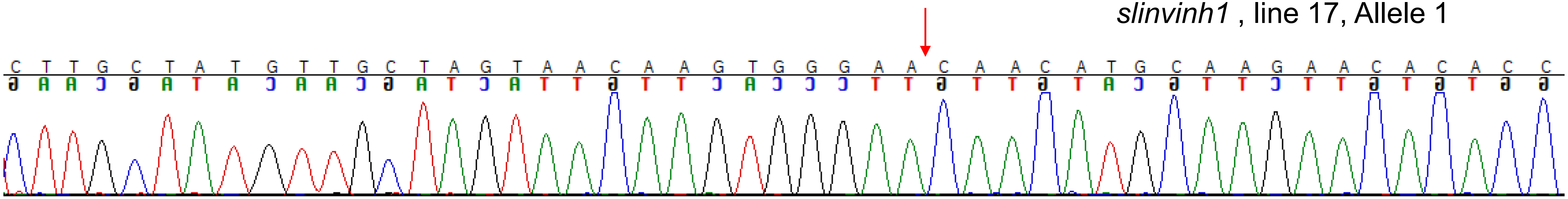


*slinvinh1* , line 17, Allele 2

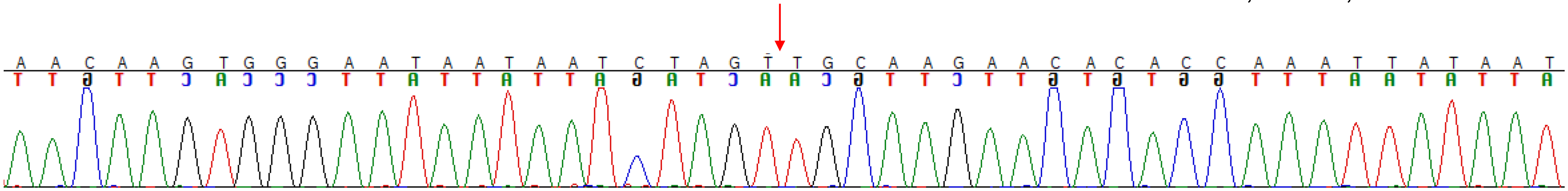
